# Engineering of highly active and diverse nuclease enzymes by combining machine learning and ultra-high-throughput screening

**DOI:** 10.1101/2024.03.21.585615

**Authors:** Neil Thomas, David Belanger, Chenling Xu, Hanson Lee, Kathleen Hirano, Kosuke Iwai, Vanja Polic, Kendra D Nyberg, Kevin G Hoff, Lucas Frenz, Charlie A Emrich, Jun W Kim, Mariya Chavarha, Abi Ramanan, Jeremy J Agresti, Lucy J Colwell

## Abstract

Optimizing enzymes to function in novel chemical environments is a central goal of synthetic biology, but optimization is often hindered by a rugged, expansive protein search space and costly experiments. In this work, we present TeleProt, an ML framework that blends evolutionary and experimental data to design diverse protein variant libraries, and employ it to improve the catalytic activity of a nuclease enzyme that degrades biofilms that accumulate on chronic wounds. After multiple rounds of high-throughput experiments using both TeleProt and standard directed evolution (DE) approaches in parallel, we find that our approach found a significantly better top-performing enzyme variant than DE, had a better hit rate at finding diverse, high-activity variants, and was even able to design a high-performance initial library using no prior experimental data. We have released a dataset of 55K nuclease variants, one of the most extensive genotype-phenotype enzyme activity landscapes to date, to drive further progress in ML-guided design.

## Introduction

The ability to engineer proteins has revolutionized applications in industry and therapeutics^1–6^. Generally, a protein engineering campaign can be divided into two stages^7–9^: *discovery* first finds a candidate protein that performs a desired function at a non-zero level of activity and then *optimization* improves its attributes, such as the binding strength of an antibody ^10^ or the catalytic activity^11^, thermostability^12,13^, or stereoselectivity of an enzyme^14^. *Directed evolution* (DE) is the standard technique for protein optimization, where a pool of genotypes is iteratively improved using *in-vitro* selection and mutagenesis^15–18^. Proteins optimized using DE have been deployed in many industrial and therapeutic settings^19^, however the technique can fail to cross valleys in the protein fitness landscape and may converge prematurely when the diversity of the pool collapses^20^.

To address these limitations, ML-guided directed evolution (MLDE), involving rounds of data collection, modeling, and model-guided sequence generation, has emerged as a potential alternative to DE^9,21–24^. However, there is limited work directly comparing the techniques across multiple rounds of high-throughput experiments, an increasingly-common setting due to advancements in automation^25^, microfluidics^26,27^, display systems^27–30^, multiplexing^31^, and continuous evolution^32,33^. Instead, many MLDE vs. DE case studies have occurred in regimes with such low throughput that a standard DE workflow based on random mutagenesis would be infeasible^34–39^, have only performed a single round of experiments^40,41^, or have sought to maintain a target phenotype instead of improving it^42^. Comparison in the high-throughput regime is important because both techniques benefit from increased population size.

This paper presents TeleProt (Figure 1), an MLDE framework that balances evolutionary data from natural homologs with protein fitness data accumulated across rounds of experiments. We compare our MLDE approaches head-to-head with DE, using the same ultra-high-throughput microfluidics platform, to engineer the endonuclease NucB to be active at pH 7.

**Figure 1:**
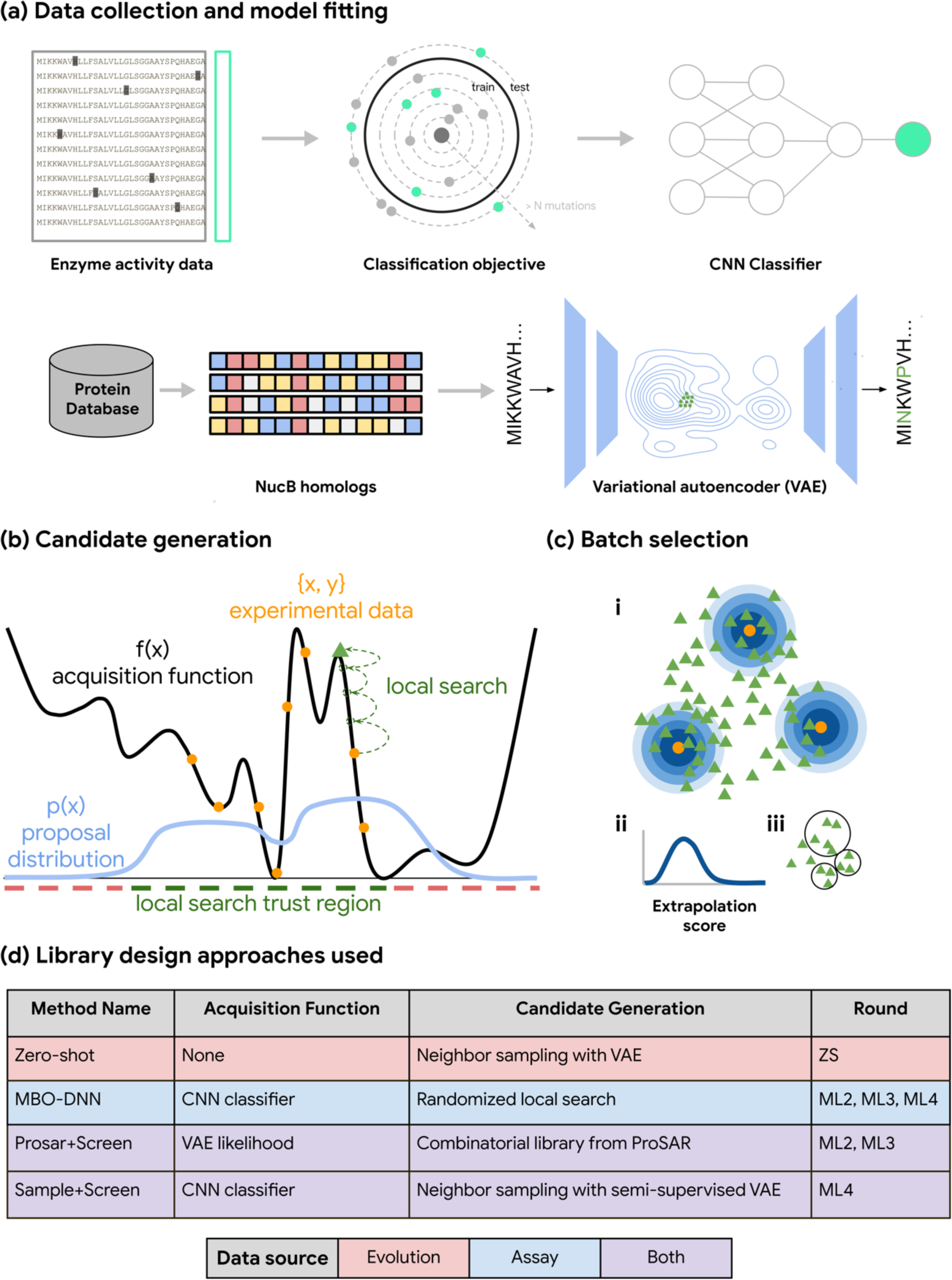
Our model-guided sequence design framework (TeleProt). (a) Models are trained on natural NucB homologs, data from prior rounds of experiments, or a combination of both. (b) Candidate generation seeks to find novel variants with high acquisition function scores. Generations are guided by either a “proposal distribution” (light blue) or a randomized “local search” (green). To avoid discovering implausible, yet high-scoring candidates, the search is constrained to remain within a trust region (green). (c.i) Candidates (green triangles) are generated using a proposal distribution, or local search from known hits (orange). Every candidate is assigned an “extrapolation score” (blue), which in ML4 is the hamming distance from a known hit. (c.ii) We trade off exploration and exploitation by imposing a target distribution of extrapolation scores. (c.iii) A final batch of diverse candidates is selected using a greedy, approximate algorithm to both satisfy this target score distribution and to limit the usage of each possible mutation. (d) In each round of our campaign (Figure 3a), we employed an ensemble of multiple MLDE methods with different exploration-exploitation tradeoffs and levels of dependence on evolutionary vs. experimental data.

NucB is secreted naturally by *Bacillus licheniformis*^43^ and can degrade the extracellular DNA required for the formation of biofilms^44^. A challenge limiting NucB application to chronic wound care^45–47^ and anti-biofouling^48^ is that enzyme activity drops ∼80% at pH 7, compared to its native alkaline pH (Supplementary Text 6C). Therefore, the goal of our protein engineering campaign was to not only restore but improve NucB’s nuclease activity at pH 7, a necessary prerequisite for its application as a wound healing therapeutic at physiological pH^49^.

We show that our MLDE methods deliver significant benefits compared to DE in both hit rate and diversity, and identify high-activity NucB variants that are promising candidates for biofilm degradation. We also demonstrate that an ML-guided zero-shot design technique can generate better initial libraries than error-prone PCR. In the process, ablation studies, where we change a single aspect of a TeleProt method, provide general guidelines for how to use TeleProt going forward. All experimental data from the project and code reproducing our analyses is available at https://github.com/google-deepmind/nuclease_design.

## Results

Our ML-guided campaign (**ML**) employed a portfolio of different instances of the TeleProt framework (Figure 1 and Methods) for blending evolutionary and experimental data. Each approach used the same general approach. First, a model was fit on natural NucB homologs, experimental data, or both (Figure 1a). This provided either an acquisition function, an in-silico predictor of enzyme quality, or a sampling distribution for novel NucB variants. Then, a large set of high-scoring candidate variants was identified, using safeguards to avoid variants outside the distribution of experimental and evolutionary data, where the models may perform unreliably (Figure 1b). Finally, a diverse subset of candidates with a desired distribution of levels of extrapolation from the training data was selected for screening (Figure 1c). Within each round, we divided our design budget across our portfolio to increase library diversity and to investigate algorithmic choices.

An ultra-high-throughput microfluidic platform was used to measure the catalytic activity of large libraries of NucB variants at pH 7 simultaneously (Figure 2a). Variants were transformed into *B. subtilis* and individually encapsulated into droplets. After growth and protein expression, the nuclease substrate, a 20-mer dsDNA carrying a dye and a quencher, was injected into the droplets. Cleavage by the nuclease causes the dye and the quencher to separate, increasing the fluorescence of the droplet (see Supplementary Figure S27). Since nuclease activity is tied to fluorescence, droplets can be sorted (Figure 2a) to (i) sequester the best-performing variants (Figure 2b) or to (ii) create genotype-phenotype datasets of amino acid sequences and corresponding functional activity (Figure 2c). Below, we further purify and validate top hits using lower-throughput assays.

**Figure 2:**
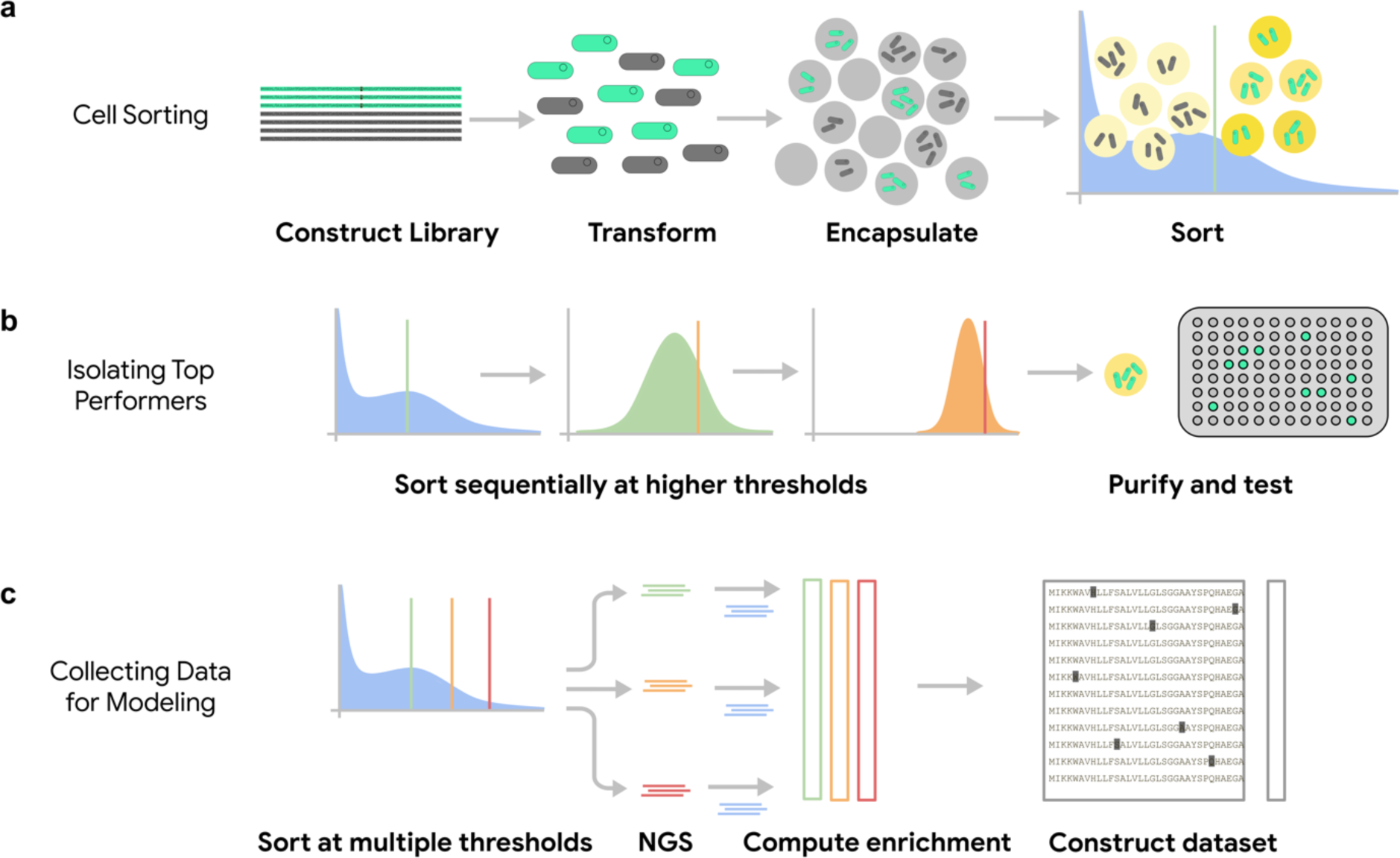
Experimental workflows for isolating high-activity variants and creating an enzyme activity dataset. (a) Selecting variants with high enzyme activity using a fluorescence-based ultra-high-throughput platform. Cells are encapsulated in droplets as a water-in-oil emulsion and injected with reagents to achieve the desired reaction conditions. (b) Repeatedly sorting a library at increasing activity thresholds applies selective pressure to identify top-performing variants. (c) Sorting at multiple thresholds in parallel produces enrichment factors that can be used to derive a dataset of genotype-activity pairs.

We ran three additional campaigns to provide qualitatively different baselines (Figure 3a). The independently-run directed evolution campaign (**DE**) involved two generations of hit selection and *in-vitro* mutagenesis using the same ultra-high-throughput platform (see Methods and Supplementary Figure S18). To enable finer-grained assessment of the overall composition of the **ML** library, we also ran a hit recombination (**HR**) campaign that was pooled with **ML** for all experiments. **HR** variants were designed *in-silico* to simulate *in-vitro* hit stacking, where novel recombinations of the top-ranked variants from the previous-round’s dataset are sampled. Finally, we designed a zero-shot library (**ZS**) using a model of natural NucB homologs to investigate whether NucB activity could be improved using no prior experimental data.

**Figure 3:**
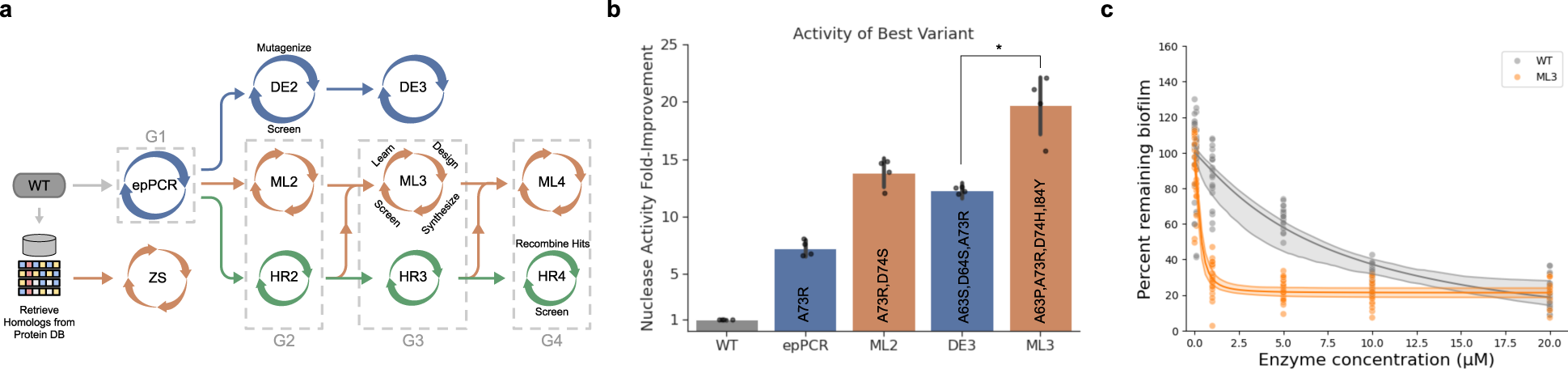
Machine learning paired with ultra-high-throughput screening designed a higher-activity nuclease enzyme than directed evolution using the same screening platform. (a) We ran parallel enzyme optimization campaigns using traditional directed evolution (DE, blue), ML-guided design (ML, orange), and hit recombination (HR, green), where all were initialized from the same error-prone PCR variant library (epPCR, G1), in which about 11% of the variants were functional and 2% had activity higher than the wildtype. As denoted by the dashed lines, ML and HR libraries were pooled during synthesis and screening. We additionally ran a single round screen of a zero-shot designed library (ZS), designed by a model fit using homologous sequences. (b) Top-performing variants from each library were purified and assessed for nuclease activity at four enzyme concentrations, normalized by the WT activity. The best variant discovered after 2 rounds of ML guided design (ML3, orange) had significantly better activity than the best variant discovered after 2 rounds of directed evolution (DE3, blue, permutation test; p < 0.05). Error bars indicate ± 1 standard deviation, which accounts for variability across enzyme concentrations and variability in kinetic rate estimation (see Supplementary Figure S14 for details on the estimate of standard deviation). (c) Dose-response curve for biofilm degradation of the best variant discovered in ML3 (orange) and the WT (gray). The best-fit line uses the Hill equation fit using the least-squares method. Confidence bands represent 5% and 95% percentiles given by the estimated Hill equation parameters. The ML3 variant provides a clear improvement over the WT at low enzyme concentration.

The **DE**, **ML** and **HR** campaigns were all built upon the same initial library, generated by applying error-prone (epPCR) to a single-site saturation mutagenesis library of the *B. licheniformis* wildtype (WT) NucB sequence (see Supplementary Figure S28). **DE** had access to an initial dataset containing at least 211K distinct epPCR variants, while the higher DNA sequencing read depth required to estimate enzyme activity, based on estimating the enrichment of a variant in the post-sort vs. pre-sort population, meant the initial dataset used for **HR** and **ML** contained just 9.4K distinct variants (see Supplementary Figure S25). The **ZS** library did not require an initial dataset, instead employing 4.6K homologs. The **ML** campaign contained about 10K variants in each round, while **HR** libraries were about 1-2K in size, akin to a large-scale plate-based high-throughput enzyme screening experiment (Supplementary Table S3). Models for **ML** were trained on all **ML** and **HR** data from prior generations.

### Machine learning paired with ultra-high-throughput screening designed higher-activity nuclease enzymes than directed evolution

To compare the performance of **ML** to **DE**, we performed a series of downselection assays on each library to isolate the best enzyme variant and then characterized its performance (see Supplemental Figure S7). First, we sorted droplets at a sequence of successively higher thresholds (Figure 2b), and cultured selected cells on agar plates. Those colonies were then assessed for nuclease activity in liquid culture plates. Finally, top-performing strains in the liquid culture plate assay were purified and promoted to a specific activity assay. The specific activity assay measures the fold-improvement of the kinetic rate of purified enzyme by comparing the temporal fluorescence trajectories of a variant vs. the wildtype (see Methods: Nuclease activity assays).

Figure 3b shows that the best enzyme discovered by two rounds of ML-guided design (**ML3**, orange) significantly outperforms the best enzyme discovered using two rounds of DE (**DE3**, blue; permutation test, p=0.014). The best **ML2** and **ML3** variants were proposed by model-based optimization on a learned neural network landscape (MBO-DNN, see Methods). After three generations, comprising ∼23K variants, **ML3** discovered (A63P, A73R, D74H, I84Y), with a 19 fold-improved activity that outperforms the best **DE3** enzyme (A63S, D64S, A73R), with a 12 fold-improved activity at pH 7. Both exceeded our goal of restoring the activity level to that of the WT NucB at its optimal pH 9: the **ML3** enzyme showed 2.4 fold-improved activity, while the best **DE3** variant showed 1.5 fold-improved activity (see Supplementary Figure S16). Since the ML and HR libraries were pooled in each round and our stringent selection process isolated only the top variants, the fact that no **HR** proteins were selected for purification suggests that **HR** did not discover any variants with activity comparable to the best from **ML3** and **DE3**. We did not purify any variants from **DE2**, as the DE campaign was run independently with the goal of characterizing the overall highest-activity variant.

The ability of the best **ML3** variant to degrade biofilms at the relevant pH for wound healing was further confirmed in a follow-up screen. We cultured *Bacillus licheniformis,* which produces biofilm naturally, and exposed the biofilm to the enzyme at neutral pH (see Methods). In Figure 3c, we show that the variant degraded these biofilms more effectively than the wildtype enzyme across a variety of doses. At 1-µM enzyme concentration, it degraded 71 ± 2% of the biofilm compared to 13 ± 3% degraded by the WT (t-test, p < 1e-15).

### Machine learning discovered hundreds of diverse hits with high activity

Over three rounds of our campaign, we achieved two key goals: we improved enzyme activity at pH 7 beyond the WT activity at pH 9 and we demonstrated that **ML** did so better than **DE**. To further investigate the capabilities of ML-guided campaigns, we developed a fourth round of designs (**ML4**) with the goal of finding *diverse*, high-activity hits. Maintaining additional diversity over directed evolution is one of the key promises of ML-guided design^50^. Most design campaigns require optimizing over multiple objectives, such as expression, toxicity, and stability. Increasing the diversity of designs without sacrificing performance can increase the chance that a variant meets the requirements for additional properties in downstream screens. Evaluating **ML** performance in terms of diversity relies on characterizing enzyme activity for a large sample of the variants in parallel, and we enabled a head-to-head comparison against **HR**, a non-ML baseline, by pooling the **ML** and **HR** designs in each round (Figure 3a). We compare designed libraries in terms of both the ‘hit-rate’, the proportion of variants with activity above a given activity threshold, and the ‘diversity’, measured as the number of clusters of hits when clustered at a given sequence identity threshold.

To characterize hits at high-throughput, we sequence both the pre-sort and post-sort libraries (Figure 2c), compute an enrichment factor for each variant, and compare this with the empirical distribution of enrichment factors for synonymous variants of a *fiducial* reference enzyme with known activity (see Methods). A variant is labeled as more active than the fiducial using a right-sided t-test, and we estimate the hit rate of a collection of variants using a Benjamini-Hochberg^51^ correction with an expected false discovery rate of 0.1 to account for multiple testing. Hit rates for all library design methods in all generations are provided in Supplementary Table S1. We refer to the pooled experiments containing both **ML** and **HR** libraries, as *generations* G1 to G4 (Figure 3a), where G1 is the initial epPCR library. When analyzing the methods’ performance in G4, we consider hit rates with respect to three fiducials of increasing activity: WT, A73R, and A73R,D74S. A73R was the top-performing variant from the initial epPCR library, with 8-fold higher activity than the WT at pH 7 (1.3-fold higher activity than the WT at pH 9). A73R,D74S was the best mutant seen in G2, with 13-fold improved activity (Figure 3b).

**ML4-MBO-DNN** (Methods) was the strongest-performing member of the portfolio of methods in **ML4** in terms of hit rate (see Supplementary Table S1). It had similar size as **HR4** (1356 vs. 1540 variants) and significantly outperformed **HR4** at finding hits above the A73R level (chi-square test; P=1.3×10^-9^): 52 ± 7 of the 1356 **ML4-MBO-DNN** variants (hit-rate 3.9% ± 0.5%; standard deviations estimated via bootstrapping) had greater activity than A73R, compared to only 7 ± 4 of the 1540 **HR4** variants (hit-rate 0.5% ± 0.2%). For discovering variants with activity greater than the WT, **ML4-MBO-DNN** significantly outperformed **HR4** (chi-square test; p < 1e-15). **ML4-MBO-DNN** found 1145 ± 15 hits out of 1356 designs (hit-rate 84.5% ± 1.1%), while **HR4** found 890 ± 17 hits out of 1540 designs (hit-rate 58.0% ± 0.1%). Finally, **ML4** discovered 31 variants with significantly better activity than A73R,D74S, of which 29 were designed by our MBO-DNN technique (2.2% ± 0.5% hit rate) and 21 had > 9 mutations (Supplementary Table S2). In contrast, **HR4** only discovered two variants better than the best variant in ML2 (0.1 ± 0.1 hit rate) and these each had just 4 mutations.

To quantify diversity for each library, we cluster hits by sequence similarity and count the number of clusters (see Methods). By sweeping over thresholds for the cluster diameter, we found that the **ML4-MBO-DNN** hits were more diverse than those in **HR4**, both for variants with activity higher than A73R (Figure 4b) and for variants with activity higher than WT (see Supplementary Figure 16). While all 5 **HR4** hits are within 7 mutations of one another, the 52 **ML4-MBO-DNN** hits maintain 10 distinct clusters even when clustering at Hamming distance 10 (cluster diameter = 10, Figure 4b). We also see that there is substantially more diversity in the pre-sort pool (Figure 4b, dotted lines) for **ML4-MBO-DNN** than for **HR4**, though the libraries are approximately the same size with ∼10^3^ variants. Furthermore, compared to **HR4** hits, which focused on 10 positions, **ML4-MBO-DNN** hits placed mutations across 38 positions in the protein, encompassing the DNA interface, all secondary structure domains, and unstructured loops (Figure 4d).

**Figure 4:**
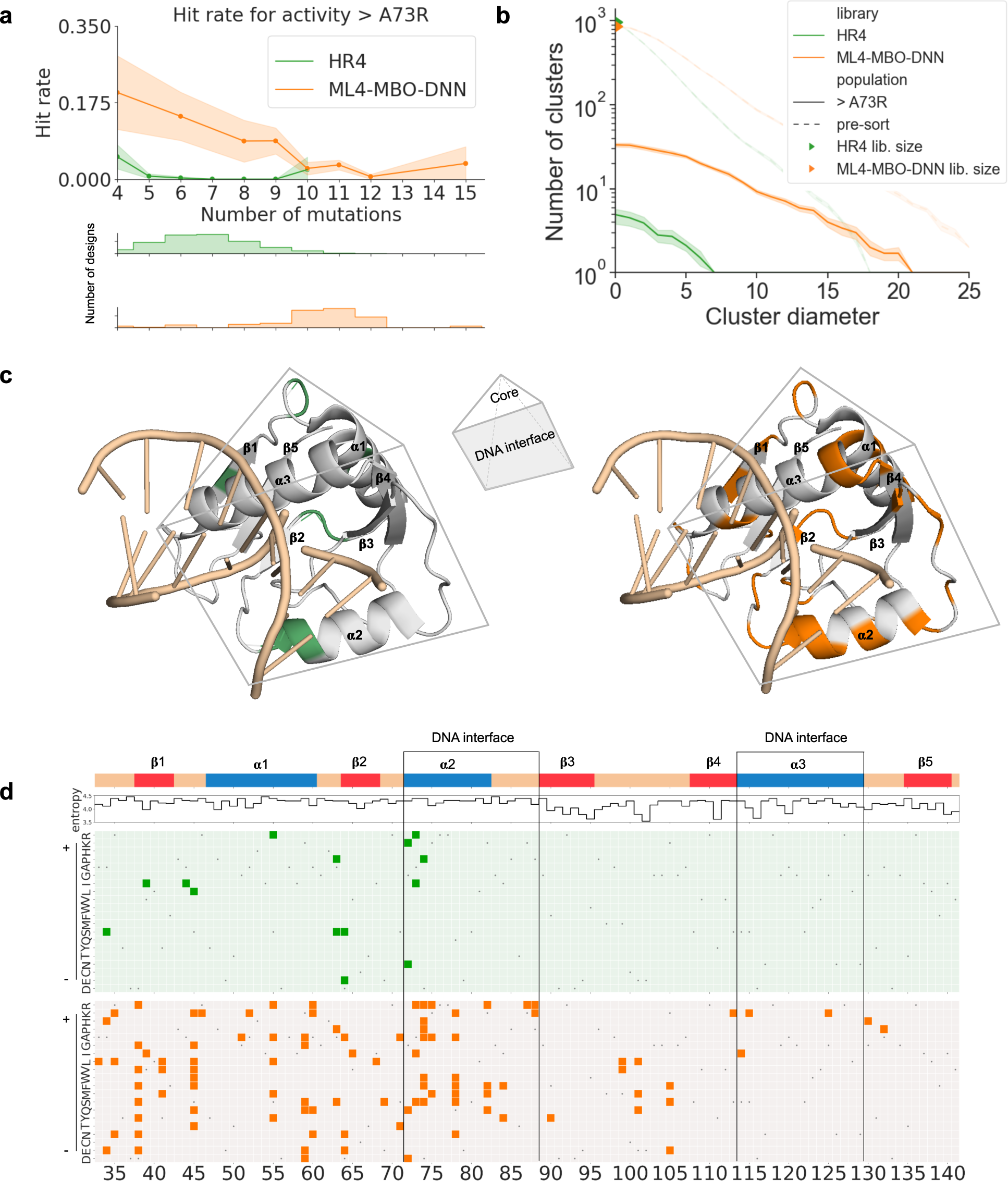
Pooled screening of ML-designed libraries alongside non-ML libraries reveals the significance of our method’s improvement to hit rates and diversity. (a) In G4, our MBO-DNN sequence design technique (ML4-MBO-DNN, orange) has a far higher hit rate than hit-recombination (HR4, green) for improving activity above the level of A73R, the best variant observed in the epPCR data. Notably, we are able to extend designs out to 15 mutations from the NucB wildtype, while still exceeding this threshold. (b) Clustering the hits by sequence similarity reveals that active ML4-MBO-DNN designs (solid orange) were substantially more diverse than active HR4 designs (solid green). The diversity of active designs was driven by both the improved hit-rate of ML4-MBO-DNN compared to HR4 and the diversity of the designed library prior to sorting (dotted lines). The cluster diameter is defined as the maximum Hamming distance between two sequences in the same cluster. (c) Three-dimensional model of NucB (PDB ID: 5OMT) with DNA based on the structural homology with Vvn endonuclease (PDB ID: 1OUP) as described in Baslè, et al.^105^. NucB forms a square pyramid structure with a positively charged DNA binding interface and a negatively charged core. Mutations found in the HR4 and ML4-MBO-DNN designs are highlighted on the structure in green (left) and orange (right), respectively. (d) NucB secondary structure, per-position Shannon entropy, and mutations identified by the HR (green) and ML (orange) campaigns. The 52 highly-active nucleases designed by ML exhibited mutations that span more diverse positions (orange) in the amino acid sequence than 9 HR hits (green). Amino acids are ordered by increasing isoelectric point (pI). Wildtype residues are indicated in gray.

### Machine learning designs revealed the ability of models to extrapolate beyond their training set

In typical directed evolution campaigns, a small number of mutations are incorporated into the protein in each generation^16^. For **HR4**, the distribution of distances from WT is dictated by the top **HR3** hits, while for **ML**, the distribution of distances is an algorithmic choice. To investigate whether our ML techniques can more rapidly traverse the fitness landscape while maintaining or improving activity, we deliberately designed variants that were many mutations beyond the highest activity variants seen in the training data. Because we wanted to test the ability to diversify the target protein, we define hits as having activity greater than WT. In Figure 5a we find that many of these **ML** hits had many more mutations than the hits from prior generations used to train the models. Further, the hits were not simple recombinations of mutations known to be active, but included mutations that had not previously been seen in a variant of the desired activity level (Figure 5b). Prior to G4, no variant with activity greater than WT and > 9 mutations was observed (Figure 5a), and no variant with activity greater than A73R with > 7 mutations was observed (not shown). Despite this, **ML4** discovered variants with greater activity than A73R with up to 15 mutations, while simultaneously improving over the best activity seen in **ML3**. We also validated two **ML4** hits containing 11 mutations with greater activity than A73R in the liquid culture plate assay (see Supplementary Figure S13).

**Figure 5:**
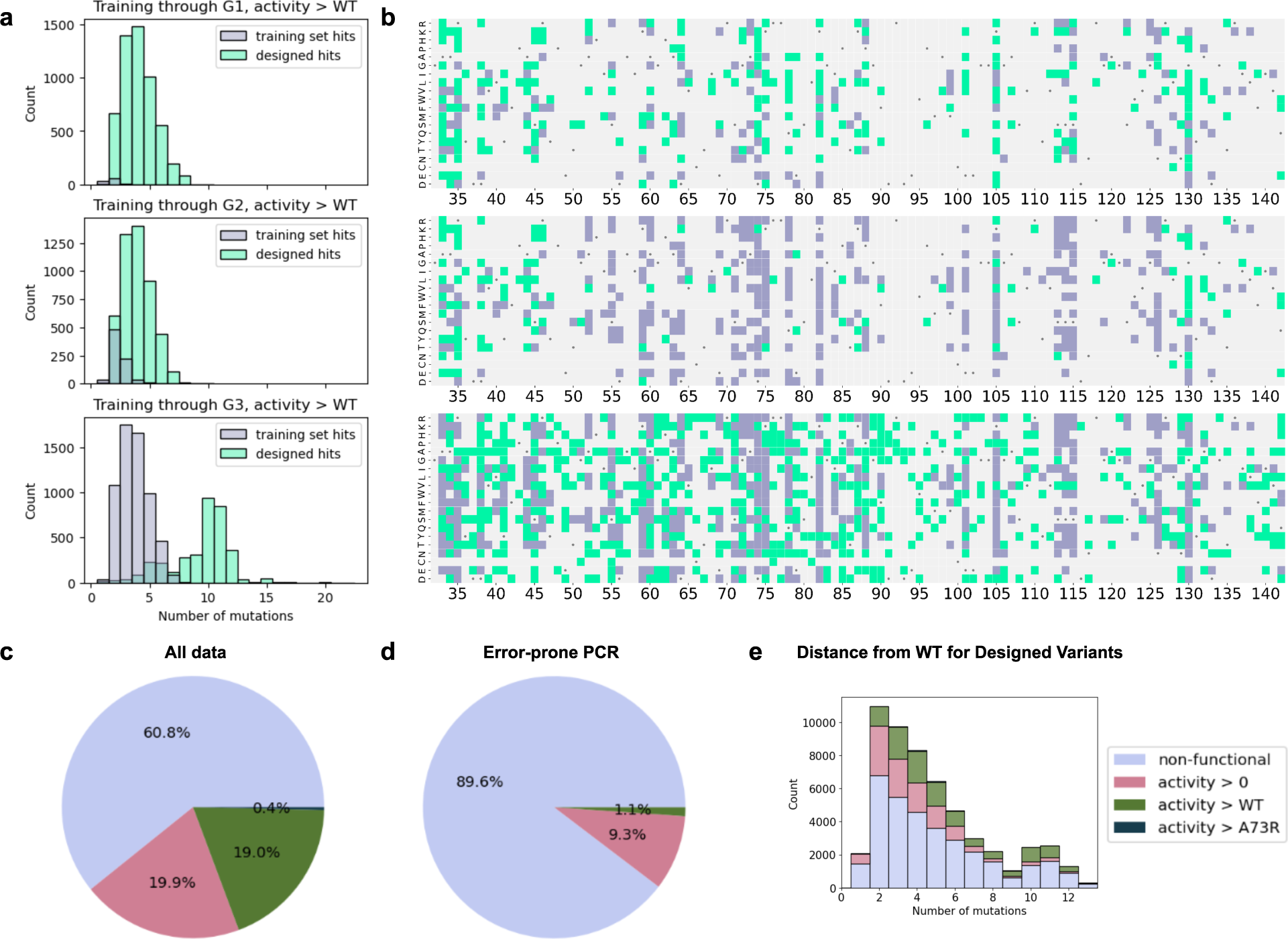
(top) Models designed functional variants are qualitatively different from the models’ training data. (a) Model-designed variants (green) were active at distances from the wildtype well beyond active variants seen in the training set (purple). In (b) we visualize the distinct mutations present in the hits in the training set and designs. The set of mutations found in active variants included novel mutations extending beyond those seen in training data from prior rounds. By definition, hit recombination is constrained to recombine hits seen in the prior rounds and thus it can not expand the set of mutations beyond the purple mutations in any round. Each row corresponds to models trained on data from a given round, i.e. the first row shows designs from models trained on G1 data (including the ‘g2_redux’ library screened in parallel with G4), the second row shows designs from models trained on G2 data, and the third row shows designs from models trained on G3 data. **(bottom) Overview of the genotype-phenotype landscape.** (c) The overall dataset, consisting of designed variants and error-prone PCR variants, provides many representatives across a broad range of activity levels. (d) The error-prone PCR dataset is more imbalanced. (e) The overall dataset contains many functional variants and variants with activity > WT that are far from the WT (x-axis clipped to show a maximum of 13 mutations).

We hypothesized that the extrapolation capabilities of our ML methods depended on models’ ability to infer epistasis, or non-additive effects of multiple mutations^52^. Epistasis can be learned from non-additive interactions observed in the experimental data^20,53,54^, from correlations between residues in related natural homologs^55–58^, or from combining both^59,60^. To better understand which components were important for such extrapolation, we paired our ML-guided libraries with ‘ablation’ sub-libraries that used the same overall design strategy but relied on an additive model (see Supplementary Text S2). In G3, we generated variants using the ProSAR method^61^, which assumes additive mutation effects, and an extension where ProSAR variants were filtered using scores from a non-additive model of nucB homologs. Doing so significantly increased the hit rate for discovering variants with activity higher than A73R from 4.5% ± 2.3% to 23.2% ± 6.2% (see ‘g3_prosar_low_unscreened’ and ‘g3_prosar_high_screen_high’ Supplementary Table S1, Supplementary Figure S3). This suggests that information from the homologs helped remove non-functional recombination variants, perhaps due to the model’s non-additivity. Inspired by this result, in G4 we tested whether this semi-supervised learning approach would have added additional value even earlier in the campaign, when experimental data was limited. Our ‘g2 redux’ library only employed G1 data and had a similar hit rate for discovering functional variants as HR2, while incorporating about twice as many mutations per variant (Supplementary Figure S3). Additional ablations (Supplementary Text S2) investigate the impact of algorithmic choices, such as the algorithm for generating candidates with high model score in MBO-DNN (Supplementary Figure S4).

### Zero-shot models improved enzyme activity using no prior experimental data

In a protein optimization campaign, the first round of screening is essential for finding promising directions for further evolution of the target. Typically, the initial library is created using either random mutagenesis (e.g., epPCR) or single-site mutagenesis (SSM). These techniques often lead to a library either with low diversity or containing many non-functional variants, both of which can hinder subsequent optimization rounds based either on ML or directed evolution^62^. To investigate whether we could have generated a better initial library than our G1 library (created using a combination of epPCR and SSM) using ML-based ‘zero-shot’ sampling (Figure 1), we ran an additional experiment. We designed variants without using any existing experimental data, instead relying on the conservation patterns observed in natural NucB homologs. We employed a variational autoencoder with fully-connected encoder and decoder based on the DeepSequence model^63^, which was fit on a multiple sequence alignment and can both score the likelihood of a given sequence or sample novel sequences.

We decided to investigate zero-shot design after initial analysis on the epPCR data (Figure 6a) demonstrated that sequences with low likelihood under the VAE tended to be non-functional, as observed in prior work for a variety of proteins and models^59,63–69^ (Figure 6a). For example, filtering the set of multi-mutant epPCR variants to remove the bottom-scoring 50% would have removed only about 10% of the functional variants (Figure 6b). In Supplementary Text S3, we further show that our model outperforms other popular mutation scoring models including ESM-1b^70^ and a profile hidden Markov model^71^ at identifying functional variants and better-than-WT variants. Here and below, variants are labeled as functional using the enrichment factors from variants containing premature stop codons to define a null distribution. Note that typically, zero-shot design approaches would not be able to validate the model on experimental data before sampling novel variants. However, our decision to allocate a portion of G4 to zero-shot samples was informed by model performance on scoring G1 variants. Our approach is still zero-shot because we did not tune any VAE parameters or hyper-parameters on experimental NucB data.

**Figure 6:**
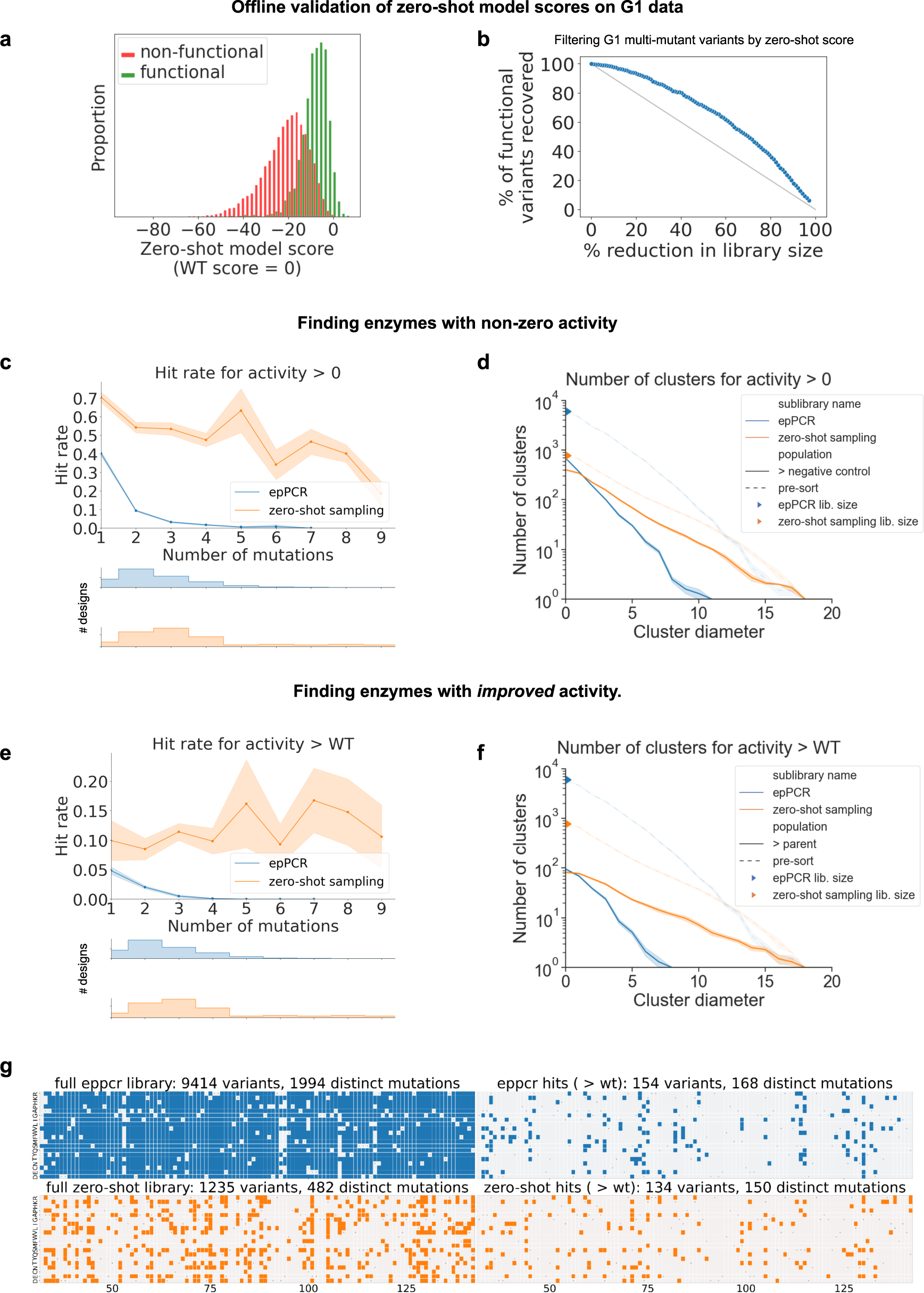
Without any experimental screening data, sampling from a zero-shot model of natural homologs is significantly better than *error-prone PCR (epPCR)* for providing an initial set of diverse hits that maintain and improve NucB function. (a) We initially validated our zero-shot model by observing that sorting the epPCR library by model score provides significant separation between non-functional and functional variants. See Supplementary Text S3 for comparisons to other zero-shot models. (b) Removing the bottom 50% of epPCR by score would only remove about 10% of the functional variants; the gray line represents the performance of random sub-sampling. (c) A significantly higher fraction of the novel variants sampled from the zero-shot model have non-zero activity than epPCR, despite incorporating more mutations per variant and (d) having higher sequence diversity. (e) The sampled library does not simply incorporate neutral mutations; the zero-shot designs also exhibit a much better hit rate for improving activity above the wildtype and (f) these improved hits are diverse. (g) While the zero-shot library was much smaller than epPCR, both in terms of the number of variants and the number of distinct mutations explored, it was able to discover a comparable number of promising mutations appearing in at least one variant with better-than-wildtype activity. The WT residue at each position is indicated by a gray dot.

We next experimentally characterized the activity of a library of novel variants sampled from the VAE (see Methods) and compared the quantity and diversity of hits to those generated by epPCR in G1. In Figure 6c, we show that the zero-shot technique generates functional variants with significantly better hit rate than epPCR (51.0% ± 1.8% vs. 10.9% ± 0.9%; chi-squared test, p < 1e-15), even when we sampled variants containing many more mutations from the WT than those from epPCR. We also show in Figure 6d that while the diversity of the pre-sort library for epPCR was comparable to that of zero-shot sampling, the diversity of the functional variants from zero-shot sampling was far higher. We find that the rate of negative epistasis ^52^, here defined as the fraction of non-functional double-mutation variants composed of two mutations that led to a functional variant in isolation (see Methods), was much higher in the error-prone PCR designs (49%; 193 of 392) than in the zero-shot designs (27%; 49 of 181).

In Figure 6e we further demonstrate the zero-shot library’s ability to discover variants with activity exceeding the WT. One concern when using a model fit on homologous sequences is that it would suggest mutations with neutral effect on activity, as these may tend to appear at positions with low conservation. However, we find that the zero-shot library has a significantly better hit rate than epPCR for finding variants that improve over the WT (10.9 ± 0.9% vs. 1.6% ± 0.1%; chi-squared test, p < 1e-15), even when incorporating many mutations in each variant. These hits are very diverse (Figure 6f), despite the fact that the zero-shot pre-sort library had far fewer variants than G1 (1235 vs. 9441).

Generally, follow-up optimization rounds build upon the initial library. For example, a common strategy is to propose variants that are composed of mutations that occurred in observed high-activity variants^11^. Thus, one measure of the quality of the library is the number of distinct mutations appearing in better-than-wildtype variants. In Figure 6g we find that the number of promising mutations discovered by zero–shot sampling and epPCR was about the same (168 vs. 150), despite the fact that there fewer distinct mutations appear in the zero-shot library (1994) than in epPCR (482). This suggests that zero-shot sampling could be particularly helpful in settings where experimental throughput is limited.

Finally, we further investigated whether a deep neural network is required to extract interesting candidate mutations from an alignment of natural homologs. We characterized a small library designed using a ‘homolog grafting’ technique (Methods) that incorporates mutations from the homologs in the VAE training data that are most close to the WT and found that it had a similar hit rate to zero-shot sampling but was able to generate far fewer variants (Supplementary Text S3B).

### Ultra-high-throughput screening enabled experimental characterization of the genotype-phenotype landscape

We released a 55,760-variant genotype-phenotype dataset containing all data from the **ML** and **HR** campaigns, one of the deepest-explored public enzyme fitness datasets to date. Most existing public landscapes either deeply explore combinations of few positions^20,72,73^ or explore single mutants for many positions^74,75^. Our dataset contains all single mutant variants, as well as thousands of multi-mutant combinations, many of which are active. Our dataset contains significantly more active variants (Figure 5a) than the randomly-collected epPCR data (Figure 5b), and many of the active variants have at least 10 mutations from the WT (Figure 5b). See Methods for how activity labels were derived. Since most of the variants come from the trajectories of particular optimization techniques, their properties are qualitatively different from those of randomly-collected mutants. For example, the rate of negative epistasis (i.e. double mutants AB, where both A and B result in a functional variant in isolation but AB is non-functional), is 49% (193 of 392) in the epPCR dataset but only 20% (883 of 4416) in the overall dataset, which demonstrates that our techniques effectively navigated around ‘holes’ in the landscape^62^. We hope our dataset will drive the development of ML methods that can perform even better extrapolation.

## Discussion

By combining TeleProt and an ultra-high-throughput microfluidic screening platform, we were able to discover a novel NucB variant with 19x better specific activity at pH 7 than the NucB wildtype, unlocking a key requirement for using the enzyme to degrade chronic wound biofilms at physiological pH. Analysis of our high-throughput data further suggests that we found a large number of diverse variants with activity greater than A73R, i.e. more than 8x better than the wildtype. Importantly, our ML campaign outperformed two directed evolution approaches that used the same platform: one that was run independently and using standard *in-vitro* techniques for hit selection and diversification and one that was designed *in-silico* and pooled with the ML-designed libraries for screening. Finally, the performance of our zero-shot library contributes to a growing body of evidence showing that samples from models fit on natural sequences can be used to generate libraries that are not only enriched for functional variants^76–78^ but also contain variants with *improved* fitness^41,59,79–85^.

Our results, taken as a whole, suggest a protocol for how to go about future campaigns using TeleProt. First, we suggest using zero-shot design (ZS) to generate an initial library of variants. Our results show that zero-shot design would have generated a larger, more-diverse pool of active variants for follow-up optimization rounds than DE approaches (Figure 6). Next, after the initial round of data collection, we suggest using Prosar+Screen for one or more subsequent rounds. The technique provides a simple way to balance evidence from the experimental data about promising mutations with priors about epistasis from natural homologs. Our results show that Prosar+Screen successfully designed higher quality libraries than DE approaches (Supplementary Figure S3, Supplementary Table S1). Finally, when enough data has accumulated to fit an accurate non-additive protein fitness prediction model, we suggest using MBO-DNN, which designed diverse, high activity libraries far from the WT (Figure 4).

Our TeleProt framework (Figure 1) is largely complementary to ongoing advancements in the modeling of protein function. For example, protein language models^69,70,77,86–90^, could be used in Prosar+Screen to filter multi-mutant variants or in MBO-DNN as proposal distributions or fitness predictors^53,75^. Prior knowledge derived from physics^91^, protein structure^89,92^, or experimental observations from related campaigns^93,94^ could also be used to further improve the ability of models to extrapolate to more distant sequences accurately. We hope that our large fitness landscape dataset will contribute to the development of such modeling improvements.

While the continually-decreasing costs of synthetic biology will make explicitly designed libraries more cost effective, it may also be fruitful to balance the cost of DNA synthesis with the level of control it provides. In early rounds of experiments, for example, one could employ ML to design random combinatorial libraries, which are significantly cheaper to assemble^37,95–102^. For a given budget, improvements to measurement protocols^103^ and enrichment factor estimation^104^ can also help improve the quality of datasets produced by sorting-based experiments.

TeleProt is not specific to enzymes and can be extended naturally to other proteins with available natural homologs and to molecular functions where high-throughput assays are practical. Going forward, we expect that ML methods for protein optimization will become increasingly standardized and target-agnostic. However, creating genotype-phenotype datasets will continue to require significant application-specific assay development. For example, our microfluidics approach relied on our ability to tie enzyme activity to fluorescence, which may not be straightforward for other protein functions. Overall, as advancements in synthetic biology allow us to screen a wider variety of functions with higher throughput, and computational advancements make models more data-efficient, we expect machine learning to dramatically reduce the amount of time it takes to go from an initial target to an optimized protein for many desired functions.

## STAR Methods

### The TeleProt framework for ML-guided protein optimization

#### Overview

Each round combined multiple sub-libraries designed using methods following the TeleProt framework depicted in Figure 1. TeleProt follows the structure outlined in Belanger, *et al.* 2019^106^ for model-based optimization in settings characterized by a small number of rounds, a large experimental throughput per round, and imperfect correlation between the objective function measured in the high throughput experiments and the downstream application. Here, it is important to discover a diverse set of high-performance hits to improve the chance of discovering at least one candidate satisfying downstream criteria.

Supplementary Table S1. provides a full breakdown of the library size and hit rate of each sub-library and Supplementary Text S1 provides additional implementation details.

**Search space:** We designed our libraries to include substitutions to the WT without allowing insertions or deletions. Our assembly protocol further limited our designs to contain mutations in a given 78-amino-acid design region. In G3 and G4 we employed two overlapping design regions, see ‘Handling multiple design regions.’

**Acquisition function:** Originally used in Bayesian optimization^107^, an acquisition function represents a cheap-to-evaluate software function that quantifies the utility of collecting data for a particular point in an optimization problem’s search space. Our acquisition functions are based on models fit on natural NucB homologs or experimental fitness measurements. In principle, the acquisition function could be defined jointly over an entire proposed batch, as the utility of a batch for future modeling will depend on the diversity of the batch. However, optimizing such a high-dimensional combinatorial acquisition function is very challenging. Therefore, we employ an alternative procedure for balancing the model-predicted quality of points in the batch with their diversity: we first discover a large set of points with high model score (candidate generation) and then we select a diverse subset (batch selection).

**Candidate generation:** We seek to find points with high acquisition function score while also providing safeguards that these points appear in the *trust region* of the acquisition function, the region where the model is expected to behave reliably. We use two general approaches to find such high-scoring points. Our *local search* approach, detailed in ‘Candidate generation using randomized local search,’ can be seen as an in-silico version of directed evolution. Our second approach samples candidates from a *proposal distribution* designed to place high probability on points that both appear in the trust region and have high acquisition function score.

**Batch selection:** We select candidates such that the batch has high diversity along two axes. First, we control the distribution of ‘extrapolation scores’ of variants, which acts as a heuristic for trading off exploration and exploitation of the library. Variants with high extrapolation score are in regions far from the training data, where discovering hits has high value because they would be difficult to reach using DE techniques. Doing so is difficult, however, because models may behave unreliably. Second, we select batches with a broad set of unique mutations from the WT. In practice, we found that naively taking the top-ranked candidates by acquisition function score resulted in certain mutations appearing in a large fraction of the batch. To increase the diversity of data used to train models in subsequent rounds, we manually ensure that the batch uses a broader footprint of mutations.

#### Models

We use three classes of models. Our convolutional neural network (CNN) models are fit on experimental data and are used as acquisition functions. Our variational autoencoder (VAE) models are fit on a combination of experimental and evolutionary data and can be used as either proposal distributions or acquisition functions. Our ridge regression models are fit on experimental data and are used to derive proposal distributions.

##### Convolutional neural network (CNN) activity classifier

We develop a sequence-to-function model in a standard way. First, we convert the experimental data into a set of sequences with corresponding discrete enzyme fitness labels. While enzyme activity is a continuous value appropriate for a regression model, we use discrete labels and classification models to reflect the level of resolution obtainable from our high-throughput platform. . Since we discretize at multiple activity levels, the model is a multi-class classifier. We used a CNN classification model due to prior results demonstrating the effectiveness of CNNs on fitness datasets^108^. To tune the hyperparameters of the model, we split the data into training and test sets, and pick the hyperparameters corresponding to the highest test set performance. The final model used to propose new designs was re-trained on the entire dataset using the optimal hyperparameters. All variants with 2 or fewer mutations are used for training and the remainder are reserved for the test set, which allows us to select models that can extrapolate beyond their training data to variants with more mutations. See Supplementary Table S4 for details of the hyperparameter grid for model selection for the G4 models and Supplementary Table S5 for comparison to baseline models. The process for the G2 and G3 models was similar. The G4 model uses 3 width-5 convolutional layers with 32 filters each, and a flattening operation to produce a single embedding for the sequence, followed by a fully-connected layer with 64 hidden units, and a final linear output layer, resulting in about 250K total parameters. All non-linearities are ReLUs.

##### VAE model of NucB homologs

The VAE is a latent variable model with latent variable z, a prior distribution P(z), a conditional distribution P(x|z) over amino acid sequence x parameterized by a ‘decoder’ neural network, and an approximate posterior distribution P(z | x) parameterized by an ‘encoder’ neural network^63,109^. Our model has fully-connected hidden layers of size 64 and 32 in the encoder and 32 and 64 in the decoder, where z has 32 dimensions, and the total number of parameters is about 400K. It is fit on an alignment of 4625 homologs obtained by querying the 2020_06 version of UniRef30 using hhblits ^110^ with an e-value cutoff of 1e-3 and 8 search iterations. Every residue in the WT is designated as belonging to a match state in the alignment, such that the VAE can score mutations to any position.

We employ the sampling technique of Giessel, *et al.*^80^ for sampling variants in the neighborhood of the WT: first, we sample a latent variable from the posterior P(z | WT) and then sample from P(x | z). This provides a principled method for sampling variants of a given reference sequence. We found that unconditional samples from the VAE, or samples from a similar alignment-based method such as a profile HMM, are more diverse than desired.

We assess the ‘naturalness’ of a novel NucB variant by comparing its likelihood under the VAE to that of the WT. Computing the VAE likelihood of a sequence is intractable, however, because it requires marginalizing over the latent variable z. We approximate the likelihood using importance sampling with 2500 samples, using the approximate posterior provided by the encoder as a proposal distribution. Inspired by the finding in Riesselman, *et al.*^63^ that a Bayesian treatment of the neural network parameters improves performance, we additionally average scores across 10 VAE models trained independently with different parameter initializations. In Figure 6a we report the log-likelihood ratio of the variant sequence to the WT. Supplementary Text S3 demonstrates that the VAE is either better than or comparable to a variety of alternative zero-shot scoring methods on our data.

##### Ridge regression model

We fit a ridge regression model to a dataset of variants to estimate the relationship between a sequence and its observed enrichment factor. We tuned the weight on the model’s regularization term to maximize the AUROC score for finding variants with activity comparable or better than the WT, using the same train-test split approach as the CNN. This model is used to estimate the effect of individual mutations, assuming that the fitness landscape is additive. The effect of a given mutation is estimated as the model’s prediction for a sequence defined by applying the mutation to the WT minus the prediction for the WT.

#### Candidate generation using randomized local search

We employ a randomized local search optimizer with many replicates, where each local search trajectory is either initialized at the WT or a top-performing variant from previous rounds of experiments. The optimizer builds up variants one mutation at a time, which enables the search to avoid negative epistasis in the model landscape, but not necessarily discover variants that exhibit positive epistasis^111^. It uses a portfolio of different local search strategies to further improve diversity^112^ and constrains the local search to only propose variants within the trust region, which is defined here simply as the set of all variants within a given radius of the WT.

We proceed with the following steps. First, we curate a list of 15 ‘seed’ sequences that had high activity in the previous round of experiments. Starting at each seed, an optimizer incrementally builds up a population of variants with high model scores by adding mutations to top-performing members of the population. We pool the results across multiple seeds, multiple search algorithms, and 100 random optimization trajectories for each method for each seed.

Each step i of the optimizer can be seen as taking a population P[i] of (sequence, model score) pairs and proposing a new set of child sequences C[i] that hopefully improve the model score. The model is then applied to C[i] and the result is appended to P[i] to obtain P[i+1]. We use three different methods for proposing C[i] from P[i]:

<u>Single-mutant walk</u>: We uniformly sample a parent sequence from one of the top-20 sequences in P[i], then we define C[i] as all possible single-mutation neighbors of the parent.

<u>Sub-sampled single-mutant walk</u>: we uniformly sample (with replacement) 500 different parent sequences from the top-20 sequences in P[i]. For each parent, we sample a single-mutation neighbor.

<u>Regularized evolution:</u>^113^ An extension of the previous method, where the sampling of parent sequences is informed by the model score as well as how recently the sequence was first added to the population.

#### Batch selection

##### Extrapolation score

In G2 and G3, we scored a variant as its number of mutations from the WT. In G4, we used the minimum distance to a high-activity hit from G3 that had been confirmed in a low-throughput plate-based assay (see below). Below, we use ‘distance’ as a synonym for ‘extrapolation score’ to make the algorithm concrete, but in future work it would be natural to perform these batch selection algorithms using alternative scoring methods.

##### Distance-stratified sampling

We group variants by their number of mutations from the WT and select the members of each group with the highest acquisition function score, where the number selected per group follows a target distribution. This distribution was set manually to balance exploration and exploitation and differed across library design methods and rounds. Some library design methods below use no acquisition function. For these, we sample uniformly within each group instead of taking the top-scoring candidates.

##### Distance-stratified sampling with mutation usage filter

We use a greedy algorithm to find a set of candidates obeying the above distance distribution subject to the constraint that each mutation appears no more than M times in the library. First, we express each candidate as a list of one or more (position, new amino acid) mutations from the WT. Next, candidates are bucketed by distance from the WT and we define an iterative sampling procedure that first selects a bucket in accordance with the target distribution and then returns the top-scoring member of the bucket. When a sample is drawn, it is discarded if any of its mutations appear more than M times in the pool of candidates selected so far. This is repeated until a library of the desired size has been selected.

### Instances of TeleProt used in our ML campaign

#### Model-based optimization with a deep neural network (MBO-DNN)

This technique depends on having sufficient data accumulated from prior rounds to fit an accurate sequence-fitness model.

<u>Acquisition function</u>: We score candidates using the CNN model’s predicted log probability of a sequence belonging to the top-scoring discrete activity class. To increase diversity, we clipped model scores such that all variants above some threshold are considered equally attractive for library selection.

<u>Candidate generation</u>: Local search (see below).

<u>Batch selection</u>: Distance-stratified sampling with mutation usage filter.

#### ProSAR

We employ the popular method of Fox, *et al.*^61^ with minimal modifications. This serves as a baseline for the ProSAR+Screen technique below.

<u>Acquisition function</u>: None.

<u>Candidate generation</u>: Using the mutation effect scores from the ridge regression model, we selected all mutations with effect above a given threshold, and sampled new variants containing combinations of 2–6 of these. The recombination is performed in-silico, but traditionally it is done in-vitro. By varying the threshold, we can create libraries with different exploration-exploitation tradeoffs.

<u>Batch selection</u>: Distance-stratified sampling.

#### ProSAR+Screen

We employ a novel extension of ProSAR that screens proposed variants using a model of natural NucB homologs. Essentially, we perform the filtering in Figure 6b for prospective designs. ProSAR relies exclusively on the experimental data to select promising mutations and does not model interactions between these mutations. On the other hand, while MBO-DNN uses a non-additive model with the capacity to capture these interactions, estimating the interactions from experimental data may be challenging in early rounds of experiments, when there is limited data. Instead, Prosar+Screen relies on the patterns in natural NucB homologs to perform negative selection against combinations of mutations that are predicted to exhibit negative epistasis.

<u>Acquisition function</u>: We score variants using their likelihood under the VAE model. Based on the correlation between model score and enzyme fitness observed in Figure 6a, which is consistent with correlations observed in other variant effect scoring datasets, we assume that variants with low score are non-functional, while variants with high score do not necessarily have higher experimental fitness. Therefore, the acquisition function is defined as min(VAE score, threshold), such that all candidates with score above the threshold are treated identically. This low-pass filter helps improve diversity during batch selection. The threshold was determined heuristically, without using experimental NucB activity data, by inspecting the distribution of model scores for randomly-sampled synthetic multi-mutation variants, many of which we expect a-priori to be non-functional.

<u>Candidate generation</u>: Same as ProSAR.

<u>Batch selection</u>: Distance-stratified sampling. Given the thresholding in the acquisition function, within each distance bucket this has the effect of uniformly sampling ProSAR candidates with VAE score above the threshold and using the top-scoring candidates with score below the threshold if insufficient candidates above the threshold are available.

#### Zero-shot sampling

We largely follow the approach of Giessel, *et al.*^80^ for sampling neighbors of the WT using a VAE, except that our candidate generation and batch selection steps have additional strategies for diversifying the library.

<u>Acquisition function</u>: None.

<u>Candidate generation</u>: We sample a large number of WT variants from the VAE using the procedure described in the VAE model section above. Any sample with more than 10 substitutions from the WT or containing gap characters was rejected. In each sample, we revert any mutations appearing outside of the design region (see below). We found that certain mutations appeared in a very large fraction of samples, due to these mutations appearing in homologs with high sequence identity to the WT. To avoid over-using these, we expanded our pool of samples to include pseudo-samples where these mutations were reverted.

<u>Batch selection</u>: Distance-stratified sampling with mutation usage filter.

#### Semi-supervised neighbor sampling

This is a novel extension of zero-shot sampling where the VAE is fit on a combination of natural NucB homologs and high-activity hits from previous rounds. By including the experimental data, the model places high probability mass on mutations that have been observed to improve fitness, while including the evolutionary data enables the model to avoid epistatic interactions among these promising mutations. A key advantage of this approach for future work is that the model is only trained on hits with high activity, which are often easier to identify than collecting a full sequence-fitness dataset.

<u>Acquisition function</u>: None.

<u>Candidate generation</u>: Same as “Zero-shot sampling.”

<u>Batch selection</u>: Distance-stratified sampling with mutation usage filter.

#### Sample+Screen

This approach is a novel extension to MBO-DNN that uses ‘semi-supervised neighbor sampling’ as a proposal distribution for candidate generation instead of local search. When seeking to design variants with many mutations, the search space is very large and local search may break down. Further, placing a simple constraint on candidates’ hamming distance from the WT may be insufficient to constrain the search to regions where the model behaves reliably. Using samples from the semi-supervised VAE, which leverages both natural and experimental data sources, is attractive because it can generate plausible high-fitness samples in a single forward pass of a model instead of relying on a high-dimensional local search procedure.

<u>Acquisition function</u>: Same as MBO-DNN.

<u>Candidate generation</u>: ‘Semi-supervised neighbor sampling’ as a proposal distribution.

<u>Batch selection</u>: Distance-stratified sampling with mutation usage filter.

#### Handling multiple design regions

Due to our DNA assembly protocol, we used multiple 78-amino acid design regions to explore a 110-amino-acid segment of NucB. Zero-shot sampling, semi-supervised neighbor sampling, and ProSAR generate independent samples from a proposal distribution without using an acquisition function. For these, we simply ran our code independently for each design region, and merged the results. For MBO-DNN, ProSAR+Screen, and Sample+Screen, we generated candidates independently for each design region, but then pooled these together for batch selection. Since our regression and classification models are trained on the entire dataset of variants, and scores are comparable across variants with mutations in either design region.

### Non-ML Library Design Approaches

**Hit recombination (HR)** was used as an *in-silico* approximation to an *in-vitro* directed evolution campaign using recombination to introduce sequence diversity. In each round, we took the top K variants with the highest enrichment factor, generated new variants by combining their mutations additively, and randomly selected a subset of the possible combinations to synthesize, stratified by their distances from the WT. The top K were selected from a pool consisting of those proposed by HR in the previous round or those that were a single mutation away from an HR variant, which mainly arise from DNA synthesis errors. Since we trained our ML models on such variants even though they were not explicitly designed by ML in a prior round, we felt that it was fair to provide HR access to them as well, particularly because the diversity in these variants would have been achievable during in vitro directed evolution.

For each pair of hits, we stacked their mutations into a single novel variant, unless the two hits contained mutations at the same position, in which case we created multiple variants. For example, if hit 1 contained mutations {A1C,T4K} and hit 2 contained mutations {R2A,T4S,K5P}, we created {A1C, R2A, T4S, K5P} and {A1C, R2A, T4K, K5P}. Note that this approach is less flexible than in-vitro recombination techniques that may also generate, for example, {A1C, K5P}. See Supplementary Text FS9 for additional details.

**Homolog grafting** is used as a baseline for the zero-shot library. We used the same MSA of natural homologs that the VAE was trained on, and inspected it manually to identify promising combinations of mutations. First, we selected the 18 homologs that had fewer than 25 substitutions from the WT, which mostly come from *B. paralicheniformis* and *B. licheniformis* strains. Then, for each homolog we split the mutations into two subsets, based on which of the two design regions they occur in. For each design region, we created a novel set of variants by applying the corresponding set of mutations to the WT.

**Broad recombination sampling:** Here, we don’t use any ML, and instead simply sample combinations of mutations that the available experimental data suggests may be promising. There are two key differences between this and ‘hit recombination’: we combine a larger footprint of mutations and we combine individual mutations, not groups of mutations that co-occur in known hits. First, we labeled variants as functional or not, using an enrichment factor threshold obtained by visually inspecting histograms of enrichment factors for negative controls and synonyms of the wildtype. Then, we labeled each mutation as ‘observed-functional’ if it appears in at least one functional variant, ‘observed-dead’ if all variants containing it were non-functional, or ‘unobserved’. We sample combinations of these mutations with 2-8 mutations per variant, by uniformly sampling within each group of labels and sampling labels with probabilities (observed_live: 0.97, observed_dead: 0.01, unobserved: 0.02). The hit rate of this library was lower than we had hoped, likely due to our ‘observed-functional’ labeling being too permissive and including observed_dead mutations too frequently in this sampling.

**Loop exploration:** In G3, we included a small library to explore a hypothesis based on the role of mutations to the loop in residues 82-89 in modulating substrate binding. We randomly combined mutations in this region that had appeared in G1 or G2 in at least one variant with comparable or better activity than the WT and used 125 distinct variants at each radius between 3 and 6 mutations.

**Single-site mutants:** In G2 we included all single-mutation variants for all positions in the design region. In G3 we included single-mutation variants at positions that were in the design region for G3 but not G2.

**Assay QC:** Each round included a number of additional variants used for the sake of verifying the consistency and quality of the assay. Our ‘stratified sample’ was obtained by assigning variants to 4 discrete activity levels and then uniformly sampling a few variants from each group. We also included every variant that had been investigated in the previous round’s plate assay described below.

### Directed Evolution (DE) Campaign

A classical directed evolution campaign was pursued in parallel to the designed strategy to compare to ML campaign (Supplementary Figure S18). The sorted library from G1 was used as input for DE2 (i.e. the same library that was used for training the model to design ML2). The DNA sequences of NucB from the input library were cloned and purified before being amplified using a process of PCR shuffling termed StEP recombination ^114^. The variants were then sorted in 8 parallel runs for the highest activity droplets (between the top 0.5–1%). Each run interrogated approximately 2.2 million droplets corresponding to 1.6 million cells. A total of 15,500 droplets passed the sorting thresholds and were collected and recovered for downstream assays and the next round of mutagenesis. A high activity variant with genotype D64S, A73R was isolated from the recovered cells using the liquid culture plate assay.

Variants for the DE3 were generated using a mixture of 1 or 2 rounds of error-prone PCR from the post-sort DE2 library, and 1 round of error-prone PCR from the D64S, A73R isolated variant. All variants were pooled and went through 6 rounds of iterative sorting. The top 0.5% droplets (Or the top 5% cells assuming an average 10% droplet occupancy) were sorted at each round. The post-sort libraries were further validated using the liquid culture plate assay, and hits identified with the specific activity assay. The highest activity variant discovered with the DE campaign had three substitutions: A63S, D64S, A73R.

### Experimental Methods

#### Designed DNA library generation

Library designs were ordered as oligo arrays from Twist Biosciences. The initial site-saturation library mutated residues 33 to 142 of *B. licheniformis* NucB, excluding the native signal peptide sequence and active site residues D93 and N117, which coordinate the active site magnesium. Oligonucleotide synthesis was limited to 300 nucleotides, which was further limited to a design region of 78 amino acids, after subtracting restriction enzyme cut sites and spacers. For simplicity, we did not design variants containing insertions or deletions. Each variant contained mutations only in a single *design region*. These were chosen to maximize the proportion of promising mutations while exploring as much of the protein as possible. G2 used a single design region at residues 33–110 and G3/G4 used two overlapping regions: residues 33–110 and 65–142.

Due to errors during DNA synthesis and amplification, the DNA libraries contained a significant number of distinct sequences beyond what was designed. 14506 of the 55760 variants that had sufficient abundance to characterize enzyme activity came from such errors. These variants are not used in our hit rate analysis, but they were utilized when designing variants, since these non-designed variants sometimes displayed non-zero activity because they often result from crossover between designed variants.

*In-silico* design methods generated amino acid sequences, which were converted to DNA sequences as follows. All unchanged codons used the WT (*B. licheniformis*) codon. Mutated codons used the canonical codon given by the *B. subtilis* codon usage table ^115^. Any designs with a *BsaI* cleavage motif (GGTCTC or GAGACC) had their first codon substituted with the next-most-likely *B. subtilis* codon

#### Random DNA library generation

We used both error-prone PCR (epPCR) (Agilent Genemorph II error-prone PCR kit) and shuffling by the Staggered Extension (StEP) Process PCR^114^. epPCR was carried out on the full length of the mature peptide according to the manufacturer’s recommendations and subsequently purified by agarose size selection. StEP was performed on enriched library variants using a PCR protocol adapted to the KAPA HiFi HotStart ReadyMix (Roche). To maximize recombination, we used a 35–40°C extension temperature, with 100 cycles of 5s extension. Products were purified by agarose size selection using the Zymoclean™ Gel DNA Recovery Kit and run on a gel, size selected and purified.

Since StEP PCR can regenerate many parental sequences, COLD-PCR^116^ was performed to enrich non-parental sequences, using an 85°C denaturation temperature for NucB which decreases PCR efficiency for DNA homodimers, but not heterodimers.

#### Nuclease expression in *Bacillus subtilis* host strain

Libraries were cloned into an IPTG-inducible shuttle vector which was ultimated integrated into *B. subtilis* for subsequent screening. The hybrid T7-lacO promoter^117^ was employed due to its low background expression in E. coli. To further reduce leaky expression in *E. coli*, a copy of LacI is included in the backbone of the shuttle vector. T7 RNA polymerase required to transcribe the gene of interest is present only in the host *Bacillus* strain. IPTG-induced expression in this system was found to be equal or greater than expression from the constitutive Pveg promoter.

##### Expression vector construction

Plasmid libraries were assembled using either Gibson or Golden Gate assembly. For Gibson assembly, the NEBuilder Hifi DNA Assembly Master Mix (NEB E2621) was used according to the manufacturer’s instructions. For Golden Gate assembly, the NEB® Golden Gate Assembly Kit (BsaI-HF®v2) (E1601) was used according to the manufacturer’s instructions. A diagram of the plasmid construct is presented in Supplementary Figure S17a. Flanking sequences from amyE allow for integration of these constructs in B. subtilis via homologous recombination at the amyE site.

Assembled plasmids were transformed into *E. coli* for propagation by electroporation of 2-µL of a 1:1 dilution of the assembly reaction in nuclease free water. E. cloni^®^ 10G SUPREME electrocompetent cells (Lucigen 60080-2) were loaded into a 1-mm cuvette as per the manufacturer’s instructions. To determine transformation efficiencies, 10, 100, and 900 µL of recovery volume were plated on LB agar plates containing the appropriate antibiotic then incubated overnight at 37°C. The assembly mix was stored at –20°C until the transformation efficiencies were determined, then the number of reactions were scaled as needed to obtain sufficient library representation. Libraries were plated on 245-mm square LB agar trays (1–2mL recovery volume for each plate). After picking colonies for library QC, the q-trays and plates were scraped with 5 mL liquid LB and pooled. The plasmids were purified by ZymoPURE II Maxiprep kit from 10 mL each of pelleted cells.

##### Bacillus subtilis host strain

We used *B. subtilis* strain SCK6 (Bacillus Genetic Stock Center ID 1A976) from Zhang and Zhang^118^. Overexpression of the master regulator ComK in this strain enables high transformation efficiencies.

##### Bacillus subtilis library transformation

The host strain was prepared for transformation by xylose-induction of ComK then frozen at –80°C as per Zhang and Zhang ^118^. Frozen aliquots of competent cells were allowed to thaw at room temperature for 15 minutes. Transformations were carried out by adding 1 µg of the plasmid libraries to 1 mL of cells in sterile, round-bottom tubes and gently mixing. The tubes were incubated at 37°C incubator for 1.5 h while shaking. Transformation efficiencies were determined by plating 1 µL of the recovery mixture on an LB agar plate containing 10 ug/mL kanamycin, 5 ug/mL erythromycin, and 100 ug/mL spectinomycin (subsequently referred to as LB-KES). Libraries were subsequently plated as 200-µL volumes onto solid agar plates (245-mm, LB-KES, Teknova L4795), yielding approximately 100,000 cfu after overnight incubation at 37°C. The number of plates were scaled as required to obtain sufficient library representation.

#### Protein production and harvest

Strains with high activity in liquid culture plates were selected for protein expression. For each strain, two 2-L baffled flasks with 500-mL LB-KES, 100 mM MOPS, pH 7.2, 2% glucose were inoculated with 5 mL overnight cultures incubated at 37 °C, shaking at 220 rpm. Once the culture reached an OD of 0.8 (approximately 4h), IPTG was added to a final concentration of 0.5 mM to induce protein production overnight. Secreted NucB nuclease was harvested from the spent media (around 880 mL total) by centrifugation at 6000 rcf for 20 min and subsequently filtered across a 0.2 µm membrane.

#### Protein purification

We purified the NucB nuclease based on the method described in Rostami, *et al.*^47^ using two chromatography steps. Briefly, proteins were precipitated from spent media using saturated ammonium sulfate solution to 65% final saturation (∼1800 mL per liter of culture) and stirred at 4°C overnight. The precipitated protein was pelleted by centrifuging at 22,000 rcf at 4°C for 60 min. The pellet was resuspended at 4°C in 5 mL of 50 mM potassium phosphate, pH 7.2, 1.5 mM β-mercaptoethanol (BME), and protease inhibitors (Roche cOmplete) to a total volume of approximately 10 mL. After sterile filtration, the protein solution was desalted by FPLC (Akta start, Cytiva) at 5 mL/min on a pre-equilibrated XK 26/10 Hitrap Desalt column (Cytiva). The mobile phase was 50 mM potassium phosphate at pH 7.2, 1.5 mM BME, and 0.5 mM PMSF. Fractions containing NucB were pooled and subsequently purified by ion exchange. Samples were loaded onto a 5-mL Q-Sepharose HiTrap column, pre-equilibrated with the same mobile phase as used in desalting. NucB was eluted with 2 M NaCl in 25 mM Tris, pH 9, 1.5 mM BME, and 0.5mM PMSF. The fractions containing NucB were analyzed by SDS-PAGE (Bio-Rad Criterion AnykD, 26-well). Fractions were concentrated using 3 kDa cutoff spin concentrators (Amicon® Ultra-15 Centrifugal Filter Unit) to a final volume of 5 mL and then dialyzed twice against 2L of 10 mM Tris, pH 9.0.

#### Nuclease activity assays

To select the set of variants to more-deeply characterize in our low-throughput, high-resolution assays, we used three different assays in series to isolate the top performers. 1) First, we used our ultra-high-throughput droplet-based microfluidics platform (“Automated Droplet-based Fluorescence Assay”) to iteratively sort at increasing fluorescence thresholds (Figure 2a). 2) Cells that were sorted were then preliminarily assessed for activity in a microtiter plate-based enzyme kinetic assay using cultured strains (“Liquid Culture Plate Assay”). The top variants from the liquid culture plate assay were then assessed with 3) a microtiter plate-based enzyme kinetic assay using purified enzyme (“Specific Activity Assay”). Finally, the best variants from each campaign were assayed for biofilm disruption (“Biofilm disruption assay“). This progression of assays trades throughput for higher resolution (Supplemental Figure S7).

Nuclease activity was assayed with DNA hydrolysis probes consisting of two sets of duplexed 20-mer oligos: one labeled with a fluorescent probe and quencher, and one unlabeled. The ratio of the two substrates was tuned to maximize the linear response curve for nuclease activity. Endonuclease activity of NucB releases the quencher on the 3’ end of G537 (3’ Iowa Black® FQ) effectively dequenching the fluorescent probe at the 5’ end of G651 (5’ Alexa Fluor® 488).

#### Automated Droplet-based Fluorescence Assay

To measure the activity of millions of nuclease enzyme variants, we built a custom microfluidic platform and devices to conduct ultra-high-throughput screening in droplet-based microfluidics^26^. Cells in expression media with 0.1% (w/v) Pluronic F-127) were encapsulated as a water-in-oil emulsion in fluorocarbon oil, stabilized with a biocompatible surfactant on a custom microfluidic chip. Cells were prepared for encapsulation by culturing overnight at 37°C in LB-KES, then back-diluted 1:200 and grown for 8h to maximize cell viability. Cells were then washed and diluted to OD 0.08 to ensure a lambda of 0.1 in drops according to the Poisson distribution (average of 1 droplet containing a cell out of 10 total droplets). The low density minimized the chance of two cells being encapsulated in the same droplet, preventing false positives. Droplet diameters were tuned to 20 µm (∼5 pL volume). Droplets were incubated at 37°C for 16 h for cell growth and nuclease expression, then passed through a second microfluidic chip where assay reagents were introduced via picoinjection^119^, incubated on-chip for 4 seconds, sorted by fluorescence intensity, then merged into an aqueous recovery channel. The picoinjected assay solution consisted of 100 mM MOPS, pH 7, 2.5 mM MnSO4, 2 µM G651/G537, 50 µM G24/G25 and 0.1% (w/v) Pluronic F-127. Sorted cells were fully recovered by breaking any residual emulsion with perfluorooctanol (PFO) and plating ∼100 drops to estimate the recovery rate. The remainder of recovered cells were grown in 25 mL of LB+KES media overnight at 37°C.

#### Liquid Culture Plate Assay

The liquid culture plate assay was used to isolate hits obtained from the ultra-high-throughput screen. Optical density was used to normalize the measured activity, and therefore does not control explicitly for enzyme titer. The culture for plate validation was prepared in two stages: the growth phase, followed by the induction phase.

In the growth phase, a 96-deep-well plate was filled with 1 mL per well of LB-KES broth. Each well was then inoculated with a single colony. To maintain sterility while allowing gas exchange, the plate was covered with a breathable seal. The plate was then placed in an incubator set at 37°C, 750 rpm for 8 hours. After the incubation period, the optical density (OD) of the cultures was measured at a wavelength of 600 nm. For the OD measurement, 10 uL of each culture was diluted in 90 uL of 150 mM NaCl saline solution. Background correction was performed considering a 10-fold dilution in 100 uL, resulting in an approximate value of 0.094 absorbance units (AU). Additionally, a volume correction factor of approximately 0.3 was applied due to the 100 uL pathlength used for the measurements. Basic normalization for growth in the liquid culture was accomplished by dividing v_max_ by OD600 of each culture.

In the induction phase, another 96-deep-well plate was filled with 1 mL per well of expression media (LB-KES supplemented with 100 mM MOPS, pH 7.2, 2% glucose, and 0.5 mM IPTG). The 8-hour cultures were used to inoculate this plate, with 5 µL added per well. As before, the plate was covered with a breathable seal and then incubated at 37°C, 750 rpm for a minimum of 16 h.

#### Specific Activity Assay

Specific activity measurements for selected variants are presented in Figure 3b. Protein purification is a labor-intensive process that must be performed serially for each strain. The following variants were purified: the top-performing variants from each ML generation, the top performing strain for the DE campaign, and the wildtype enzyme. Each variant was evaluated at 4 concentrations (12.5, 25, 50 and 100 ng/mL) and its v_max_ was normalized to the kinetic rate of the wildtype enzyme at the same concentration, resulting in a measurement of fold-change activity. To determine whether one variant was higher activity than another, a permutation test was performed on the average of the fold-change activities measured at the 4 concentrations.

Both the liquid culture and specific activity kinetic assays measured activity at 25°C in 200mM MOPS pH7 buffer, 5mM MnSO4, 0.2% Pluronic in 384-well plates (Corning 3577). The fluorescently-labeled substrate (G651/G537) was present in the assay at 0.4 µM and the unlabeled substrate (G24/G25) was at 100 µM. Fluorescence was measured over a 40-60 minute timecourse with excitation at 495 nm and emission at 525 nm at fixed interval (12s for specific activity assay, and 1 min for liquid culture assay). Activity was calculated by fitting the linear slope of the initial fluorescence intensity change^120^. Examples of kinetic curves are shown in Supplementary Figure S14.

#### Biofilm disruption assay

In Figure 3c and Supplementary Figure S15 we evaluate the ability of the purified nuclease variants to disrupt *Bacillus licheniformis* biofilms. *B. licheniformis* was cultured in 10 mL of LB at 37°C and 220 rpm for 2 days. The culture was then diluted 1:100 into fresh LB media and dispensed in 150-µL volumes into the wells of sterile, 96-well, clear round-bottom plates for biofilm to develop for 2 days at 37°C and 100 rpm. The biofilms were disrupted by pipetting, and the residual biofilm left on the plate walls was gently washed three times with 200 µL saline. The washed residual biofilm was incubated at 37°C and 100 rpm with 200 µL of purified enzyme (a dilution series in 25 mM Tris, pH 7.5, 0.1% pluronic and 2.5 mM MnSO_4_) for 20 h. The remaining biofilm was washed in 200 µL saline three times, and incubated with 200 µL of 0.1% crystal violet for 10 minutes at room temperature before being removed by aspiration, and washed once with saline. The now-stained, residual biofilms were then disrupted by adding 200 µL of 96% EtOH and 2% acetic acid and transferred to a flat-bottom plate to measure absorbance at 590 nm. Each variant’s biofilm disruption ability is determined by dividing absorbance at 590 nm by the average absorbance of a no-enzyme control, which yields the percentage of biofilm remaining after digestion. To infer dose-response curves, we fit a Hill equation to the observed biofilm degradation using scipy.optimize.curve_fit. To estimate confidence bands, we took 200 resamples of the Hill equation parameters from the inferred multivariate normal distribution.

#### Selecting fluorescence thresholds for ultra-high-throughput sorting

In each round, the population of cells was sorted in parallel at multiple fluorescence thresholds, where each threshold was determined by the fluorescence of a fiducial enzyme sequence with known activity. Droplets of a sample of the library and of the fiducial sequences were generated and varying quantities of mFluor Green 620 acid (AAT Bioquest catalog number 1144) were added to the droplets before emulsification to differentiate the droplet samples in the assay. Droplets were then pooled and assayed for fluorescence intensity in our microfluidic platform. A 488-nm laser was used to excite both the mFluor Green 620 acid and the 5’ Alexa Fluor® 488, which were measured with TRITC/CY3 620/52nm (Thorlabs, MF620-52) and FITC 530/43nm (Thorlabs, MF530-43) BP Emission Filters, respectively. Gates were selected relative to the activity of the fiducial sequences. For example, gates with fluorescence threshold at the center of the fiducial distribution are used to categorize variants having equivalent activity to the fiducial sequence. Gates set at the right tail (i.e. greater than the mean of the fiducial fluorescence distribution) are used to categorize variants as having improved activity relative to the fiducial (See Supplementary Figure S27).

#### Next-generation sequencing (NGS)

Variant sequences were determined by next-generation sequencing on the MiSeq platform with paired-end reads. Variable regions from each library were amplified by PCR with forward primer sequence TATTCTCCTCAGCATGCCGAAGGCG and reverse primer sequence AGATTACTACTGATAACCACTGTTGATTG. The adapter and sequencing primers are appended to the 3’ end of the primers, with a variable 0–7 base-pair spacer to enable demultiplexing of pooled libraries. The fastq files were preprocessed using fastp, and aligned to the reference DNA sequence using Bowtie2. The details of the sequencing depth is summarized in Supplementary Table S6.

### Processing data from ultra-high-throughput experiments

We convert the observed NGS counts to discrete enzyme activity labels for the following reasons. First, since cell sorting is a discrete procedure, using discrete labels better reflects the nature of the underlying measurement, i.e. that variants that passed through the sort achieved the fluorescence threshold. Second, estimates of enrichment factors include noise from biological and technical variance. By discretizing, we can reduce models’ overfitting to noise, and for these reasons we train multi-class classification models. Third, discrete labels allow us to analyze a library’s hit-rate, which is easily interpretable. Finally, by using discrete activity buckets derived from fiducial sequences that appear in multiple rounds of experiments, it is straightforward to aggregate measurements from multiple rounds to create a single genotype-phenotype dataset (Supplementary Figure S24). See Supplementary Figure S23 for a schematic overview of the data processing procedure.

**Computing enrichment factors:** We perform NGS on both the pre-sort (input) library and the post-sort (output) variant pools. For a variant V, we estimate its abundance as:

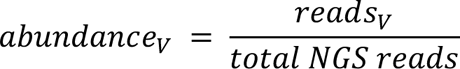

The *enrichment factor* of V is the ratio of post-sort abundance to pre-sort abundance:

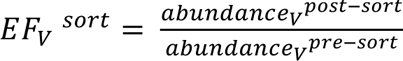

Estimating the EF is sensitive to errors in the denominator. Therefore, we only kept variants that had abundance above a given threshold in the input pool. For example, in G4, only variants with ≥50 counts in the input pool were included, which left 15404 variants whose enrichment factors could be accurately estimated. This is only 20% of the total variants present with ≥2 reads (see Supplementary Figure S25).

**Fiducial sequences:** To interpret the enrichment factor of a screened variant, we compared it to the distribution of enrichment factors for a given fiducial sequence with known activity. In each library, we included multiple synonymous variants of each fiducial, where these can be distinguished during sequencing. Variants’ enrichment factors for each sort were computed after translating to amino-acid space, while fiducial enrichment factors were computed based on DNA sequence reads. Variability in the enrichment factors of synonymous variants allows us to account for noise in the assay (i.e. from variability in growth rate, expression level, cell cycle, sorting, and other sources of technical and biological variability).

**Constructing Fiducial Sequences:** Synonymous variants were constructed by taking a stratified sample over the number of codons to modify. For a given number of codons C, C unchanged codons (i.e. those that did not include any amino acid mutations relative to the wildtype) were selected uniformly at random and changed to the next most common synonymous codon in the *B. subtilis* codon usage table. Error-prone PCR introduced synonymous codons and premature stop codons in G1. In each subsequent (designed) generation we picked fiducial sequences based on variants that had known activity confirmed in specific activity assays. In G2 and G3, we introduced synonyms of the wildtype and A73R, the best variant discovered in G1. In G4, we additionally included synonyms of (A73R, D74S) and (A63P, A73R, D74H, I84Y), found to have high activity in G3. Fiducials were designed for each design region. See Supplementary Text S10 for details.

**Hit-calling relative to a fiducial:** To determine that a variant has activity greater than a fiducial in a given sort, we compute the t-statistic and associated p-value for the right-sided test where the null distribution is given by the fiducial’s enrichment factors (see Supplementary Figure S24) and the observation contains a single enrichment factor for the variant EF_V_^sort^. Given a set of variants with associated p-values, we assign discrete labels to them jointly, controlling the overall expected false discovery rate (EFDR) by applying a Benjamini-Hochberg (BH) correction ^51^. A variant with a significant p-value after correction is labeled as “hit”, but this labeling will vary depending on the context. For example, at times we report the hit rate of an entire library design method, while elsewhere we report separate hit rates of that method for variants at different distances from the WT. Here, the BH correction is applied within each pool of variants that a hit rate is reported for. We use an EFDR of 0.1 throughout the paper.

**Obtaining resolution for wide-ranging activity levels:** To characterize variants into multiple discrete activity levels that cover a broad range of enzyme activity, we sorted in parallel at multiple fluorescence thresholds (Figure 1c). This is necessary because the enrichment factor is a sigmoidal attenuation of the underlying enzyme activity. For example, high activity variants cannot be distinguished by sorts performed at low fluorescence threshold (Supplementary Figure S24) because nearly all high-activity variants have the same sorting outcome. For each generation, fluorescence thresholds were chosen prior to sorting according to the procedure outlined above.

**Obtaining the final genotype-phenotype dataset.** To construct the final 55760-variant dataset of sequences from all generations, we needed to reconcile multiple, potentially-conflicting, activity labels for a given variant. Within a generation, the hit-calling with respect to multiple fiducials may be inconsistent, since different fiducials may use data from different gates. This does not guarantee, for example, that if we label a variant as better than WT we also label it as better than the negative control. Second, the labeling may be inconsistent across generations. For labels that appear in multiple generations, we take the label from the latest generation. Within a generation, we further assign each variant the highest activity label for which it was labeled as better than the corresponding fiducial. For example, 5 variants across G3 and G4 failed to reject the null when compared to the negative control, but rejected the null when compared to the WT and A73R fiducials and 973 variants across G3 and G4 failed to reject the null when compared to the negative control, but rejected the null when compared to the WT.

### Quantifying diversity with sequence clustering

We used a method inspired by the clustering in UniRef, where clusters are defined by the maximum distance from the seed: i.e. in UniRef90 clusters are composed of sequences that have at least 90% sequence identity with the seed sequence of the cluster^121^. To build clusters for a given set of variants, we first set the maximum distance D between two sequences in the same cluster. Then, we build a dendrogram between sequences by starting with one node per sequence and iteratively connecting two nodes if the distance separating them is the smallest among all pairwise distances. The distance between two nodes u and v is given by the “farthest point” inter-cluster distance i.e. d(u, v) = max_ij_ d(u[i], v[j]). Nodes are joined iteratively until all sequences are connected. Clusters are formed by the largest nodes that contain sequences whose maximum pairwise distance is no larger than D.

### Quantifying negative epistasis

Epistasis is often quantified for continuous fitness measurements, where it measures the extent to which the impact of mutations on fitness is non-additive^111^. For our discrete activity labels, we instead measure the extent to which the impact of applying multiple mutations to the protein renders it functional vs. non-functional in a way that is consistent with mutations impacting the protein independently. Many protein engineering approaches seek to combine mutations that have been previously observed to maintain or improve protein function (i.e., they have been labeled as having non-zero activity). Therefore, we investigate the rate of negative epistasis: how often combining such mutations produces a non-functional protein.

Our dataset contains experimental measurements for nearly all single-mutation variants of the WT. From this, we obtain a list of mutations that were observed to not destroy the function of the protein using our above labeling scheme. Then, we select the set of multi-mutation variants that only contain mutations in this list. The negative epistasis rate is defined as the fraction of such variants that are non-functional. Under a null model of the protein fitness landscape where mutations impact the protein independently, we would expect the negative epistasis rate to be 0. Given that the labeling scheme has an expected false positive rate of 0.1, and an additional chance of false-negatives, however, the expected empirical epistasis rate is non-zero even under the null model. We only report negative epistasis for double-mutation variants, where the impact of false positives on the analysis is smaller than it would be for higher-order mutants.

## Acknowledgments

We thank the entire Triplebar team for their support. For helpful feedback and discussions, we would like to thank Andreea Gane, Ankur Parikh, Christof Angermueller, David Andre, Hemai Parthasarathy, Jue Wang, Mathias Voges, Melis Anahtar, Nilesh Tripuraneni, Tim Beissinger, Zach Wu, and Zelda Mariet.

## Author contributions

Conceptualization: NT, DB, JJA, LJC. Software: NT, DB. Investigation: NT, DB, HL, KH, MC, KI, VP, KDN, LF. Formal Analysis: NT, DB, CX. Visualization: NT, DB, CX, JWK. Project administration: AR, KGH, JJA, LJC. Supervision: JJA, LJC. Writing – original draft: NT, DB, CX, KH. Writing – review & editing: NT, DB, CX, KH, MC, KI, VP, KDN, JWK, CE, JJA, LJC

## Supplementary

**Supplementary Figure S1:**
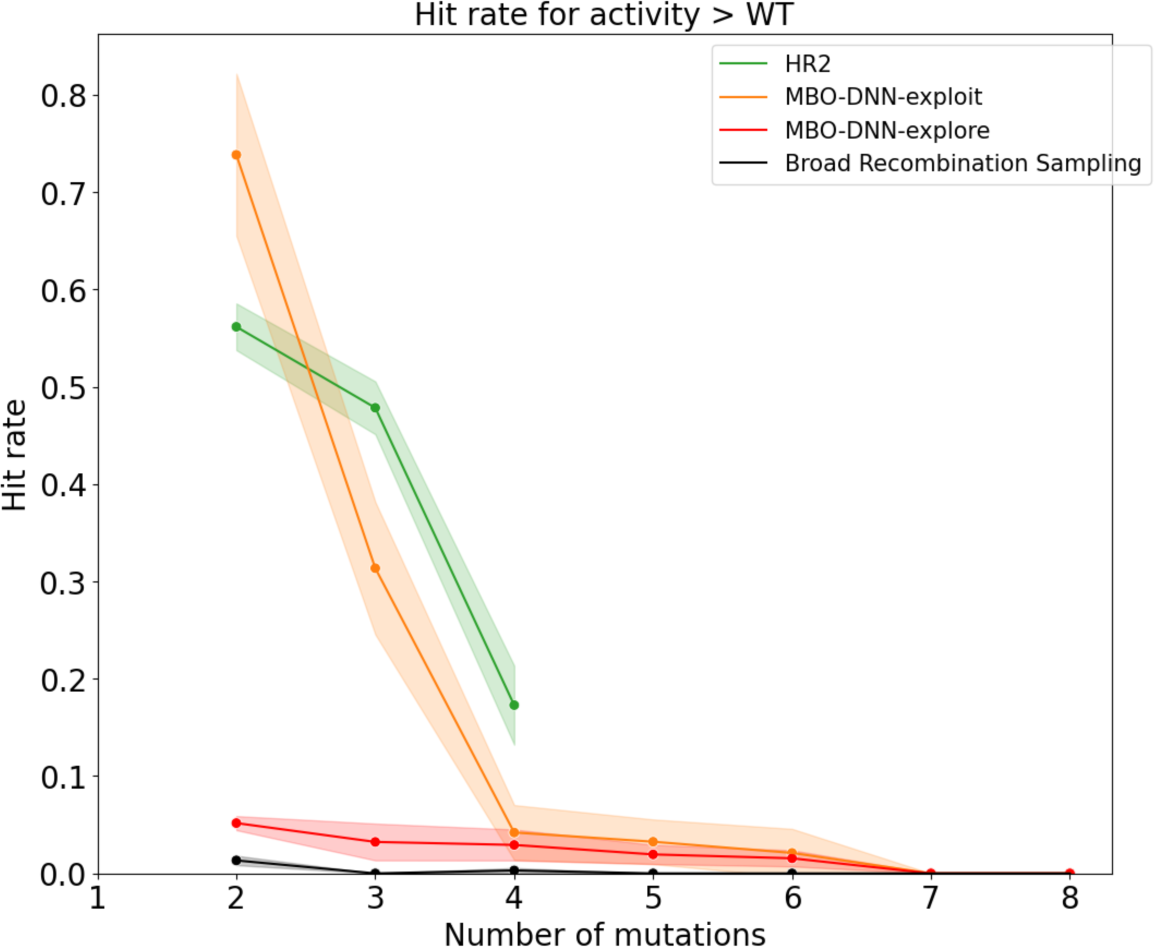
hit rates for key sub-libraries in G2.

**Supplementary Figure S2:**
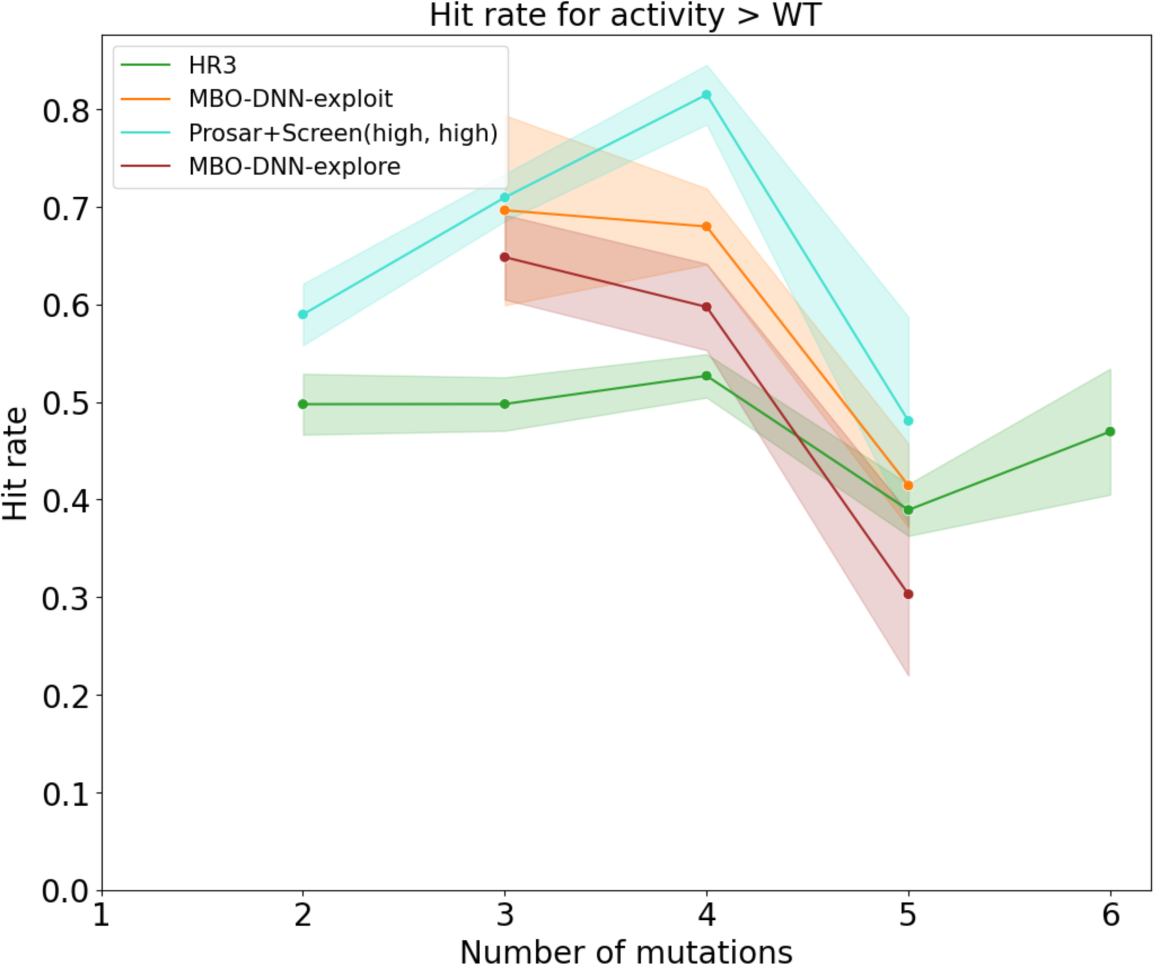
hit rates for key sub-libraries in G3.

**Supplementary Figure S3:**
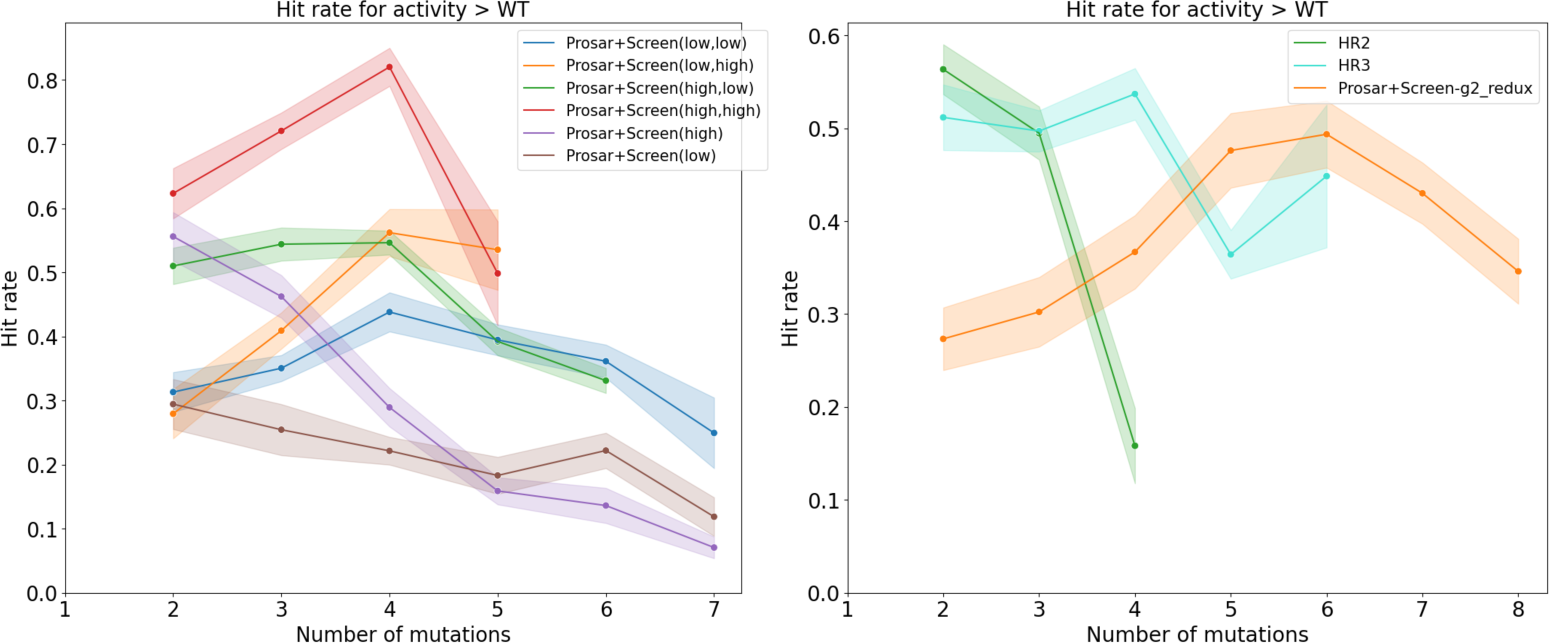
(left) Hit rates for ablations of the Prosar+Screen method where we vary the threshold used for selecting mutations to include in the combinatorial library and the threshold used for dropping recombination variants with low VAE score. (right) Using Prosar+Screen as an alternative design approach in G2 has a similar hit rate to HR2, but the hits include far more mutations per variant.

**Supplementary Figure S4:**
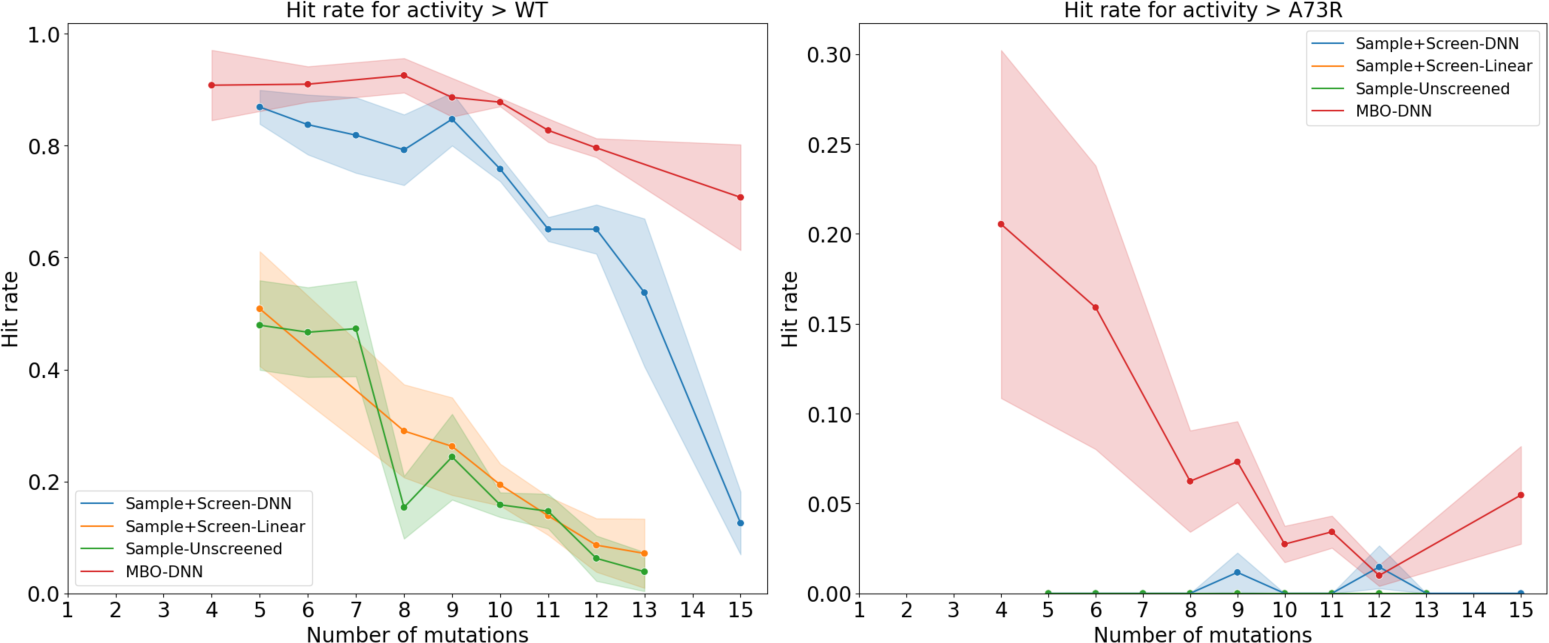
Contrasting the performance of different methods for finding sequences with high model score (Sample+Screen-DNN vs. MBO-DNN) and different types of models (Sample+Screen-DNN vs. Sample+Screen-Linear vs. Sample-Unscreened).

**Supplementary Figure S5:**
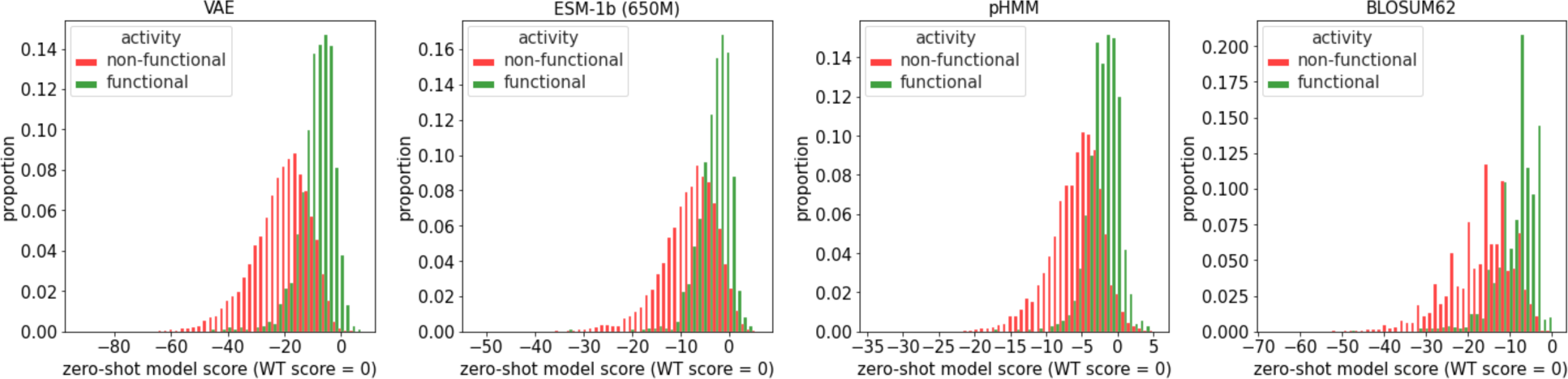
Histograms of zero-shot scores for functional and non-functional G1 variants. All methods provide some separation between functional and non-functional variants, with the distributions most separated for the VAE.

**Supplementary Figure S6:**
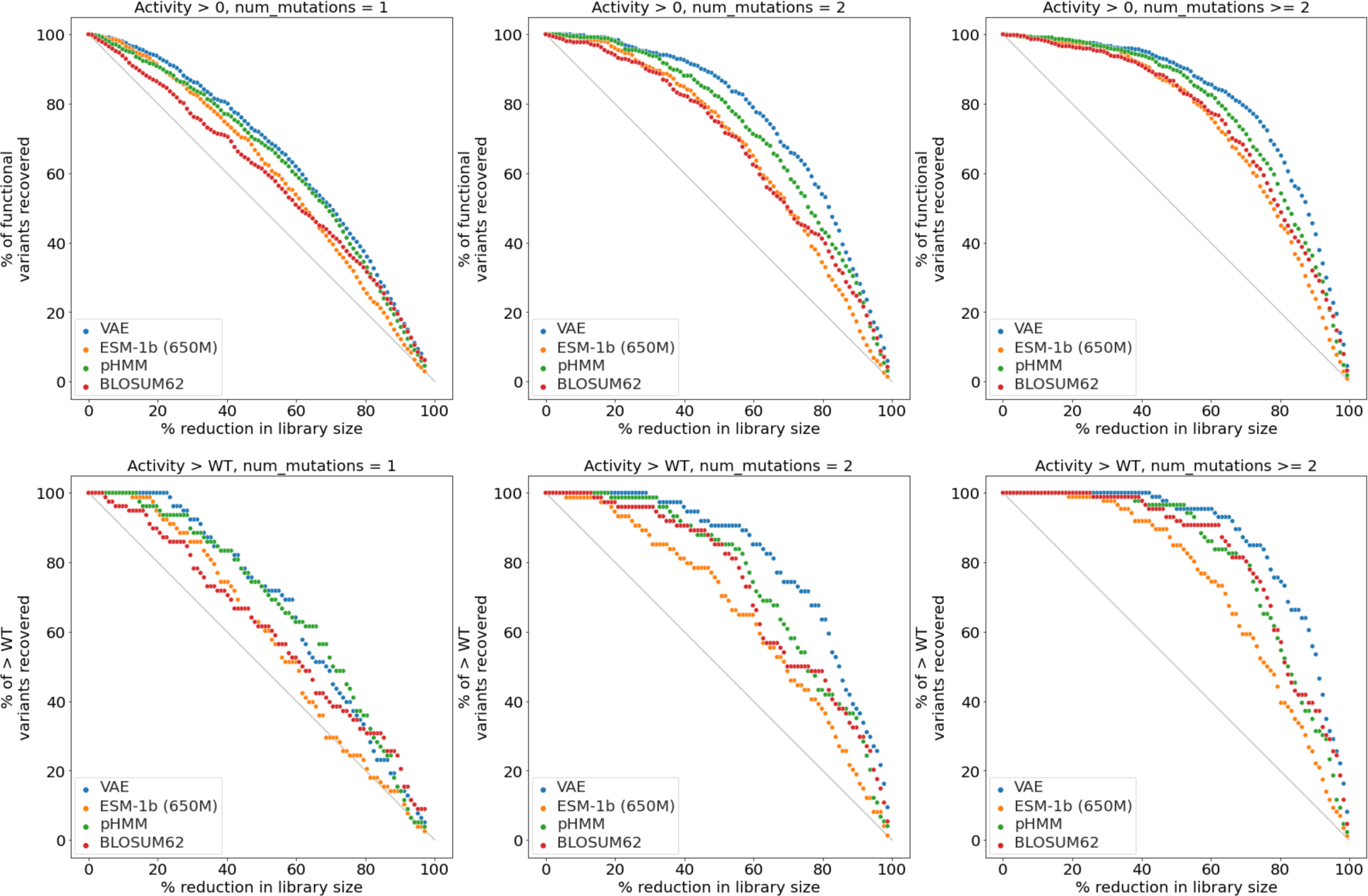
Effectiveness of sub-sampling the G1 library using various zero-shot scoring methods. Sweeping over the X axis provides a tradeoff between how much the library size is reduced and how many variants with target activity are discovered. Across values dataset reduction percentages, the VAE performs comparably or better than alternative methods at discovering such variants. Subfigure rows: which level of activity is used to define a hit. Subfigure columns: which set of variants is analyzed.

**Supplemental Figure S7:**
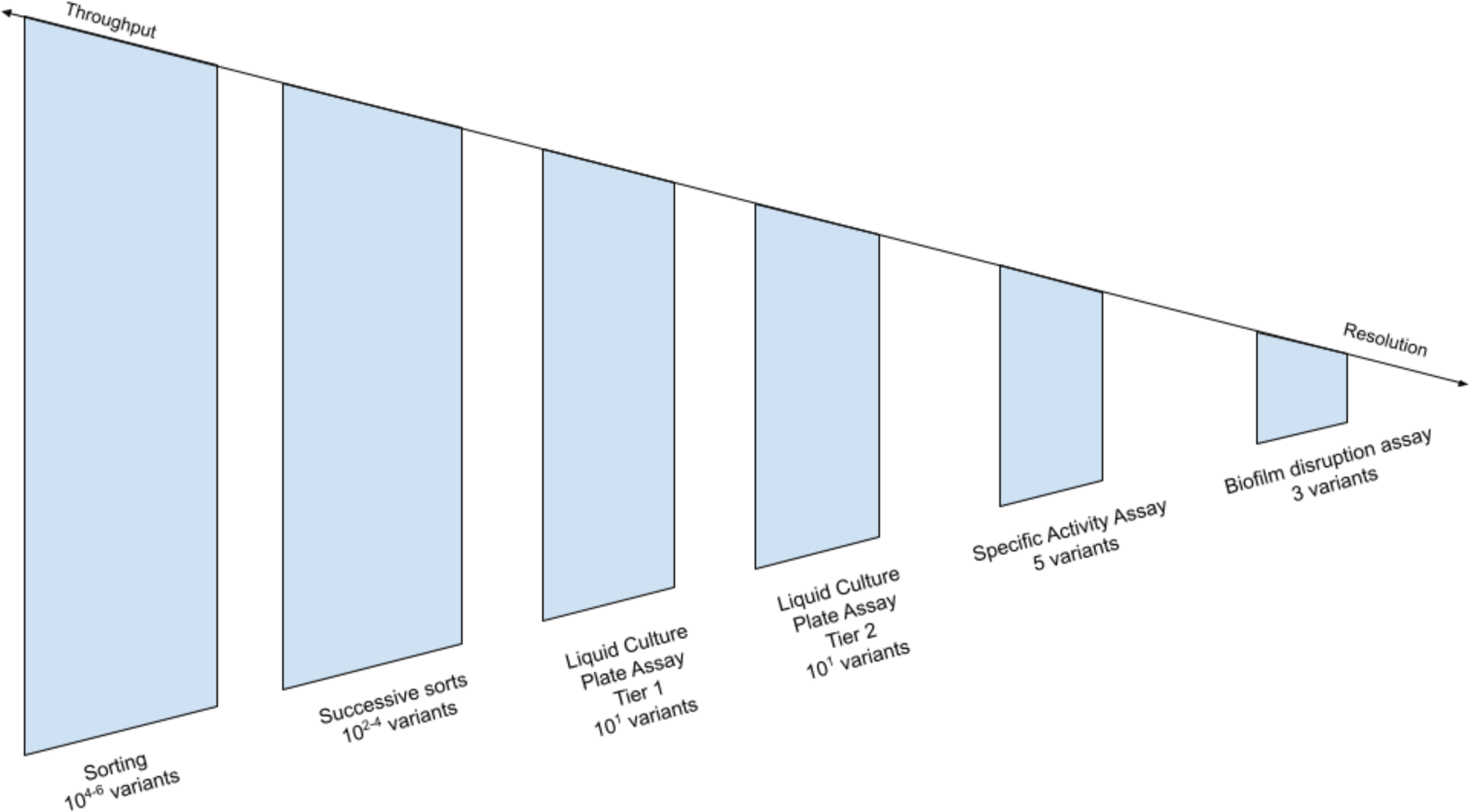
The sequence of filtering steps leading from our highest throughput assays to our highest resolution assays. Our selection procedure proceeds as follows, with the best performers promoted to the next assay at each stage: we screen millions of variants in high throughput, successively sorting at higher fluorescence thresholds. We then screen tens of variants in our plate assay (Tier 1) and verify them with replicates (Tier 2). We then purify the top several performers and screen for specific activity. The very best performers are validated in a biofilm disruption assay.

**Supplementary Figure S8:**
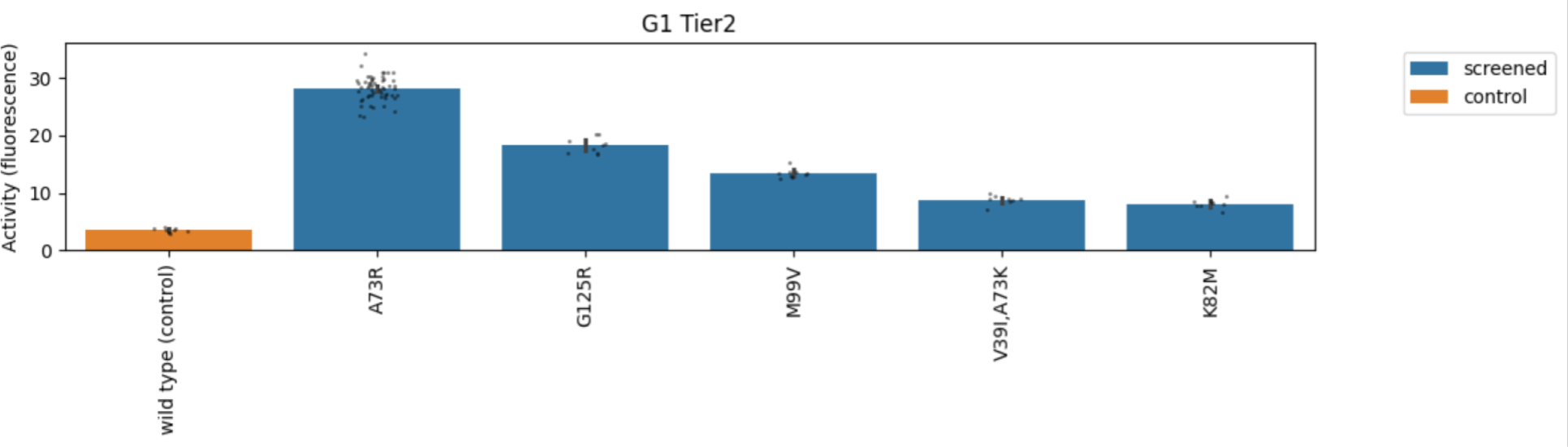
G1 Tier 2 liquid culture plate assay activity assessment. The y axis (“activity”) measures fluorescence normalized by optical density. On the x axis are genotyped strains. Variants in orange were added to plates as controls. A73R was selected for purification. Error bars represent ±1 standard error of the estimated activity.

**Supplementary Figure S9:**
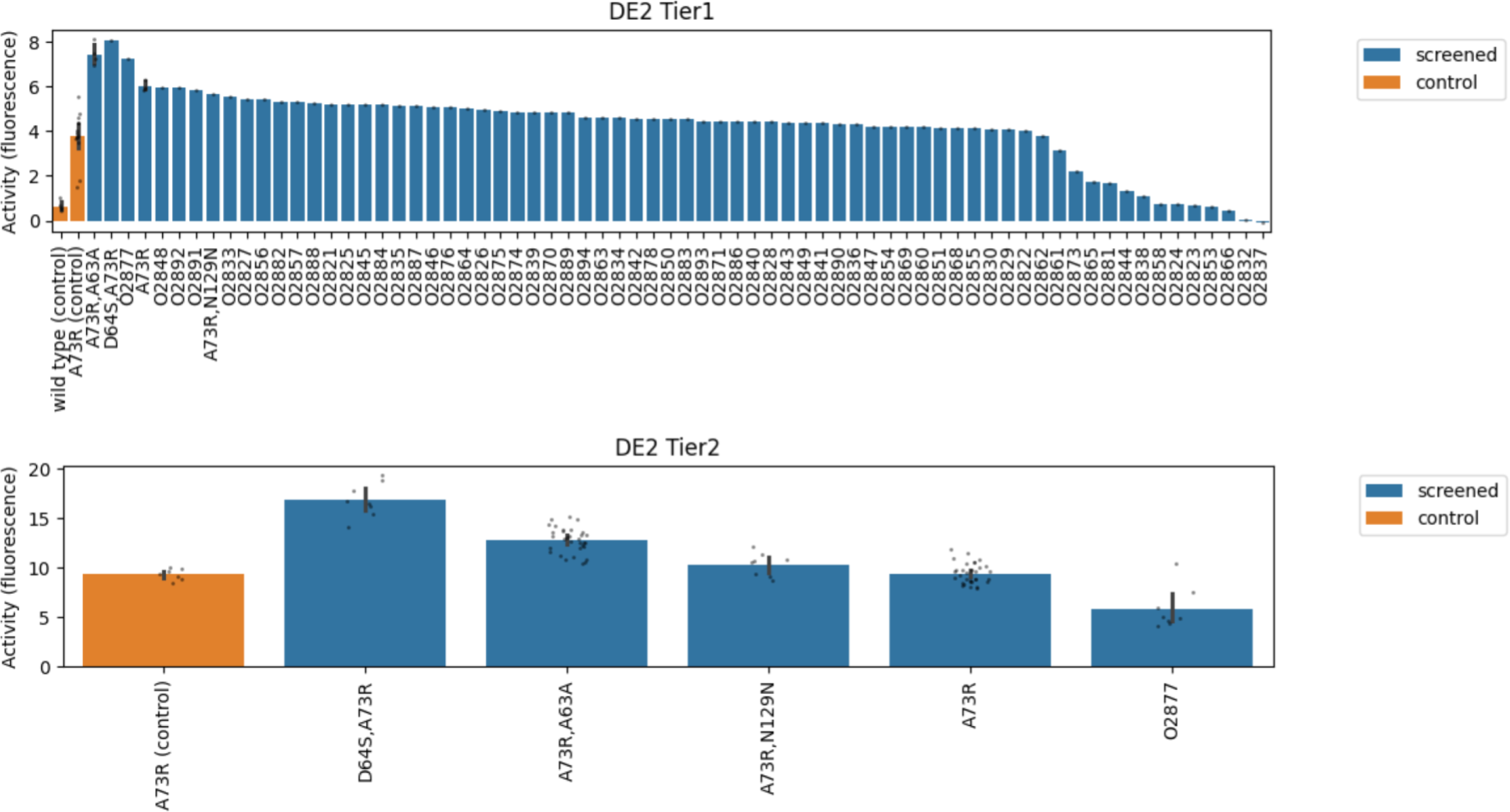
DE2 Tier 1 and 2 liquid culture plate assay activity assessment. The y axis (“activity”) measures fluorescence normalized by optical density. On the x axis are genotyped strains. Variants in orange were added to plates as controls. In Tier 1, variants that were not selected were not genotyped and are reported with their numerical strain identifier starting with “O”. D64S,A73R has the highest activity in the Tier 2 screen. Error bars represent ±1 standard error of the estimated activity.

**Supplementary Figure S10:**
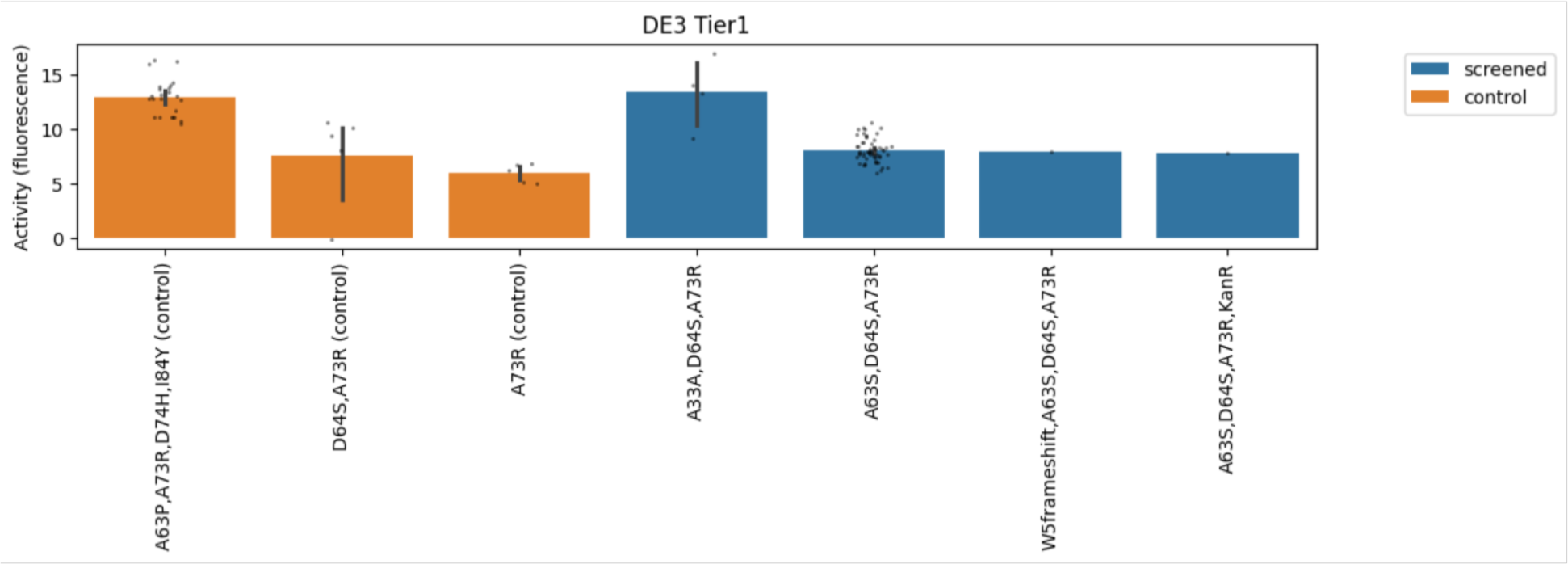
DE3 Tier 1 liquid culture plate assay activity assessment. The y axis (“activity”) measures fluorescence normalized by optical density. On the x axis are genotyped strains. Variants in orange were added to plates as controls. A63S,D64S,A73R was selected for purification. A Tier 2 screen was unnecessary because there were sufficient replicates of each genotype Tier 1. Despite its high activity, A33A,D64A,A73R contained a promoter mutation and was excluded from further analysis due to a change in titer confirmed by SDS PAGE (data not shown). Error bars represent ±1 standard error of the estimated activity.

**Supplementary Figure S11:**
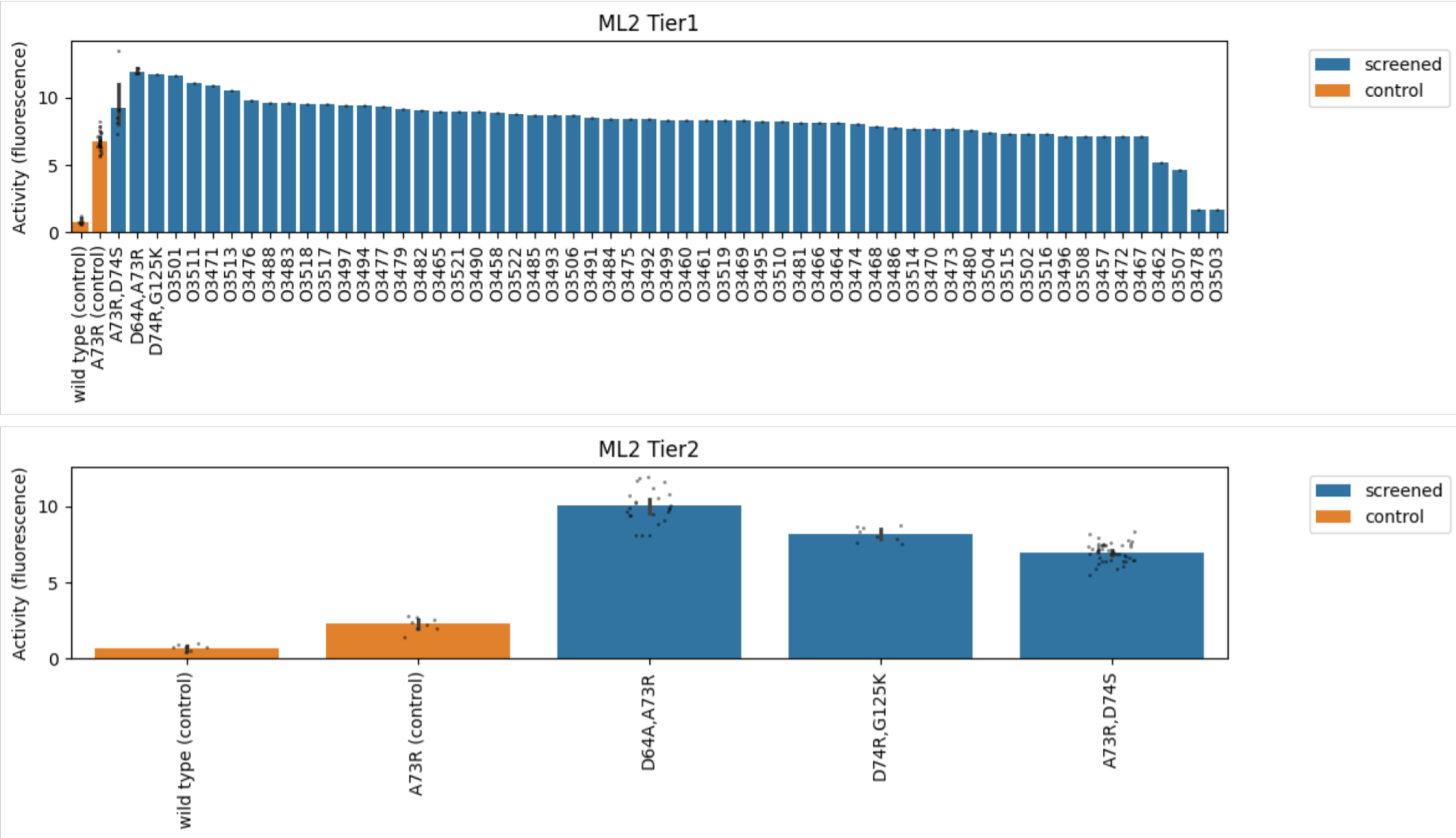
ML2 Tier 1 and 2 liquid culture plate assay activity assessment. The y axis (“activity”) measures fluorescence normalized by optical density. On the x axis are genotyped strains. Variants in orange were added to plates as controls. In Tier 1, variants that were not selected were not genotyped and are reported with their numerical strain identifier starting with “O”. D64A,A73R and A73R,D74S were selected for purification. Error bars represent ±1 standard error of the estimated activity.

**Supplementary Figure S12:**
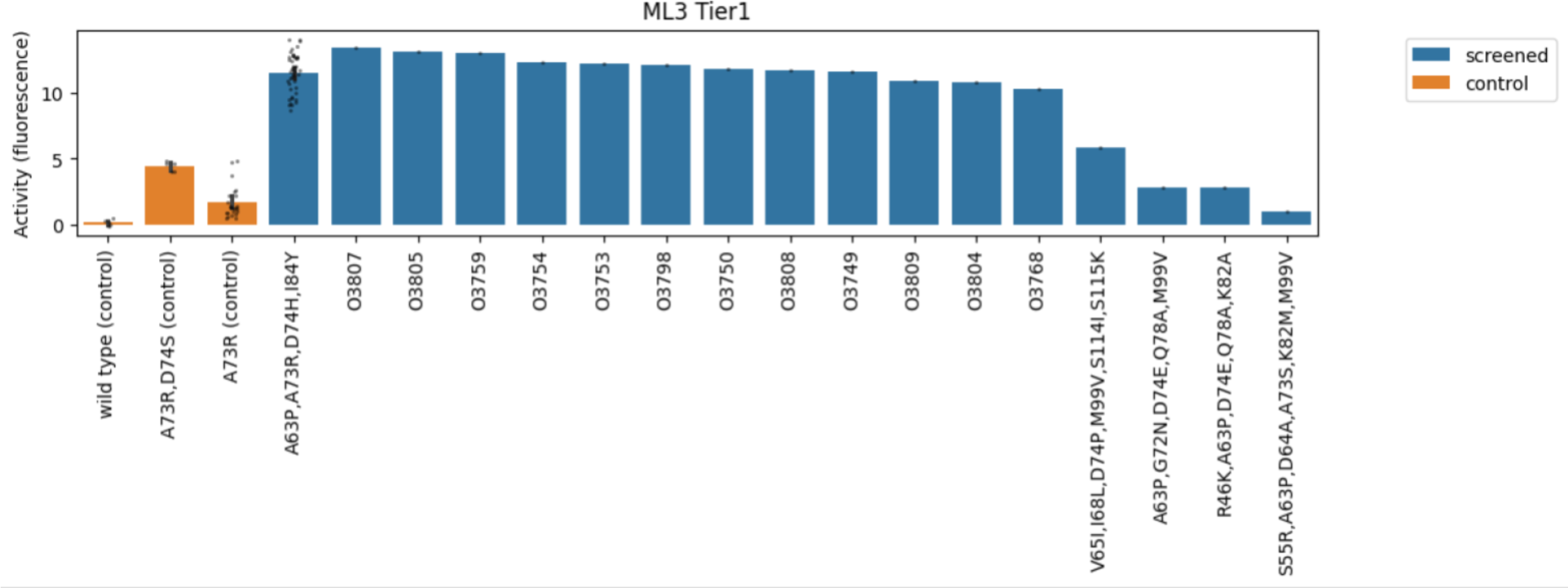
ML3 Tier 1 liquid culture plate assay activity assessment. The y axis (“activity”) measures fluorescence normalized by optical density. On the x axis are genotyped strains. Variants in orange were added to plates as controls. In Tier 1, variants that were not selected were not genotyped and are reported with their numerical strain identifier starting with “O”. A63P,A73R,D74H,I84Y was selected for purification. A Tier 2 screen was unnecessary because there were sufficient replicates of the highest performing genotype in Tier 1. Error bars represent ±1 standard error of the estimated activity.

**Supplementary Figure S13:**
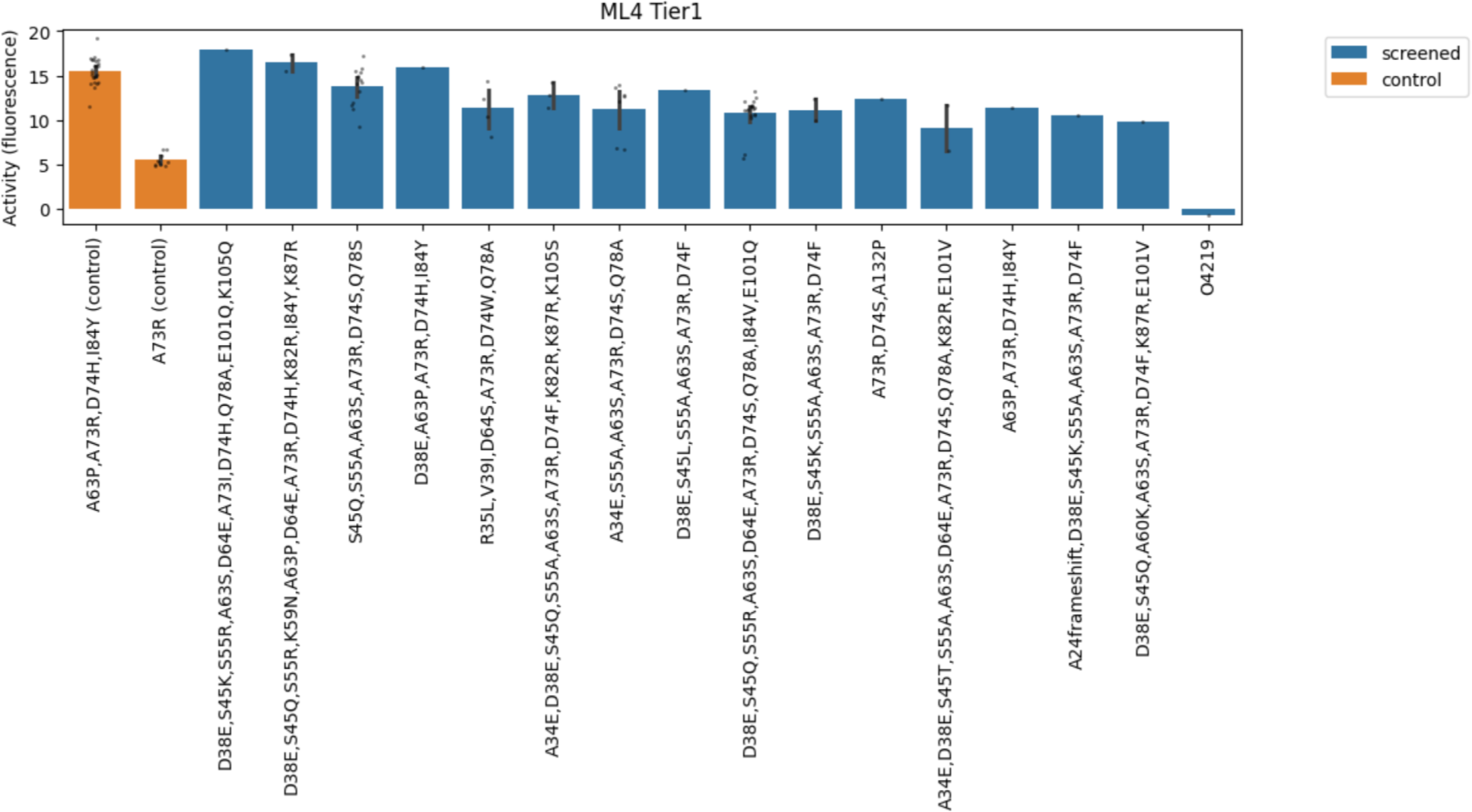
ML4 Tier 1 liquid culture plate assay activity assessment. The y axis (“activity”) measures fluorescence normalized by optical density. On the x axis are genotyped strains. Variants in orange were added to plates as controls. No variants were purified from this screen, but the screen indicates that 2 variants, one with 10 mutations and one with 11 mutations, achieved activity at or above that of the highest performing variant in ML3 (A63P, A73R, D74H, I84Y, orange).

**Supplementary Figure S14:**
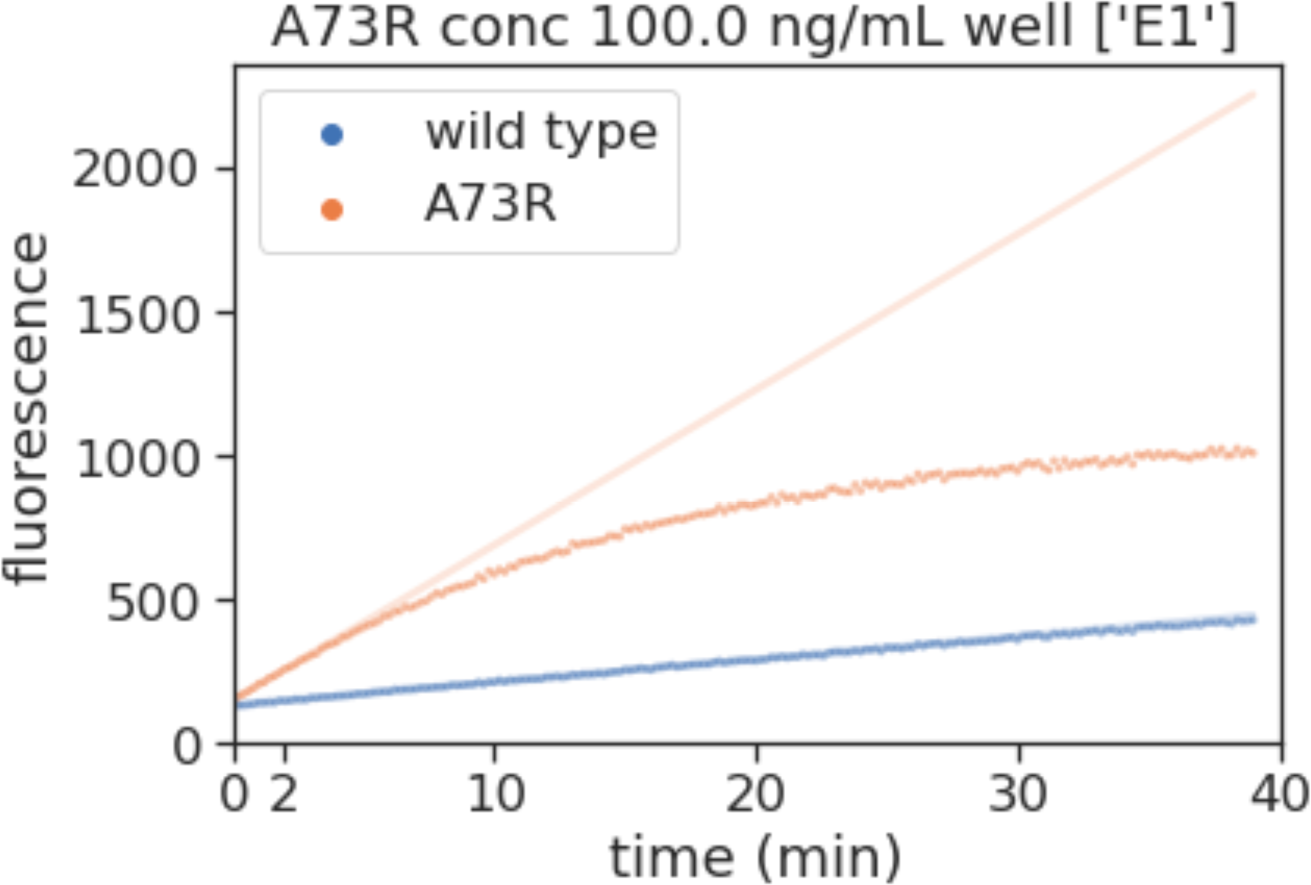
fluorescence signal over time for a reaction trajectory. The estimated slope is the initial velocity of the reaction. The slope of the variant (orange) was computed using the first 2 minutes of reaction data (10 points), while the slope of the wildtype (blue) was computed using 20 minutes of reaction trajectory (100 points). The ratio of the orange slope to the blue slope is the fold-change activity of the variant. The fold change was computed across 4 different concentrations (12.5, 25, 50, 100 ng/mL). In Figure 3b, the overall fold-change activity reported was the average across 1000 samples from the posterior of the estimated slope parameter in each of the 4 concentrations.

**Supplementary Figure S15:**
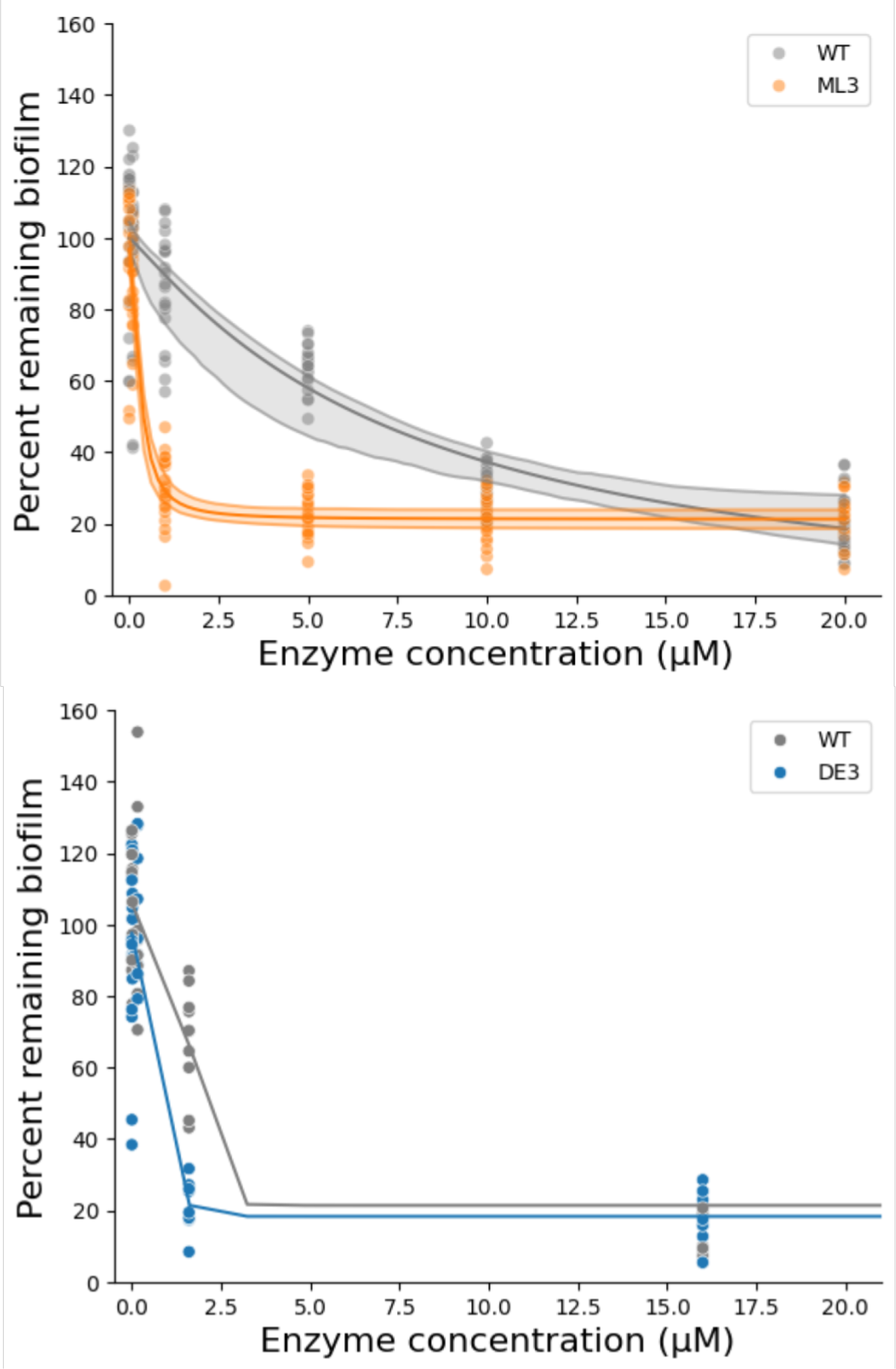
Dose-response curves for the best variant discovered in ML3 and DE3. The line of best fit was computed by the empirical fit of the observed data to the Hill equation using the least-squares method. The confidence bands represent the 5% and 95% percentiles of 200 resamples of the Hill equation parameters from the multivariate normal distribution inferred by scipy.optimize.curve_fit. We show the best fitting curve without bands for the DE3 (blue) best variant because there were insufficient replicates and insufficient tested conditions. There is clear improvement for biofilm degradation of the ML3 (orange) variant over the WT (gray) enzyme at low enzyme concentration. Similarly, there is clear improvement for biofilm degradation of the best DE3 variant over the WT at low enzyme concentration.

**Supplementary Figure S16:**
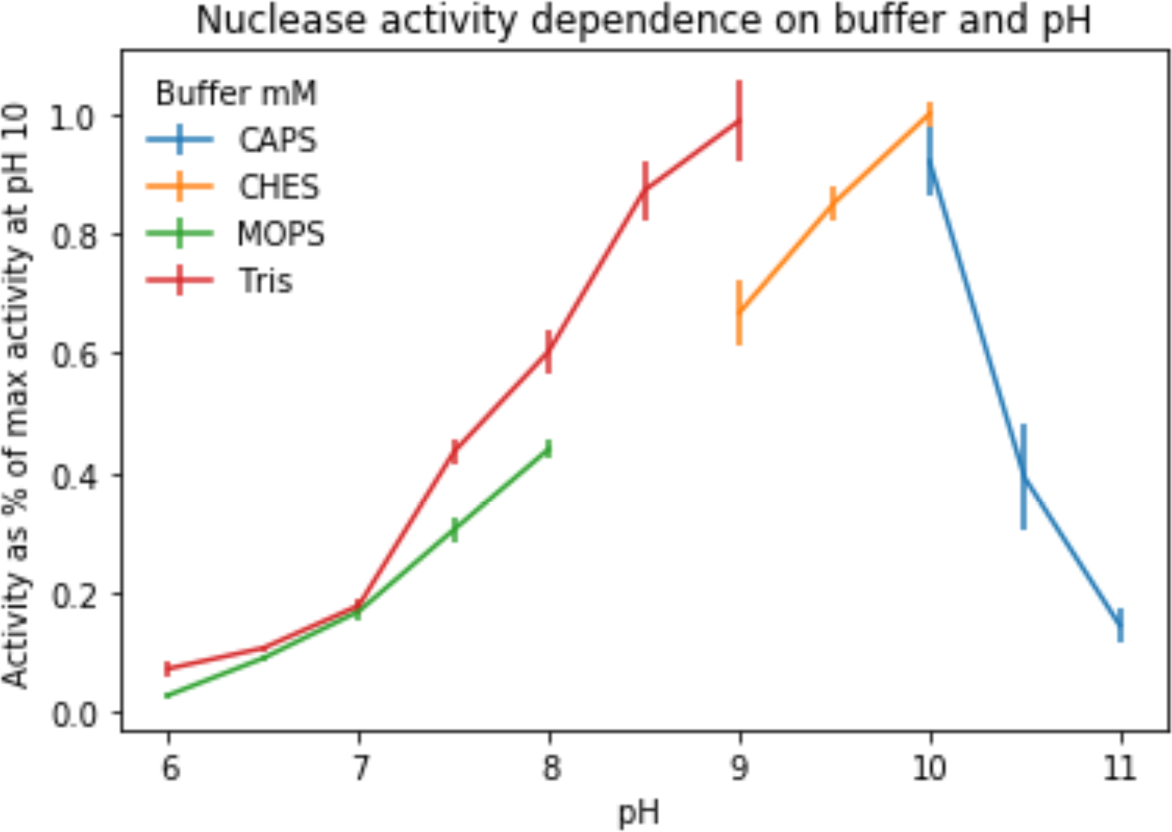
Wildtype NucB activity dependence on buffer and pH. The maximum enzyme activity was achieved in the CHES buffer at pH 10 (with 98.8% of the max activity achieved in the Tris buffer at pH 9). At pH 7, the activity is reduced: in MOPS the activity level is 16.9 ± 1.5% of the max activity, and in Tris, 17.8 ± 1.2%. Activity measurements vary depending on the buffer. Each point is an average of three technical replicates.

**Supplementary Figure S17:**
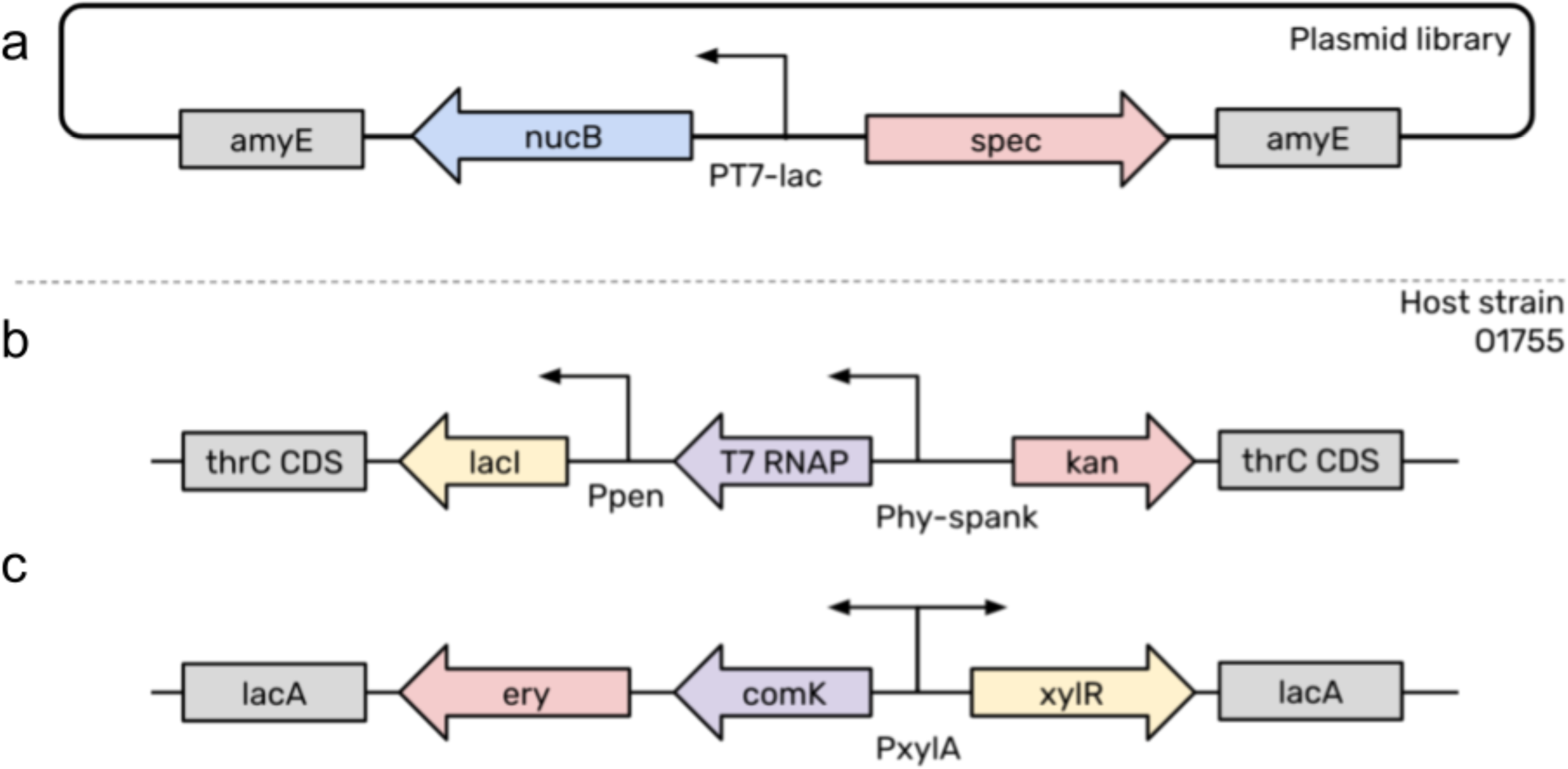
Diagram of plasmid library construct and host strain genetic modifications in the final expression strain. The expression strains consisted of the following: (a) the NucB library with unmodified native signal peptide under an IPTG-inducible T7 promoter integrated into the amyE locus with spectinomycin resistance; (b) LacI and an IPTG-inducible T7 RNA polymerase under the Phy-spank promoter integrated into the thrC locus with kanamycin resistance; (c) xylose-induced comK integrated into the lacA locus with erythromycin resistance.

**Supplementary Figure S18:**
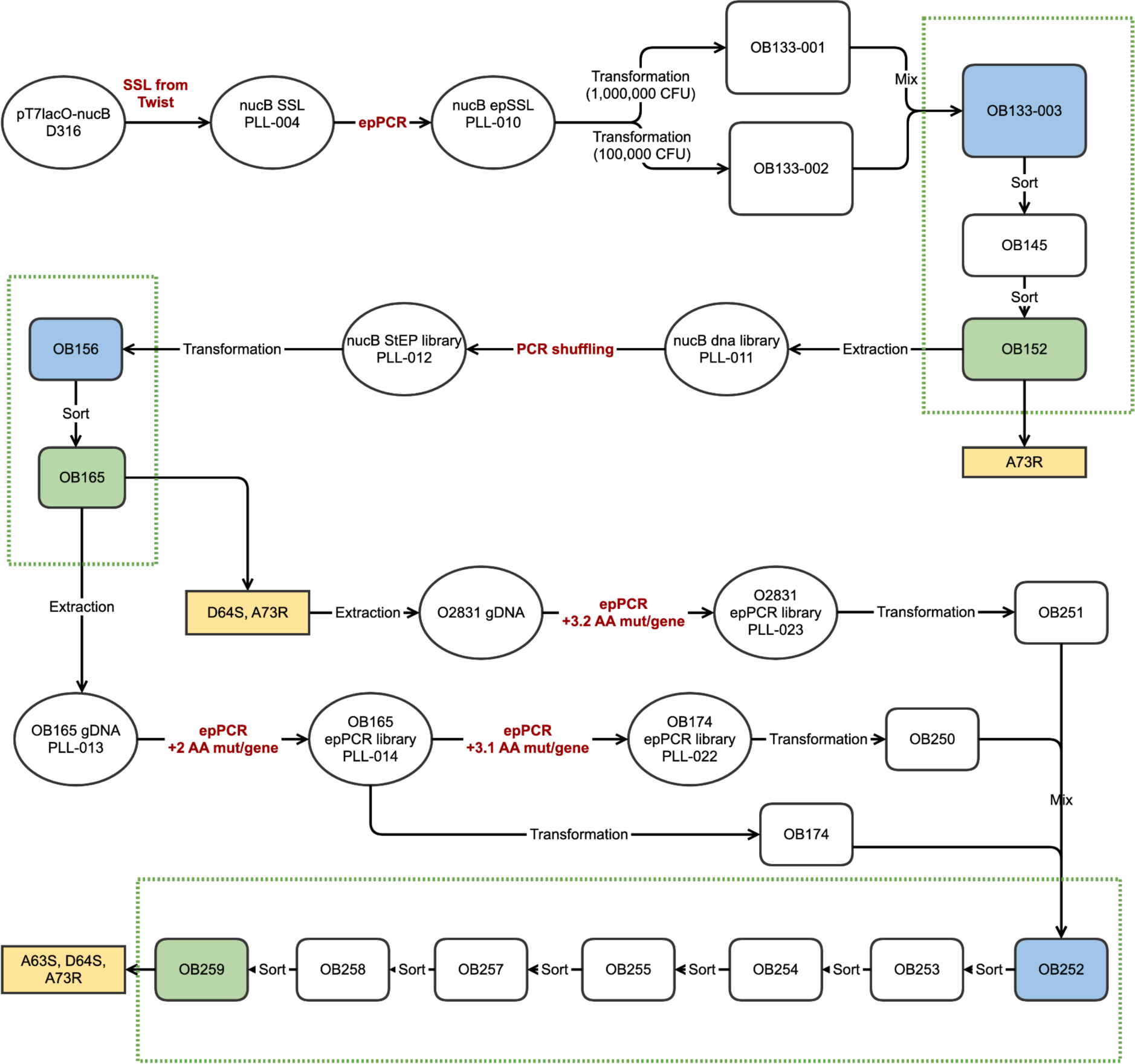
Schematic representation of the experimental workflow of the directed evolution (DE) campaign. The G1 library contained Site-Saturated Sequences (nucB SSL) with additional mutations added using error-prone PCR. For G1, one of the high-gate, post-sort libraries (OB145) was sorted again and underwent culture plate validation. The highest performing G1 strain A73R was found and sequenced from this library. In DE2, the sorted library (OB152) underwent recombination and was sorted again, resulting in a library (OB165) from which the highest performing DE2 strain D64S,A73R was found and sequenced. In DE3, three random libraries were mixed (OB252), one based on epPCR diversification of the highest performing DE2 strain, and the other based on epPCR diversification of an intermediate sorted library from DE2 (OB165). After successive sorts, the highest performing DE3 strain A63S,D64S,A73R was found and sequenced. All samples with names starting with OB represent libraries containing mixed strains. Each diversification step is highlighted in red. The starting library of each generation is colored blue and the ending library of each generation is colored in green. The highest activity genotype isolated from each generation is colored in yellow.

**Supplementary Figure S19:**
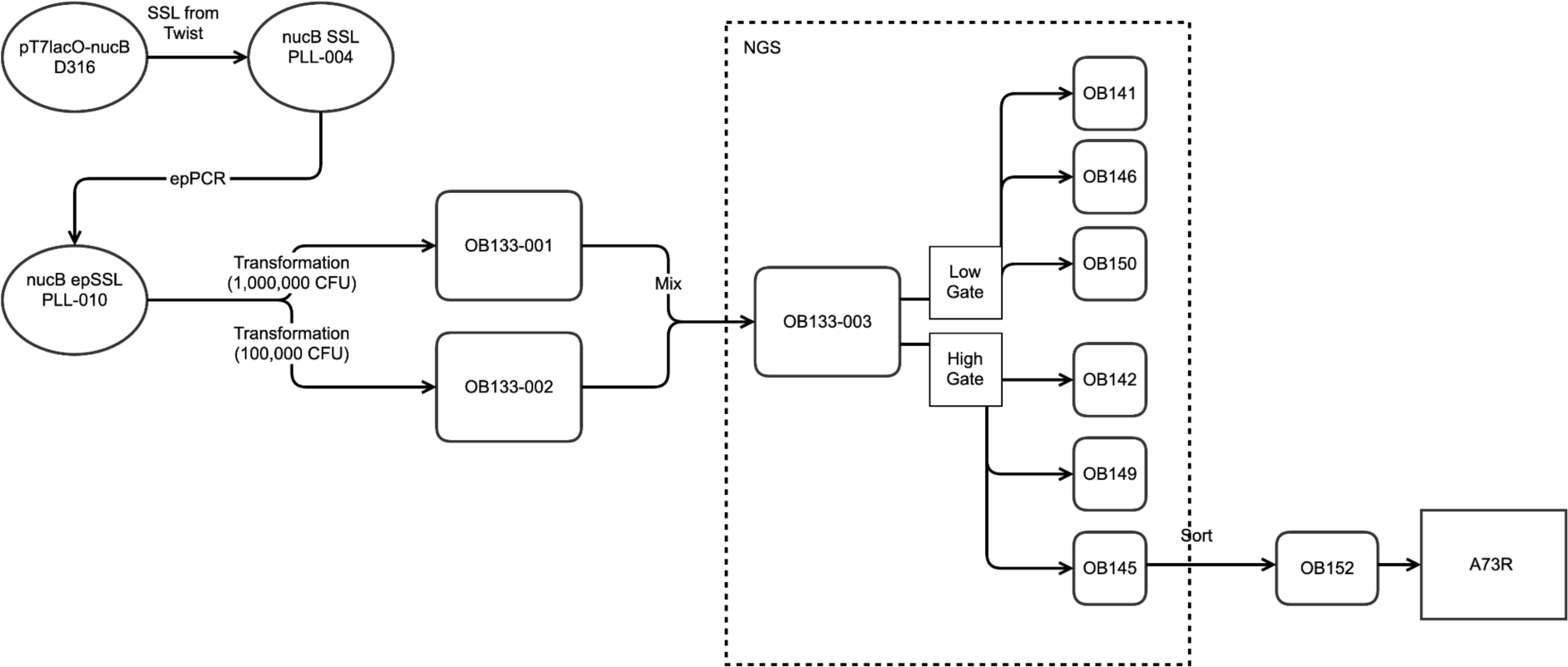
Schematic of G1. High and low gates were sorted and sequenced with NGS. The G1 library went through high and low gate sorting, with three replicates each. An additional sort (OB152) after a high gate sort led to the identification of the A73R variant. All libraries that were sequenced using NGS are enclosed in dotted lines.

**Supplementary Figure S20:**
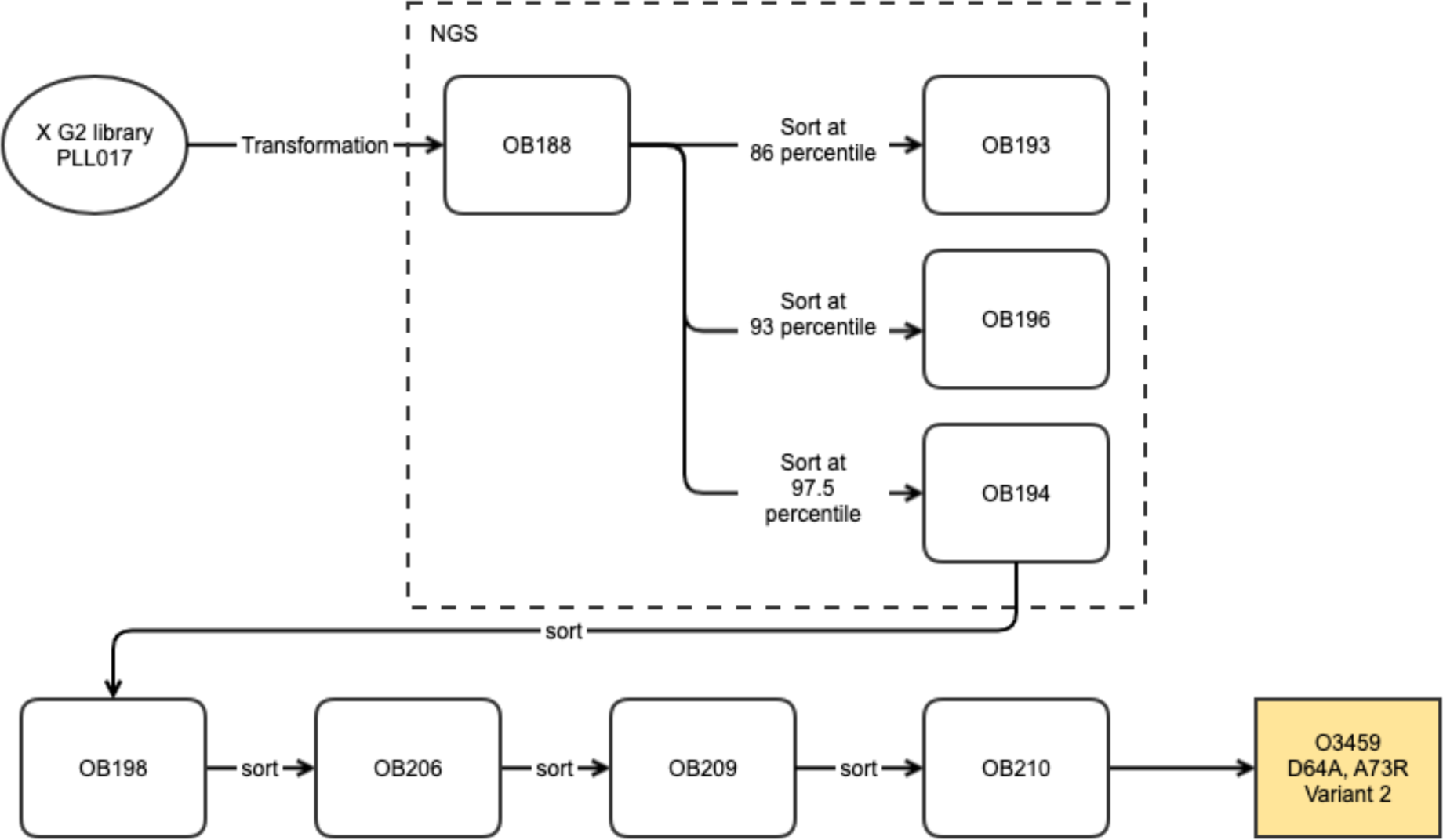
Schematic of G2. Three gates were used for data generation in G2, corresponding to different library fluorescence percentiles. An additional 4 sorts after a 97.5 percentile sort led to the identification of the D64A,A73R variant. All libraries that were sequenced using NGS are enclosed in dotted lines.

**Supplementary Figure S21:**
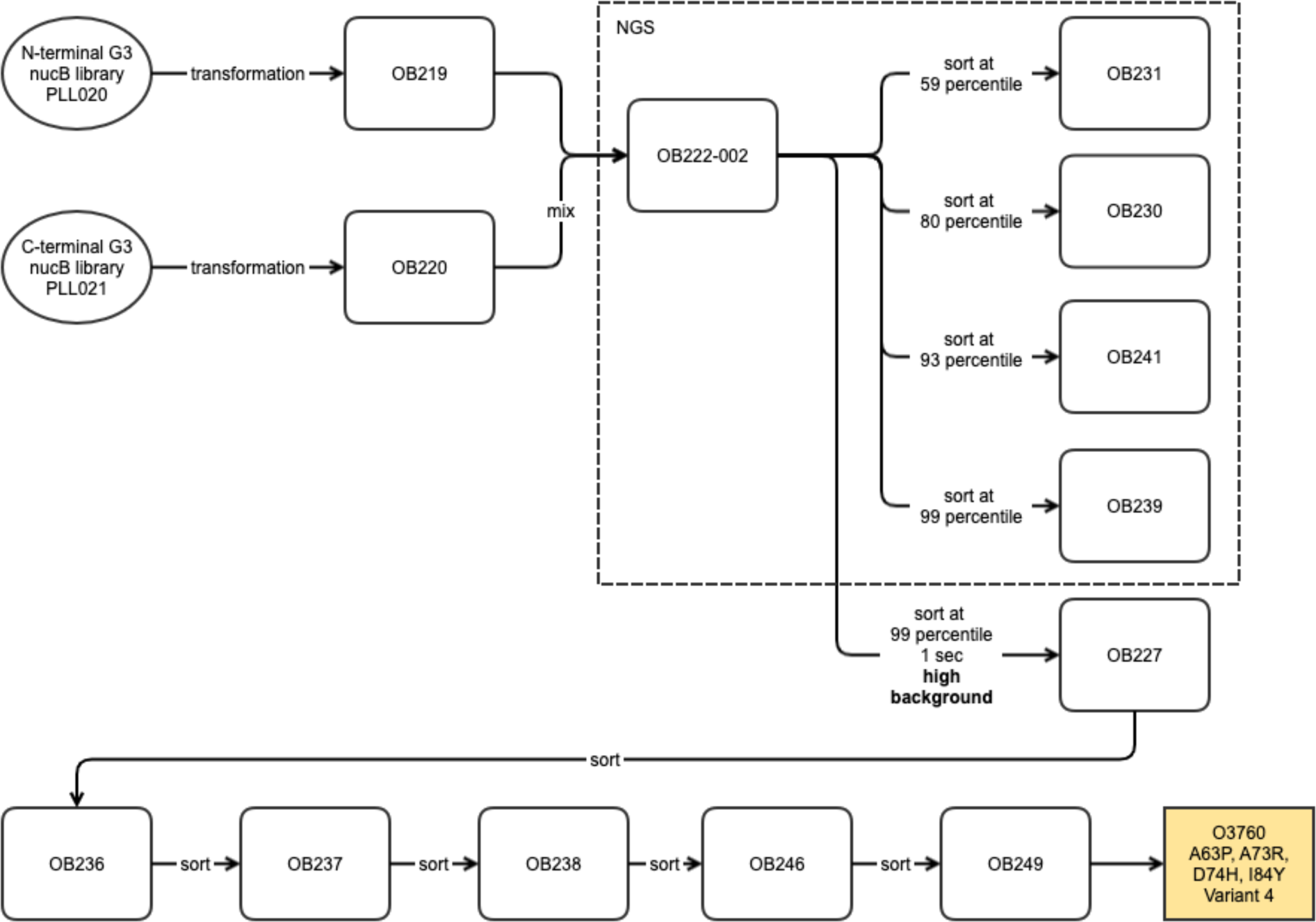
Schematic of G3. Four gates were used for data generation in G3, corresponding to different library fluorescence percentiles. An additional 6 sorts after a 99th percentile sort (not sequenced) led to the identification of the A63P,A73R,D74H,I84Y variant. All libraries that were sequenced using NGS are enclosed in dotted lines.

**Supplementary Figure S22:**
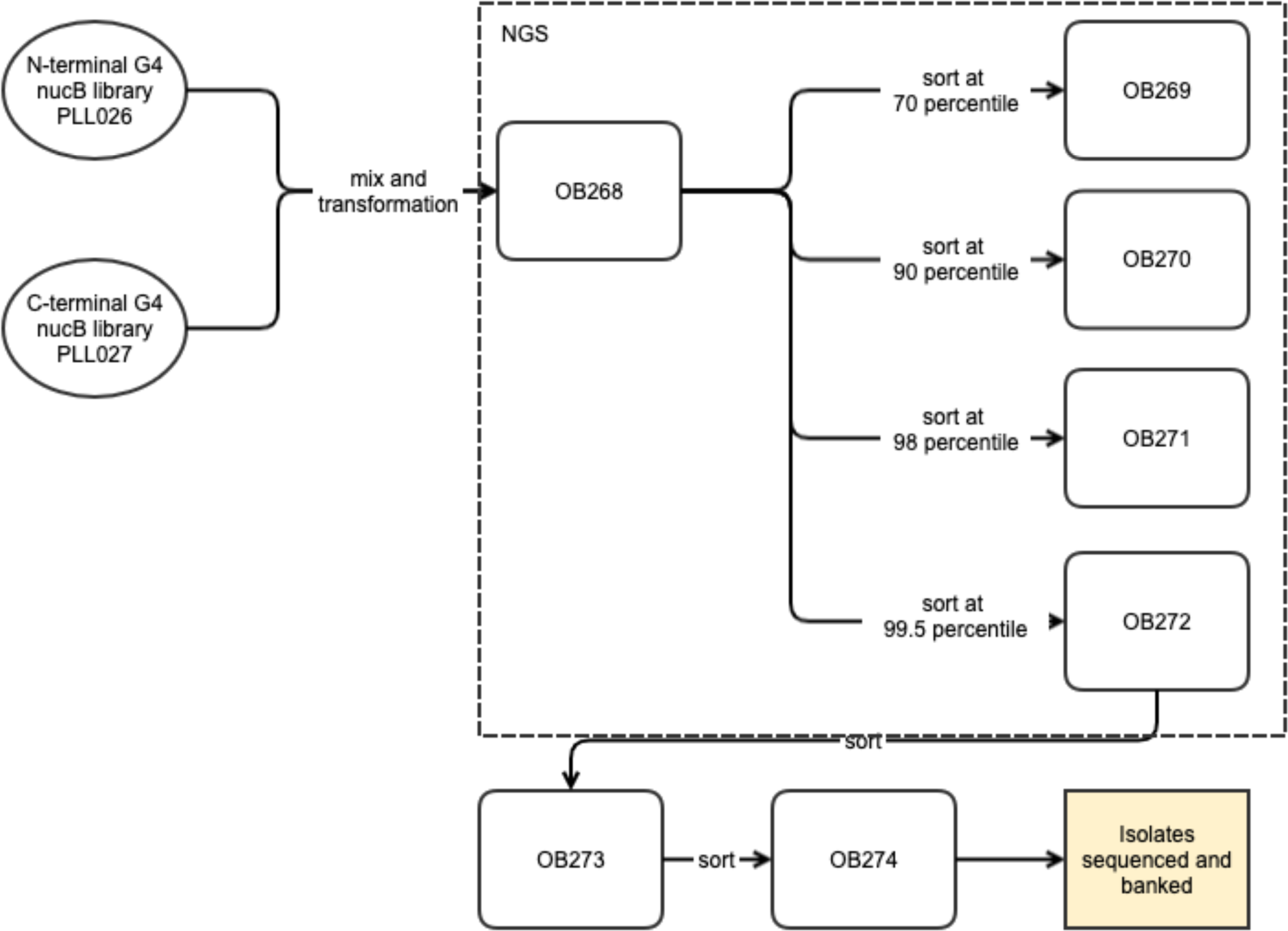
Schematic of G4. Four gates were used for data generation in G3, corresponding to different library fluorescence percentiles. All libraries that were sequenced using NGS are enclosed in dotted lines. The final isolates were sequenced and banked.

**Supplementary Figure S23:**
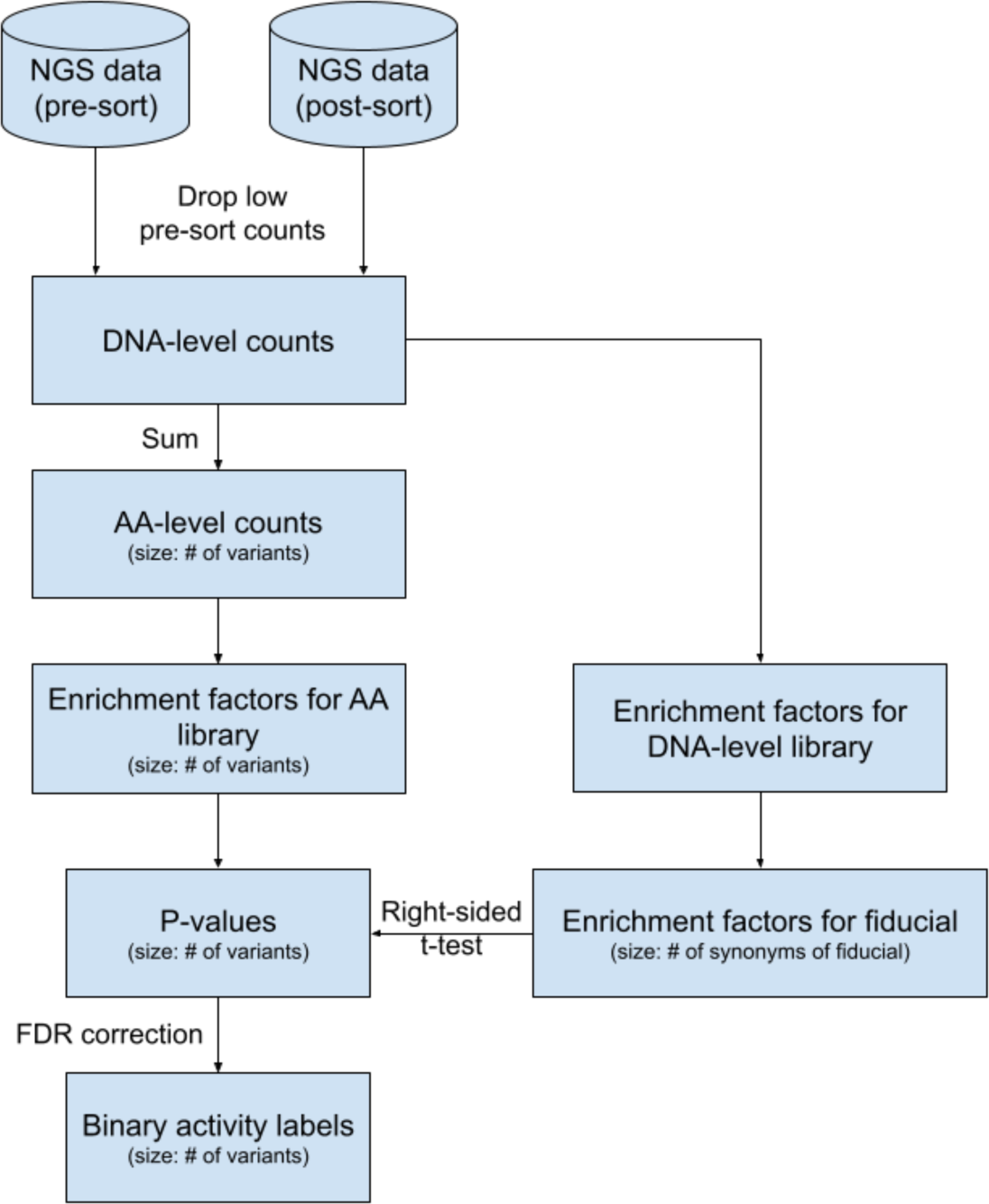
schematic of data processing. The per-variant activity labels are derived as follows: load all NGS counts for the screened library, drop variants with low input counts, compute enrichment factors at both the DNA and amino acid level, compute p-values for a right-sided t-test with the null distribution given by a fiducial’s DNA-level enrichment factors, and label variants as having higher activity than the fiducial sequence, using a false-discovery rate correction.

**Supplementary Figure S24:**
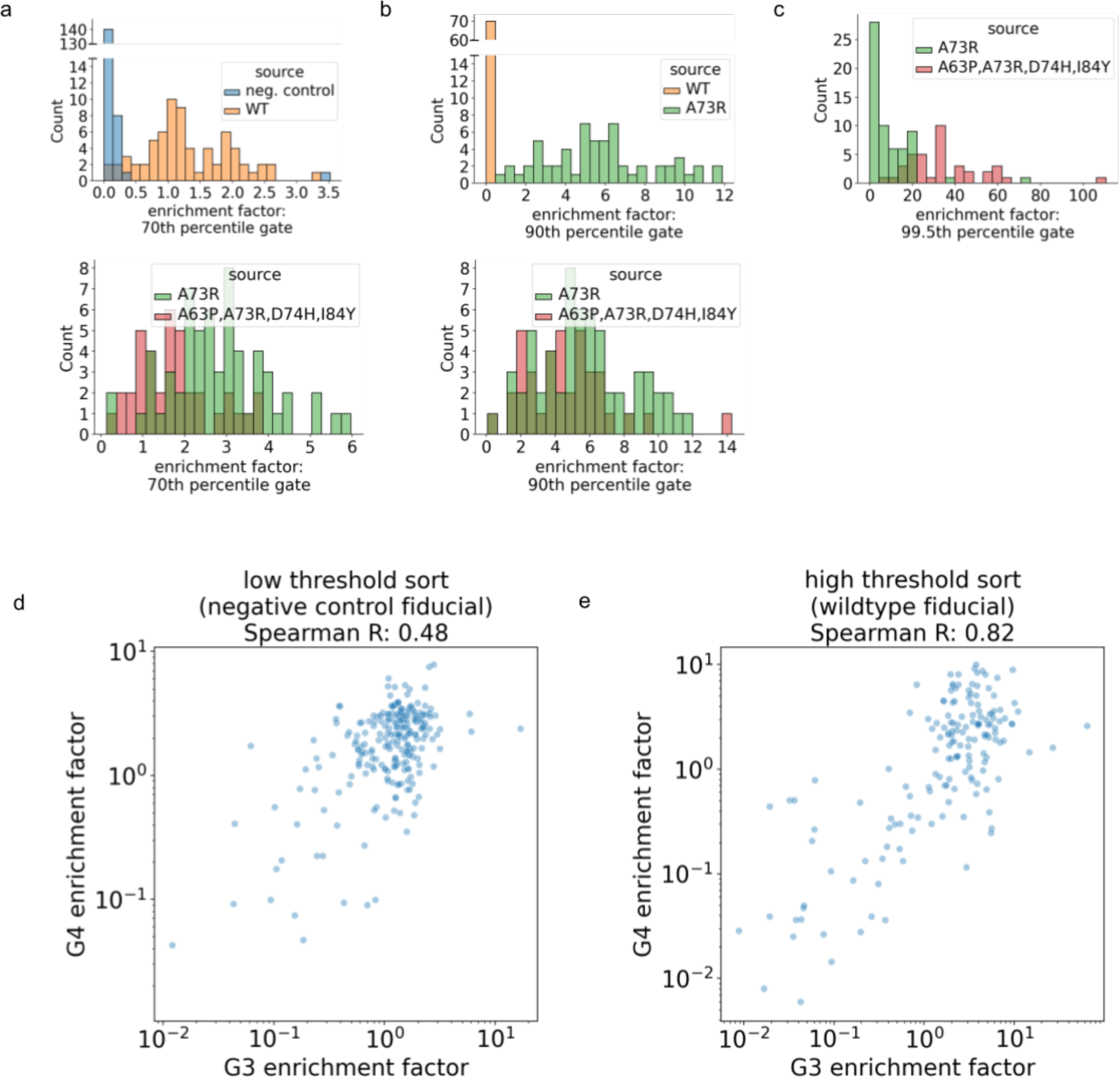
sorts with higher fluorescence thresholds separate fiducials by activity while low threshold gates exhibit saturation behavior. In (a), (b), and (c), the enrichment factor distribution of two fiducials is shown in a series of successively-higher gates used in G4, where the fluorescence threshold is lowest in (a) and highest in (c). Based on culture plate assays, we expect the activity of the fiducial sequences to be ordered by negative control < WT < A73R < quad, where “negative control” refers to stop codons introduced in the middle of the coding sequence, which should produce inactive proteins. We see that the gate used in (a) separates negative control and WT fiducials (top panel), the gate used in (b) separates WT and A73R fiducials (top panel). While the gate used in (c) separates A73R and quad (A63P, A73R, D74H, I84Y) fiducials (top panel), the lower threshold gates used in (a) and (b) are unable to separate the A73R and quad fiducials (bottom panel), as most replicates of high activity variants pass the lower threshold screen. In (d) and (e), we plot the enrichment factors for 251 functional variants screened in both G4 and G3. The G4 enrichment factor (y axis) and G3 enrichment factor (x axis) are plotted for the sorts used to determine activity level relative to negative control (d) and wildtype (e) fiducials, analogous to the negative control sort in subfigure (a) and wildtype sort in subfigure (b), respectively. The variants screened in both G3 and G4 were enriched for functional variants, and as such, we see enrichment factor saturation in the low threshold sorts (d), leading to a lower enrichment factor correlation across generations than in the high threshold sorts (e) (Spearman R 0.48 vs 0.82). However, our discrete activity labels were concordant across generations. Of the variants sorted with a threshold calibrated to the negative control fiducial (d), 206 were concordantly predicted to be better than the negative control in both G3 and G4, 24 were concordantly predicted to be worse than the negative control, and 21 were predicted discordantly. Of the variants sorted with a threshold calibrated to the wildtype fiducial, 131 were predicted to be better than wildtype in both G3 and G4, 96 were predicted to be worse than wildtype in both rounds, and 24 were predicted discordantly.

**Supplementary Figure S25:**
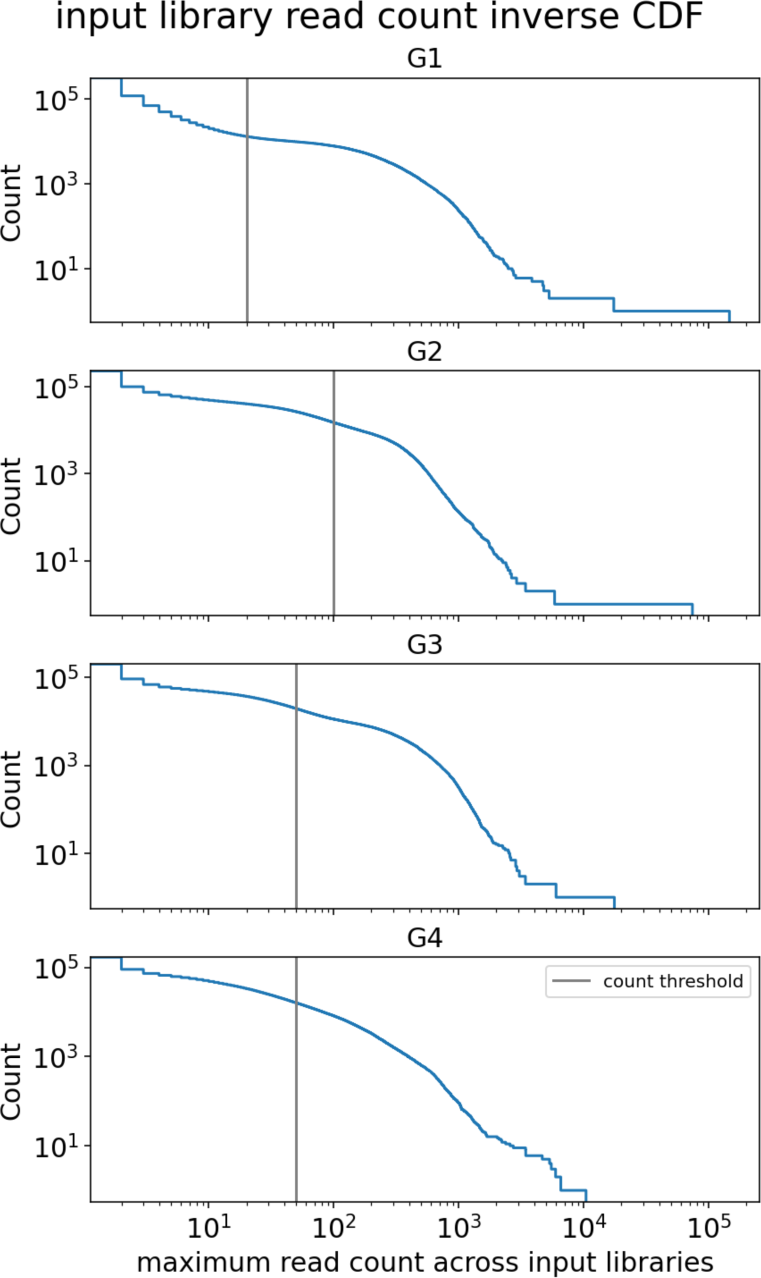
Read count thresholding determines the number of unique variants we can characterize. Read count thresholds were determined heuristically in each generation to assure accurate enrichment factor estimates. In order from G1 to G4 (top to bottom), the read count threshold was set to 20, 100, 50, and 50 reads, respectively. In each generation, this corresponds to filtering the library from ∼10^5^ (no thresholding) to ∼10^4^ variants (with thresholding).

**Supplementary Figure S26:**
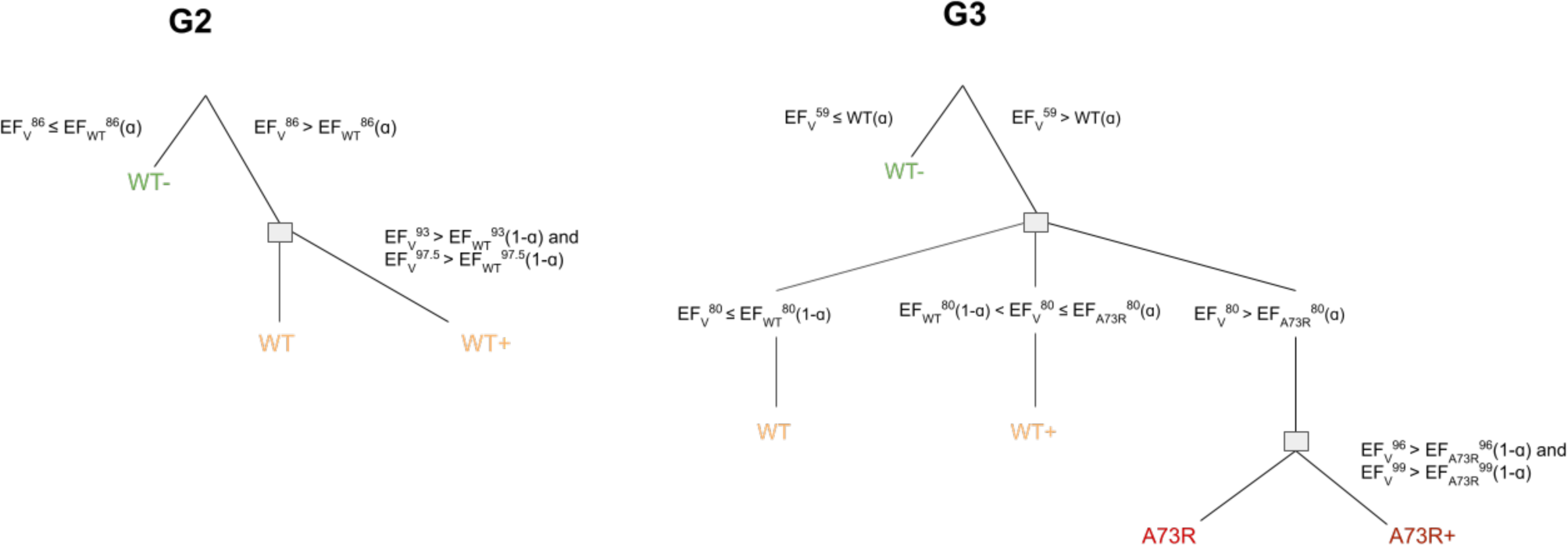
Procedure for assigning discrete ML training labels for G2 and G3 data. Activity labels were assigned through a series of comparisons, utilizing all sorts for which there was data. The variant V enrichment factor in a given sort is indicated by the library percentile of the sort (EF_V_^sort percentile^). Edges are labeled with the comparison used to make an assignment. EF_fiducial_^sort^(α) (e.g. EF_A73R_^80^(α)) and refers to the α quantile of the empirical synonym enrichment factor distribution for the fiducial. The final label is in the set {WT-, WT, WT+, A73R, A73R+} and was used as a classification objective for model training.

**Supplementary Figure S27:**
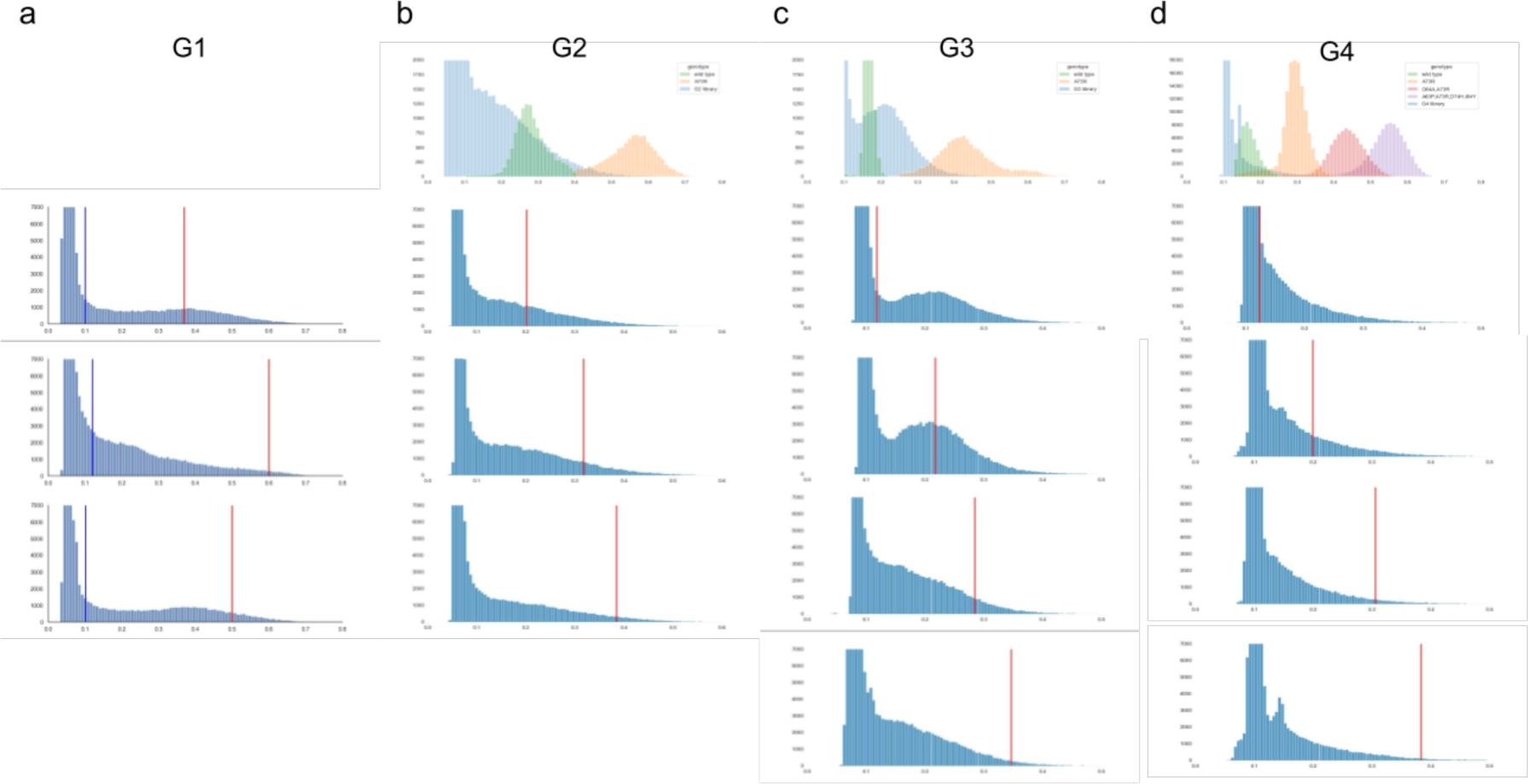
Gate fluorescence thresholds compared to fiducial sequence fluorescence for all generations. Columns a, b, c, and d correspond to screens of G1, G2, G3 and G4, respectively. The histogram in the first row show the overall library fluorescence distribution (blue) and the fiducial fluorescence distributions (WT in green, A73R in orange, D64A,A73R in red, and A63P,A73R,D74H,I84Y in purple). G1 was a random library, and does not include fiducials. The histograms in each subsequent row show the fluorescence level for each sort in that generation. The red line indicates the minimum fluorescence threshold for a cell to be sorted. In column a the low gate sort threshold is additionally indicated by the blue line, while the high gate sort threshold is indicated by the red line.

**Supplementary Figure S28:**
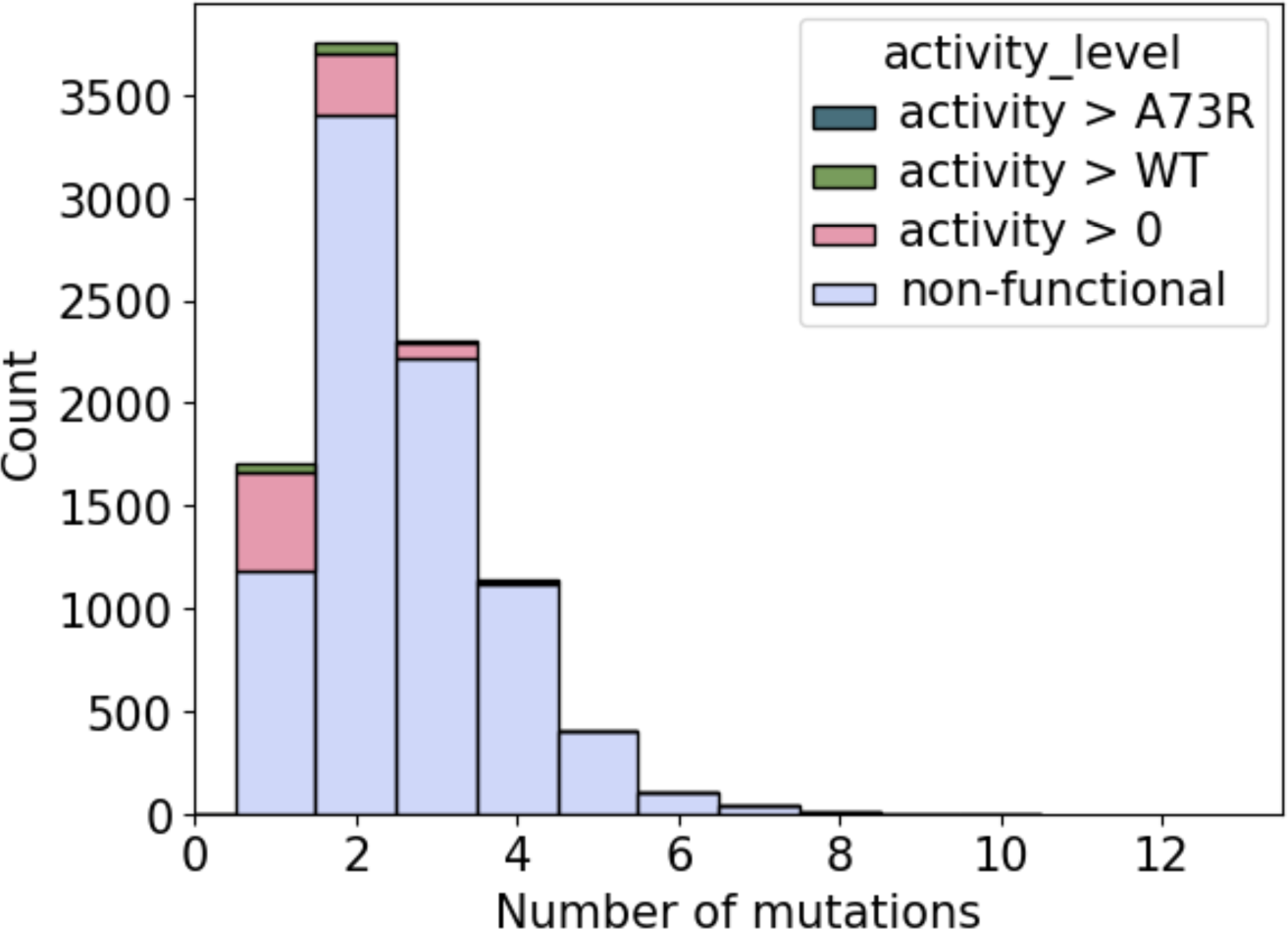
Activity level distribution for G1. G1 variants, colored by estimated activity levels. G1 included a single-site saturation mutagenesis library as well as variants generated by error-prone PCR. On average, G1 variants had 2.5 mutations, while functional variants had, on average, 1.6 mutations.

**Supplementary Table S1:** hit rates of all sub-libraries for both the WT and A73R fiducials. In each generation, the non-control cell with the highest hit-rate at each activity level threshold is bolded. Control sub-libraries are denoted separately, and were included in parallel with each generation’s library, but are not compared to MLDE techniques. Note that the zero-shot library (an alternative to G1) was screened in parallel with G4. Similarly, the “prosar+screen_g2_redux” library (an alternative to G2) was screened in parallel with G4. The zero-shot library was screened in parallel with G4, but is displayed next to G1. The total number of hits here differs slightly from the activity labels in our final genotype-phenotype fitness landscape file. This is because we applied a Benjamini-Hochberg false discovery rate correction within each row in this table, whereas it was applied globally when producing the landscape.

| generation | name | library size | num hits (> neg_control) | hit rate % (> neg_control) | num hits (> WT) | hit rate % (> WT) | num hits (> A73R) | hit rate % (> A73R) |
| --- | --- | --- | --- | --- | --- | --- | --- | --- |
| g1 | g1_eppcr | 9441 | 1084.5 +/- 33.0 | 11.49 +/- 0.35 | 155.1 +/- 10.0 | 1.64 +/- 0.11 | N/A | N/A |
| g1 (alt) | zero_shot | 1235 | 629.0 +/- 22.3 | **50.93 +/- 1.81** | 134.8 +/- 10.8 | **10.91 +/- 0.87** | 0.0 +/- 0.0 | 0.00 +/- 0.00 |
|  | G2 |  |  |  |  |  |  |  |
| g2 | g2_hit_recombination | 951 | 830.9 +/- 12.5 | **87.37 +/- 1.32** | 470.0 +/- 10.1 | **49.42 +/- 1.06** | N/A | N/A |
| g2 | g2_broad_recombination_sampling | 3598 | 99.5 +/- 9.5 | 2.76 +/- 0.26 | 6.1 +/- 3.5 | 0.17 +/- 0.10 | N/A | N/A |
| g2 | g2_mbo_dnn_exploit | 287 | 104.5 +/- 9.7 | 36.40 +/- 3.37 | 43.1 +/- 7.5 | 15.03 +/- 2.63 | N/A | N/A |
| g2 | g2_mbo_dnn_explore | 4083 | 1393.9 +/- 27.3 | 34.14 +/- 0.67 | 171.7 +/- 20.0 | 4.20 +/- 0.49 | N/A | N/A |
| g2 (alt) | prosar+screen_g2_redux | 1464 | 1034.5 +/- 15.3 | 70.66 +/- 1.04 | 559.3 +/- 19.5 | 38.21 +/- 1.33 | 0.0 +/- 0.0 | 0.00 +/- 0.00 |
|  | G2 Controls |  |  |  |  |  |  |  |
| g2 | g2_g1_plate_assay_variants | 5 | 5.0 +/- 0.0 | 100.00 +/- 0.00 | 3.7 +/- 0.9 | 73.33 +/- 17.99 | N/A | N/A |
| g2 | g2_single_mutants | 1402 | 384.4 +/- 12.7 | 27.42 +/- 0.91 | 43.0 +/- 6.3 | 3.07 +/- 0.45 | N/A | N/A |
| g2 | g2_stratified_sample | 40 | 16.2 +/- 3.6 | 40.50 +/- 9.12 | 2.7 +/- 1.7 | 6.83 +/- 4.27 | N/A | N/A |
| g2 | g2_unmatched | 4365 | 446.8 +/- 18.3 | 10.24 +/- 0.42 | 103.9 +/- 11.8 | 2.38 +/- 0.27 | N/A | N/A |
| g2 | g2_wt_synonyms | 1 | 1.0 +/- 0.0 | 100.00 +/- 0.00 | 0.0 +/- 0.0 | 0.00 +/- 0.00 | N/A | N/A |
|  | G3 |  |  |  |  |  |  |  |
| g3 | g3_hit_recombination | 1712 | 1443.9 +/- 19.0 | 84.34 +/- 1.11 | 815.8 +/- 21.3 | 47.65 +/- 1.24 | 26.8 +/- 7.7 | 1.57 +/- 0.45 |
| g3 | g3_mbo_dnn_exploit | 369 | 277.3 +/- 9.2 | 75.16 +/- 2.51 | 196.3 +/- 11.1 | 53.21 +/- 3.01 | 11.2 +/- 3.6 | 3.04 +/- 0.97 |
| g3 | g3_mbo_dnn_explore | 299 | 233.4 +/- 6.9 | 78.06 +/- 2.32 | 174.3 +/- 8.4 | 58.28 +/- 2.80 | 7.7 +/- 3.0 | 2.59 +/- 1.00 |
| g3 | g3_prosar_high_screen_high | 670 | 634.7 +/- 4.5 | **94.74 +/- 0.68** | 472.3 +/- 9.7 | **70.49 +/- 1.44** | 23.2 +/- 6.2 | **3.46 +/- 0.92** |
| g3 | g3_prosar_high_screen_low | 2080 | 1690.9 +/- 9.6 | 81.29 +/- 0.46 | 979.7 +/- 24.7 | 47.10 +/- 1.19 | 20.2 +/- 7.3 | 0.97 +/- 0.35 |
| g3 | g3_prosar_high_unscreened | 1315 | 766.9 +/- 16.1 | 58.32 +/- 1.23 | 362.1 +/- 16.2 | 27.54 +/- 1.23 | 1.0 +/- 2.0 | 0.08 +/- 0.15 |
| g3 | g3_prosar_low_screen_high | 820 | 765.9 +/- 5.4 | 93.40 +/- 0.66 | 345.1 +/- 17.1 | 42.08 +/- 2.09 | 3.5 +/- 1.6 | 0.43 +/- 0.19 |
| g3 | g3_prosar_low_screen_low | 1930 | 1568.3 +/- 18.1 | 81.26 +/- 0.94 | 707.5 +/- 20.7 | 36.66 +/- 1.07 | 7.6 +/- 3.3 | 0.39 +/- 0.17 |
| g3 | g3_prosar_low_unscreened | 1108 | 711.3 +/- 14.7 | 64.20 +/- 1.32 | 245.9 +/- 14.0 | 22.20 +/- 1.27 | 4.5 +/- 2.3 | 0.40 +/- 0.21 |
|  | G3 Controls |  |  |  |  |  |  |  |
| g3 | g3_a73r_synonyms | 1 | 1.0 +/- 0.0 | 100.00 +/- 0.00 | 1.0 +/- 0.0 | 100.00 +/- 0.00 | 0.0 +/- 0.0 | 0.00 +/- 0.00 |
| g3 | g3_g1_stratified_sample | 38 | 12.9 +/- 3.0 | 33.86 +/- 7.89 | 2.1 +/- 1.9 | 5.44 +/- 4.92 | 0.0 +/- 0.0 | 0.00 +/- 0.00 |
| g3 | g3_g2_plate_assay_variants | 6 | 6.0 +/- 0.0 | 100.00 +/- 0.00 | 4.9 +/- 0.8 | 81.11 +/- 13.90 | 0.0 +/- 0.0 | 0.00 +/- 0.00 |
| g3 | g3_loop_explore | 389 | 123.1 +/- 11.3 | 31.64 +/- 2.91 | 2.3 +/- 1.5 | 0.58 +/- 0.38 | 0.0 +/- 0.0 | 0.00 +/- 0.00 |
| g3 | g3_single_mutants | 577 | 221.1 +/- 9.6 | 38.32 +/- 1.66 | 4.1 +/- 2.2 | 0.70 +/- 0.38 | 0.0 +/- 0.0 | 0.00 +/- 0.00 |
| g3 | g3_wt_synonyms | 1 | 1.0 +/- 0.0 | 100.00 +/- 0.00 | 0.0 +/- 0.0 | 0.00 +/- 0.00 | 0.0 +/- 0.0 | 0.00 +/- 0.00 |
| g3 | g3_unmatched | 7796 | 3831.5 +/- 38.9 | 49.15 +/- 0.50 | 1853.0 +/- 36.8 | 23.77 +/- 0.47 | 76.3 +/- 13.0 | 0.98 +/- 0.17 |
|  | G4 |  |  |  |  |  |  |  |
| g4 | g4_hit_recombination | 1540 | 1009.6 +/- 22.7 | 65.56 +/- 1.47 | 886.9 +/- 14.1 | 57.59 +/- 0.91 | 8.5 +/- 4.2 | 0.55 +/- 0.28 |
| g4 | g4_mbo_dnn | 1356 | 1095.5 +/- 15.4 | **80.79 +/- 1.14** | 1147.0 +/- 15.8 | **84.59 +/- 1.17** | 52.4 +/- 6.8 | **3.86 +/- 0.50** |
| g4 | g4_mbo_linear | 280 | 2.1 +/- 1.3 | 0.76 +/- 0.47 | 13.7 +/- 3.5 | 4.90 +/- 1.24 | 0.0 +/- 0.0 | 0.00 +/- 0.00 |
| g4 | g4_sample_and_screen_dnn | 1298 | 977.8 +/- 14.1 | 75.33 +/- 1.09 | 898.5 +/- 17.2 | 69.22 +/- 1.32 | 1.9 +/- 1.3 | 0.15 +/- 0.10 |
| g4 | g4_sample_and_screen_linear | 568 | 186.1 +/- 12.8 | 32.77 +/- 2.25 | 114.1 +/- 8.0 | 20.08 +/- 1.40 | 0.0 +/- 0.0 | 0.00 +/- 0.00 |
| g4 | g4_sample_unscreened | 592 | 161.3 +/- 12.8 | 27.24 +/- 2.16 | 113.3 +/- 11.6 | 19.13 +/- 1.97 | 0.0 +/- 0.0 | 0.00 +/- 0.00 |
|  | G4 Controls |  |  |  |  |  |  |  |
| g4 | g4_a73r_synonyms | 1 | 1.0 +/- 0.0 | 100.00 +/- 0.00 | 1.0 +/- 0.0 | 100.00 +/- 0.00 | 0.0 +/- 0.0 | 0.00 +/- 0.00 |
| g4 | g4_double_synonyms | 1 | 1.0 +/- 0.0 | 100.00 +/- 0.00 | 1.0 +/- 0.0 | 100.00 +/- 0.00 | 0.0 +/- 0.0 | 0.00 +/- 0.00 |
| g4 | g4_g3_hit_constituents | 241 | 234.1 +/- 2.0 | 97.15 +/- 0.81 | 209.1 +/- 4.2 | 86.78 +/- 1.74 | 1.4 +/- 1.1 | 0.58 +/- 0.44 |
| g4 | g4_g3_plate_assay_variants | 14 | 14.0 +/- 0.0 | 100.00 +/- 0.00 | 14.0 +/- 0.0 | 100.00 +/- 0.00 | 0.4 +/- 1.1 | 2.86 +/- 7.54 |
| g4 | g4_homolog_graft | 11 | 5.6 +/- 1.8 | 50.91 +/- 16.76 | 0.0 +/- 0.0 | 0.00 +/- 0.00 | 0.0 +/- 0.0 | 0.00 +/- 0.00 |
| g4 | g4_mbo_seeds | 27 | 27.0 +/- 0.0 | 100.00 +/- 0.00 | 26.0 +/- 0.8 | 96.30 +/- 3.13 | 0.4 +/- 0.7 | 1.48 +/- 2.73 |
| g4 | g4_quad_synonyms | 1 | 1.0 +/- 0.0 | 100.00 +/- 0.00 | 1.0 +/- 0.0 | 100.00 +/- 0.00 | 1.0 +/- 0.0 | 100.00 +/- 0.00 |
| g4 | g4_wt_synonyms | 1 | 1.0 +/- 0.0 | 100.00 +/- 0.00 | 0.0 +/- 0.0 | 0.00 +/- 0.00 | 0.0 +/- 0.0 | 0.00 +/- 0.00 |
| g4 | g4_other | 4494 | 1147.9 +/- 40.7 | 25.54 +/- 0.91 | 944.9 +/- 28.6 | 21.03 +/- 0.64 | 0.7 +/- 1.1 | 0.02 +/- 0.02 |
| g4 | g4_unmatched | 2360 | 425.4 +/- 20.1 | 18.03 +/- 0.85 | 330.3 +/- 16.0 | 14.00 +/- 0.68 | 1.9 +/- 2.3 | 0.08 +/- 0.10 |

**Supplementary Table S2.** hit rates for G4 libraries for the A73R,D73S fiducial. . For brevity, only sub-libraries with non-zero hit rate are included. Note g4_quad_synonyms contains synonyms for a high-performance fiducial, so its hit rate is high by construction.

| generation | name | library size | num hits (> A73R,D74S) | hit rate % (> A73R,D74S) |
| --- | --- | --- | --- | --- |
| g4 | g4_hit_recombination | 1540 | 1.4 +/- 2.1 | 0.09 +/- 0.13 |
| g4 | g4_mbo_dnn | 1356 | 29.1 +/- 6.9 | 2.15 +/- 0.51 |
| g4 | g4_mbo_seeds | 27 | 0.5 +/- 1.2 | 1.73 +/- 4.61 |
| g4 | g4_quad_synonyms | 1 | 1.0 +/- 0.0 | 100.00 +/- 0.00 |
| g4 | g4_sample_and_screen_dnn | 1298 | 1.9 +/- 1.6 | 0.14 +/- 0.12 |

**Supplementary Table S3:** Overall library sizes for ML and HR campaigns. The third column reports the total number of variants considered so far by the campaign. For ML, this is the size of the model training set that aggregates data from all prior generations. HR recombined hits from the previous round of experiments and did not explicitly use data from prior rounds. However, for the sake of consistency we report the cumulative number of unique variants considered across rounds for both methods. Note that the third column is not simply the cumulative sum of the fourth column, since additional data was available for variants that arose from DNA synthesis and replication errors (referred to as “unmatched” in Supplementary Table S1). See ‘Hit recombination’ in Methods for details regarding how each round’s data was used by HR. Our full genotype-phenotype landscape includes additional variants, for example, positive and negative controls.

| Generation | Type of Sequence | Total number of variants used as input to design the library | Number of designed variants that data was obtained for |
| --- | --- | --- | --- |
| G1 | - | - | 9441 |
| G2 | HR | 9441 | 951 |
| G3 | HR | 10600 | 1712 |
| G4 | HR | 13252 | 1540 |
| G2 | ML | 9441 | 9370 |
| G3 | ML | 24173 | 9557 |
| G4 | ML | 43284 | 8588 |

**Supplementary Table S4:** Hyper-parameter grids for supervised models. Each model was tuned over a subset of hyper-parameter values that were established as reasonable ranges using prior investigations on other protein fitness datasets.

| Parameter Name | Non-Default Values |
| --- | --- |
| **Sklearn logistic regression** |  |
| C | logspace(1e-4, 1e4, 10) |
| 'max_iter' | 1000 |
| **LGBM classifier** |  |
| learning_rate | [0.01, 0.05, 0.1, 0.2, 0.4, 0.6] |
| n_estimators | [64, 100, 128] |
| **CNN** |  |
| batch_size | 256 |
| epochs | [2000, 3000] |
| learning_rate | [0.001, 0.00005] |
| dropout_rate | [0.05], |
| num_conv_layers | [0, 1, 2] |

**Supplementary Table S5:** Results for the best hyper-parameter setting for supervised models. We trained 3 models: linear logistic regression from <u>sklearn</u> ^122^, a gradient-boosted decision tree implemented using the LGBM <u>package</u> ^123^, and a convolutional neural network implemented in keras with tensorflow ^124^ using an ADAM optimizer with default hyper-parameters.

| model | hits@1000_wt | hits@1000_A73R |
| --- | --- | --- |
| Linear | 0.44 | 0.09 |
| LGBM | 0.71 | 0.15 |
| CNN | 0.82 | 0.21 |

**Supplementary Table S6:** Sequencing depth and NGS statistics for each library of variants.

| Generation | Library ID | Sort Percentile | Raw Reads | Merged Reads | Aligned Reads<br>(not indels) | Indels |
| --- | --- | --- | --- | --- | --- | --- |
| G1 | OB133-3-1 | N/A | 957949 | 627457 | 626924 | 58419 |
| G1 | OB133-3-2 | N/A | 958026 | 620219 | 619680 | 64609 |
| G1 | OB133-3-3 | N/A | 1500000 | 965560 | 964742 | 98036 |
| G1 | OB141 | Low Gate 1 | 992370 | 568382 | 567963 | 31143 |
| G1 | OB142 | High Gate 1 | 1064074 | 684620 | 684137 | 25680 |
| G1 | OB145 | High Gate 2 | 715124 | 430440 | 430153 | 18339 |
| G1 | OB146 | Low Gate 2 | 1012146 | 602674 | 602262 | 29599 |
| G1 | OB149 | High Gate 3 | 2264079 | 1605909 | 1605907 | 58407 |
| G1 | OB150 | Low Gate 3 | 1479973 | 1012855 | 1012239 | 37917 |
| G1 reseq | OB133-3-1 | N/A | 8796092 | 5021977 | 5018574 | 4640424 |
| G1 reseq | OB133-3-2 | N/A | 6652302 | 4288809 | 4285469 | 3927337 |
| G1 reseq | OB145 | High Gate 2 | 5042739 | 2960423 | 2958639 | 2843013 |
| ML G2 | OB188 NS 1 | N/A | 4405113 | 3813180 | 2385507 | 220066 |
| ML G2 | OB188 NS 2 | N/A | 2955910 | 2542995 | 2507476 | 75533 |
| ML G2 | OB193 | 86 | 1603156 | 1388681 | 1369732 | 45585 |
| ML G2 | OB194 | 97.5 | 1569513 | 1365392 | 1348237 | 43699 |
| ML G2 | OB196 | 93 | 4302646 | 3772972 | 2540279 | 239685 |
| ML G2 | OB188 | N/A | 14472539 | 13493968 | 8739074 | 1102314 |
| ML G3 | OB222-002 | N/A | 2154847 | 1881694 | 1552418 | 384799 |
| ML G3 | OB222-002 | N/A | 1861556 | 1605073 | 1321291 | 333557 |
| ML G3 | OB222-002 | N/A | 2224181 | 1948395 | 1621873 | 389077 |
| ML G3 | OB222-002 | N/A | 2126053 | 1904016 | 1573763 | 388317 |
| ML G3 | OB222-002 | N/A | 3393046 | 2659408 | 2165138 | 558286 |
| ML G3 | OB222-002 | N/A | 2883162 | 2327413 | 1912344 | 482233 |
| ML G3 | OB230 | 80 | 1379986 | 1199077 | 1194422 | 57309 |
| ML G3 | OB231 | 59 | 2408748 | 2120119 | 2098794 | 117612 |
| ML G3 | OB239 | 99 | 2626136 | 2065878 | 2057901 | 126476 |
| ML G3 | OB241 | 93 | 2372781 | 1879010 | 1873513 | 90589 |
| **ML G4** | **OB268** | N/A | 11923629 | 11233345 | 5248728 | 2462416 |
| **ML G4** | **OB269** | 70 | 1902601 | 1799002 | 1668940 | 87817 |
| **ML G4** | **OB270** | 90 | 2719025 | 2572901 | 2439231 | 99799 |
| **ML G4** | **OB271** | 98 | 2455165 | 2338449 | 2225624 | 87312 |
| **ML G4** | **OB272** | 99.5 | 1736640 | 1644487 | 1533991 | 80748 |

**Supplementary Table S7:**
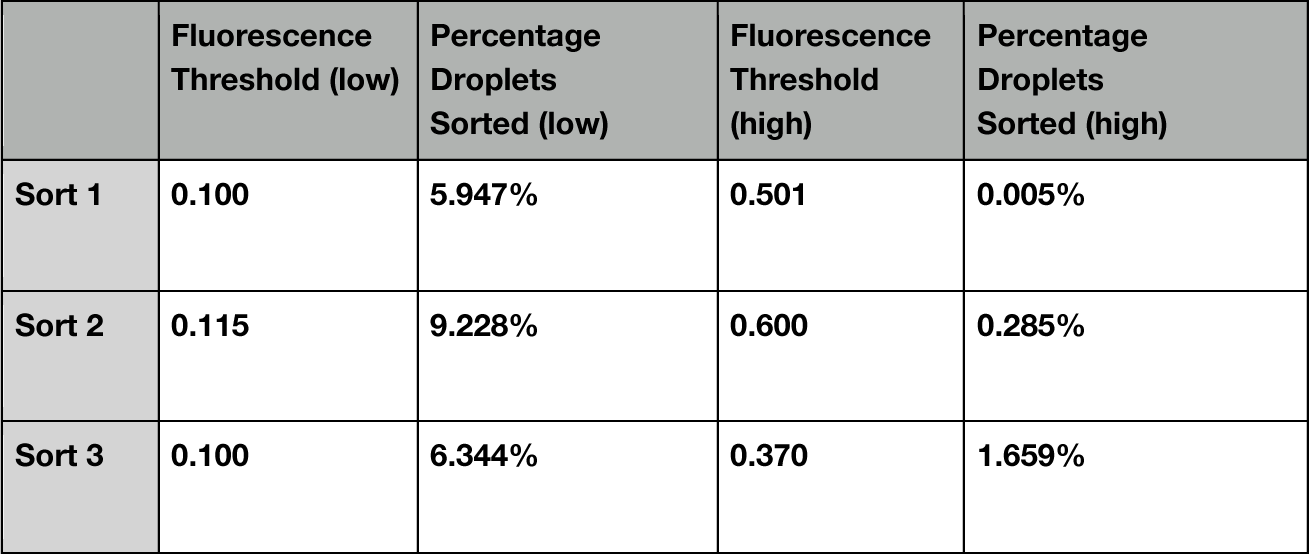
G1 sorting gate statistics. In G1 each sort was run with a low gate followed by continued sorting at a high gate. “Percentage droplets” indicates the percentage of droplets that passed the gate, where lower percentages indicate more stringent sorts at higher fluorescence threshold.

|  | Fluorescence Threshold (low) | Percentage Droplets Sorted (low) | Fluorescence Threshold (high) | Percentage Droplets Sorted (high) |
| --- | --- | --- | --- | --- |
| Sort 1 | 0.100 | 5.947% | 0.501 | 0.005% |
| Sort 2 | 0.115 | 9.228% | 0.600 | 0.285% |
| Sort 3 | 0.100 | 6.344% | 0.370 | 1.659% |

**Supplementary Table S8:** sorting statistics for gates in G2, G3, and G4. “Percentile cells” indicates the percentile of the cell-containing library we targeted for sorting. “Percentage droplets” indicates the percentage of droplets that passed the gate, where lower percentages indicate more stringent sorts at higher fluorescence threshold.

|  | Percentile cells (target) | Fluorescence threshold | Percentage Droplets Sorted |
| --- | --- | --- | --- |
| G2 |  |  |  |
| Sort 1 | 96% | 0.201 | 0.32% |
| Sort 2 | 93% | 0.318 | 0.83% |
| Sort 3 | 97.50% | 0.385 | 1.95% |
| G3 |  |  |  |
| Sort 1 | 59% | 0.118 | 5.73% |
| Sort 2 | 80% | 0.217 | 2.63% |
| Sort 3 | 95% | 0.285 | 0.65% |
| Sort 4 | 99% | 0.346 | 0.12% |
| G4 |  |  |  |
| Sort 1 | 70% | 0.124 | 4.25% |
| Sort 2 | 90% | 0.199 | 1.51% |
| Sort 3 | 98% | 0.306 | 0.26% |
| Sort 4 | 99.50% | 0.384 | 0.17% |

## Supplementary Text

### S1: Composition of the ML library in each round

#### MBO-DNN

##### G2

*model:* We used a CNN regression model with 3 convolutional layers containing 32 width-5 filters and two fully-connected layers of width 64. The input to the model is a shape = [sequence_length, 20] one-hot-encoded array. After selecting a model architecture and training optimizer hyper-parameters using cross validation, we retrained the model on all available data using 5 replicates with different random parameter initialization. The model score used by MBO is the average prediction of this ensemble. Unlike other rounds, we fit models on a continuous-valued transformation of enrichment factors (Supplementary Text S7), though in hindsight we feel that the discrete labeling used in subsequent rounds was more reliable.

*candidate generation:* First, we drew 500M samples from the ‘broad recombination sampling’ distribution.

mbo-dnn-exploit: take all samples with score > 3.

mbo-dnn-explore: take all samples with score > 0.

*batch selection:* same as G4; with restrictions on mutation usage and a specified target distribution over number of mutations from WT.

Note that the ML2 variant A73R,D74S in Figure 3b was proposed by mbo-dnn-explore.

##### G3

*model:* We used a similar approach as in G2, except that the model had 3 convolutional layers of 32 width-5 filters and a single fully-connected layer of width 64. The model was fit on discrete activity labels.

*candidate generation:* We recombine candidates from two approaches: (1) the same local-search technique as used in G4 and (2) samples from the G3 Prosar methods.

*batch selection:*

mbo-dnn-exploit: the top-200 candidates, ranked by model score.

mbo-dnn-explore: same as G4; with restrictions on mutation usage and a specified target distribution over number of mutations from WT.

##### G4

Described in the Methods section.

#### Prosar

In G3 we use ‘prosar_high’ and ‘prosar_low’ to denote the use of mutation effect score thresholds of 2.0 and 0.5.

#### Prosar+Screen

We use ‘prosar_x_screen_y’ to denote filtering samples from prosar_x (see above) using either a high or low VAE score threshold y. These thresholds were set heuristically by examining the distribution of VAE scores for samples from prosar_x. The VAE likelihood of a given variant is estimated using importance sampling from the posterior with 2500 samples.

#### Prosar+Screen “G2 redux”

Prosar+Screen worked well in G3 and so we assessed how it would have performed in G2, where only the G1 epPCR was available. The VAE score threshold was set heuristically to the 90th percentile of scores for sampled recombination variants with 4 mutations, which resembles the high-threshold filtering used in G3. We heuristically defined the mutation set as the top 45 mutations, since this was about the size of the mutation set used in the best prosar+screen system from G3.

#### sample+screen-linear

This is an ablation of the G4 sample-and-screen method described in the Methods section where we use a linear model instead of a neural network to classify a variant’s activity level.

#### mbo-linear

This is an ablation of the G4 mbo-dnn where we used a linear multi-class logistic regression model.

#### G4 and G3 hit constituents

For a given multi-mutation variant X, the set of ‘constituents’ of X is all possible variants that combine subsets of the mutations in X. In G4 we include the constituents of a few top-performing variants from the G3 liquid culture plate assay.

### S2: Discussion of hit-rates for additional sub-libraries

#### G2 sublibrary hit-rates

Broad recombination sampling (BRS) sampled combinations of mutations from the G1 data that were inferred to be non-deleterious to protein function. However, as seen in Supplementary Figure S1, BRS performed quite poorly. In hindsight, we were too permissive in defining this set of non-deleterious mutations (see ‘broad recombination sampling’ in Methods). Since we sample variants with many mutations, many variants probably contained at least one false-positive mutation. MBO-DNN-Explore screened samples from BRS using a neural network model. This is marginally better than BRS, but was still hurt by the overall low quality of the pool of samples. MBO-DNN-Exploit performed comparably to hit recombination (HR). Had BRS been tuned to perform better, it may have led to downstream improvements in the overall campaign.

#### G3 sublibrary hit-rates

The purified protein results shown in Figure 3 confirm that G3 contained at least one variant with activity much better than A73R. This is also confirmed by our hit calling procedure based on NGS counts, where we see in Supplementary Table S1 that many G3 sublibraries proposed variants that were labeled as having activity higher than A73R. For example, MBO-DNN-Exploit found 11 such variants. However, the hit rates for finding such variants are sufficiently low that we cannot stratify the analysis by num_mutations.

In response, we analyzed methods’ hit rate for designing variants with activity greater than WT. We consider the HR campaign and the key versions of the two types of ML-based design approaches in our G3 portfolio: MBO-DNN and Prosar+Screen. We find that these outperformed hit recombination (HR). However, there are a number of hyper-parameters for each that impact performance and may be difficult to set in advance. In Supplementary Figure S2, for example, we see that both MBO-DNN systems performed similarly, but the left panel of Supplementary Figure S3 demonstrates that Prosar+Screen was sensitive to the parameters we used to trade off exploration and exploitation.

#### Prosar+Screen Ablations

In Supplementary Figure S3 (left) we analyze the G3 sub-libraries designed using the Prosar+Screen method with different design choices for trading off exploration and exploitation, including ‘Prosar’ which performs no screening with a zero-shot model. See Prosar+Screen in Section S1 for the prosar_x_screen_y notation. First, we find that all Prosar+Screen versions outperformed Prosar for designing variants with many mutations. Second, the exploit-focused versions of the method had the highest hit rates.

Drawing on the success of Prosar+Screen in G3 and the small number of active variants in the G2 library due to the BRS sampling (see above), we investigated how Prosar+Screen would have performed in G2 as an alternative library design approach (see ‘g2 redux prosar+screen’ above). We find in Supplementary Figure S3 (right) that it was significantly better than hit recombination in G2 (HR-G2) at designing variants with many mutations and is an interesting competitor to HR-G3.

#### Sample+Screen Ablations

In G4, we ran an investigation to analyze the impact of our classification model for screening variants. We ran two versions of Sample+Screen, where a large pool of candidate variants is sampled from a VAE that was trained on a combination of homologs and G1-G3 hits and then filtered using a classification model. Sampling is done using the same technique as in ‘Zero-shot neighbor sampling’. In Supplementary Figure S4 (left) we consider methods’ ability to find variants with activity higher than the WT and find that filtering using our convolutional neural network (Sample+Screen-DNN) is significantly better than filtering using a linear model (Sample+Screen-Linear), which performs about the same as using raw samples with no filtering (Sample-Unscreened). We also contrast Sample+Screen-DNN with MBO-DNN, which uses local search to find high-scoring candidates instead of ranking a fixed pool of samples and find that MBO-DNN performs better. In Supplementary Figure S4 (right) we consider the methods’ ability to find variants with activity higher than A73R. Here, MBO-DNN is dramatically better than Sample+Screen. We expect this is because including the homology data when training the sampling model biases the samples away from the known experimental hits towards the variability of the natural homologs. For MBO-DNN, however, it optimizes a model-based fitness landscape based entirely on experimental data.

### S3: Zero-shot results

#### A: Benchmarking Zero-Shot Scoring

Our zero-shot VAE model provides two capabilities: scoring the likelihood of a given variant and sampling a novel variant. There are many other computational methods that also provide these, ranging, for example, from context-independent models of amino acid substitution probabilities to deep neural networks that define a full joint distribution over variable-length protein sequences. Here, we evaluate a collection of such models in terms of their ability to perform zero-shot scoring on the G1 epPCR data.

In Supplementary Figure S5 and Supplementary Figure S6 we provide the same figures as Figure 6a and b, but using scores from 4 different approaches: our VAE, the 650M-parameter ESM-1b model ^70^, a profile hidden Markov model (pHMM) fit using the hmmer package ^71^ on the same alignment used to train the VAE, and context-independent scoring using substitution scores from the BLOSUM62 matrix ^125^. In Supplementary Figure S5 we show qualitatively that the VAE scores provide the most noticeable separation between functional and non-functional variants.

In Supplementary Figure S6 we analyze methods’ ability to filter a library to a smaller size while still keeping the hits. The rows of the grid of sub-figures correspond to which level of activity is used to define a hit and columns correspond to which set of variants we analyze. In the top row, we analyze models’ ability to identify variants with non-zero activity. This is a setting where we expect zero-shot scoring to perform well, since mutations that render a protein non-functional may be easy to infer using patterns of conservation in natural sequences. In the bottom row, we explore models’ ability to identify variants with activity that is greater than WT, which is a primary protein optimization goal. In the left column, analyzing variants with num_mutations = 1 reflects a use-case constrained by low-throughput experiments where assaying even all single-mutant variants of a starting sequence would be infeasible, and thus it would be helpful to filter this set *in-silico*. On the right, we consider all G1 variants with multiple mutations, which probes models’ ability to identify epistatic interactions. One challenge with the right figure is that enzyme activity is correlated with the number of mutations from WT. To control for this, we only consider variants with exactly two mutations in the middle column. Overall, we find that the VAE performs comparable or better than the alternative methods across this range of evaluation settings.

#### B: Comparing Zero-Shot Sampling to Homolog Grafting

‘Homolog grafting’ is a model-free zero-shot design approach that incorporates groups of mutations appearing in nearby homologs to the WT (Methods). Of the 11 homolog grafting variants, 5 ± 2 were functional (Supplementary Table S1), similar to the overall hit rate of the VAE-designed zero-shot library (hit rates for functional variants of 50.9 ± 16.8% vs 50.9 ± 1.8%, respectively). While this model-free approach showed a promising hit-rate, natural homologs are often quite far from the wildtype sequence, so generating a large library within a certain mutational distance of the WT is not straightforward, as grafting from a distant homolog would result in many mutations. VAE sampling, on the other hand, trains a model on distantly-related homologs, but can sample many new nearby neighbors of the WT.

### S4: G4 MBO-DNN Model Selection

We assess model performance using the following steps:

1. Create a train-test split of the G1, G2, and G3 data by using all variants with less than or equal to two mutations for training and the remainder for testing.
2. Give each variant two binary labels using the procedure described in Section S7: whether it has activity higher than the WT and whether it has activity higher than A73R. These were then merged into a single multi-class label. Of note is that we do not perform any EFDR correction when producing these labels; each variant is processed independently.
3. Train models using a multi-class classification loss.
4. For each model and for each activity level, define a sequence-to-scalar ranking function as score(sequence; activity_level) = P(sequence | activity_label >= activity_level). For example, score(sequence; WT) = P(sequence | label = WT) + P(sequence | label = A73R).
5. compute hits@1000_wt, the fraction of the 1000 variants with highest score(sequence; WT) that have been labeled as better-than-WT. And similarly, compute the hits@1000_a73R metric, where variants are ranked using score(sequence; A73R).
6. To optimize over hyper-parameters for each model, we sorted by the hits@1000_A73R metric, as we felt this best reflected our goal of finding new, improved variants. See Supplementary Table S4 and **Supplementary Table S5** for hyperparameters and model performance metrics.

### S6: Additional details about assays of enzyme activity

#### 6A Isolating the highest activity variants using a liquid culture plate assay

After stringently sorting at successively higher thresholds, cells that pass the selection were plated and single colonies screened in a “Tier 1” liquid culture plate assay. In Tier 1, each colony was assayed (Methods: Liquid Culture Plate Assay) in a single well on a 96-well plate that included controls of high activity variants selected in prior rounds. The ranking of activity (normalized by optical density) in Tier 1 was used to identify high performing variants that were genotyped and promoted to “Tier 2”. In Tier 2, each variant was assayed at higher replication to confirm their relative ranking. High performing variants in the Tier 2 screen were promoted for protein purification and assayed for specific activity. Results for Tier 1 and 2 screens for all generations are presented in Supplementary Figure S8 - Supplementary Figure S13.

#### 6B Specific activity assay

To determine the specific activity of a purified variant we compare the reaction kinetic curves between the variant of interest and the wildtype at a series of enzyme concentrations, as described in Methods: Nuclease activity assays. In Supplementary Figure S14 we present an example kinetic curve and describe how we compute specific activity.

#### 6C Dependence of wildtype nuclease activity on pH

To determine how the wildtype activity differs under different pH conditions, we conducted enzyme activity assays using several buffers to cover a pH range from 6 to 11, see Supplementary Figure S16. These assays consisted of 25 mM buffer, 0.1% pluronic, 2.5 mM MnSO4, 50 uM G25/G24, 0.2 uM G651/G537, while the enzyme buffer also contained 25 mM buffer with 0.1% pluronic. Rates were determined from the linear region of the kinetic curve. corresponding to 30-750s. Note that most assays are conducted with 100mM buffer concentrations to compensate for the alkalinity in spent media, but in this assay only 25mM was used because the enzyme was diluted in matched pH buffer before the assay. In our kinetic assays, the MOPS buffer was used.

Our protein engineering campaign was conducted at pH 7, and we report relative activity everywhere with respect to the wildtype enzyme at pH 7 unless otherwise specified. A 5.9-fold improvement relative to the wildtype is necessary to restore the enzyme to its maximum activity (i.e. its activity at pH 9), an 11.8-fold improvement in our campaign would imply an enzyme with 200% activity of the maximum at pH 9, etc.

### S7: Generating ML Training Labels

In the Methods section, we describe a procedure for calling ‘hits’ with respect to a fiducial sequence, i.e., a set of variants with activity that is likely above that of a fiducial sequence. This approach was used when reporting all results about hit rates and to create the final dataset that we have released. Note, however, that we also needed to generate a dataset of sequences and corresponding activity labels during each round of the campaign for training the machine learning models that would design the next round, and our approach for creating this differed from the above procedure in a few key ways.

First, we did not perform any false-discovery rate correction; each variant was compared to fiducial sequences independently. This increases the false-positive rate of our labeling, which we felt was desirable to increase ‘optimism in the face of uncertainty’, a general design principle for trading off exploration and exploitation in model-based optimization ^126^. Second, we performed both right and left-sided tests with respect to fiducials. This was used to provide fine-grained labels, such as ‘WT+’, where the activity is labeled as better than WT, but worse than A73R. Third, some variants appeared in multiple rounds. To reconcile potentially conflicting labels across rounds, we took the label from the most recent round.

Our labeling approach for G2 and G3 (i.e. the training data used to design G3 and G4) can be represented as a decision tree, which we established heuristically by reasoning about which gates had resolution for labeling activity as higher or lower than which fiducials. We visualize these decision trees in Supplementary Figure S26. Let, for example, EF_A73R_^sort^ (1-α) refer to the (1-α) quantile of the enrichment factor distribution of the synonyms of fiducial A73R in a particular sort. If a variant has an enrichment factor greater than EF_A73R_^sort^ (1-α) quantile of a fiducial, we label the variant as being “better” than A73R. If it has an enrichment factor greater than the (α) quantile, we label the variant as being “comparable” to A73R. We do such comparisons at each node in the decision tree, using different fiducials and different gates.

Finally, to design generation G2 using MBO-DNN, we used continuous regression targets for each G1 variant instead of discrete activity labels. In hindsight, these were more noisy than we thought, and we would use discrete labels if we were re-designing G2 again. This can be seen in our ‘g2 redux’ investigation, which was based on discrete labels. There were 3 high threshold sorts and 3 low threshold sorts, each with negative control and WT fiducials. We computed the mean μ and standard deviation σ of the WT fiducial distribution (trimmed at the top/bottom 1% to remove outliers). Then, for each variant we computed the z-score of its enrichment factor z_V_^sort^ = EF_V_^sort^ - μ_WT_^sort^ / σ_WT_^sort^. To combine the information across all sorts, for each variant we assigned an aggregate score by taking the max z score across the 3 high threshold sorts, and clipping the scores to be between [-4, 4]. In addition, any variant with an enrichment factor <= 1 in all low threshold sorts was mapped to score = -4.

### S8: Gate Selection

For each sort, we had to select fluorescence thresholds for sorting cells to enable us to characterize variants at different levels of activity. Selecting at higher fluorescence thresholds selects for higher activity. With the exception of the first generation, we selected gates based on the fluorescence distribution of fiducial sequences; activity relative to fiducials was then used to characterize our high-throughput libraries. Our procedure for determining our gates was as follows. We first performed a fluorescence read experiment with fiducial sequences together with the library to determine the fluorescence distribution of our fiducials and our library. For the activity levels we were targeting (i.e. “greater than A73R”), we picked the appropriate fluorescence thresholds and computed the corresponding percentile of the overall library. For the sorting experiment, we also count the percentage of droplets that pass the gate.

#### G1 Gating design

In G1, tested variants were drawn from a randomly generated library (epPCR) on top of a single-site saturation mutagenesis (SSM). Low gate cutoffs were chosen to differentiate between very poor performers (“dead”) and active variants (“alive”). High gate cutoffs were chosen to differentiate between top performers and the rest of the library. Different from all the other libraries, no fiducial sequences were used to determine the gate value of this library.

#### G2 Gating Design

In G2, we focused on differentiating between variants that were comparable to, or better, than the parent. G2 gates were set at the 86th, 93rd, and 97.5th percentile of the library distribution, corresponding to the left tail, mean, and right tail of the parent fiducial fluorescence distribution, respectively.

#### G3 Gating Design

In G3, we focused on differentiating between variants that were better/comparable to the parent and better/comparable to the best variant in G2, i.e. A73R. G3 gates were set at the 59th, 80th, 95th, and 99th percentiles of the library distribution, corresponding to the mean of the parent fiducial, an intermediate value between the parent and A73R fiducial distributions, the left tail of the A73R fiducial, and the mean of the A73R fiducial, respectively.

#### G4 Gating design

In G4, we had multiple target activities we wanted to assess, since sublibraries were focused on both ablations and improvements on previous rounds. G4 gates were set to 70th, 90th, 98th, and 99.5th percentiles, corresponding to the mean fluorescence of the parent, the best G2 variant A73R, the best G3 double (D64A, A73R), and the best G3 quadruple (A63P, A73R, D74H, I84Y), respectively (see Supplementary Figure S27d).

### S9: Additional Sublibrary Details

#### Hit-recombination

In G2, we recombined the top 20 hits from G1 that contained mutations in the G2 design region (see ‘Methods: Designed DNA library generation’). The G3 design regions contained residues not appearing in the G2 design region, so we considered the following pools for recombination: the top 30 variants appearing in (or 1 mutation from) the G2 HR library, the top 30 variants from G1 that had previously not been feasible in the G2 design region but were now possible in the expanded G3 regions, and the top 10 single-mutation variants observed in G2 that were not screened in G1. In G4, we recombined the top 70 variants designed by (or 1 aa from) the HR library in G3.

#### Designed synonyms

In G1, WT synonyms were randomly generated via epPCR. In G2 we picked 50 of the WT synonyms that had appeared in G1, where 37 had 1 nucleotide mutation and 13 had 2 nucleotide mutations. In G3 we constructed 50 WT synonyms and 50 A73R synonyms, stratified by the number of codon mutations: at each of C = 1, 2, 3, 4, 5 we selected c codons within the design region at random and replaced the codon with its highest frequency synonym. We did this 10 times for each value of C, leading to 50 synonyms total. We also included the 50 WT synonyms used in G2. In G4 we constructed 100 synonyms of WT, A73R, and A73R,D74S using a similar procedure, constructing 10 synonyms each at C = 2,3, …, 11.

